# Functional genomics of trypanotolerant and trypanosusceptible cattle infected with *Trypanosoma congolense* across multiple time points and tissues

**DOI:** 10.1101/2025.01.31.635858

**Authors:** Gillian P. McHugo, James A. Ward, John A. Browne, Grace M. O’Gorman, Kieran G. Meade, Emmeline W. Hill, Thomas J. Hall, David E. MacHugh

**Affiliations:** UCD School of Agriculture and Food Science, University College Dublin, Belfield, Dublin, Ireland; UK Agri-Tech Centre, Innovation Centre, York Science Park, York, UK; UCD Conway Institute of Biomolecular and Biomedical Research, University College Dublin, Dublin, Ireland; UCD One Health Centre, University College Dublin, Dublin, Ireland

## Abstract

Human African trypanosomiasis (HAT), or sleeping sickness, is a neglected tropical disease caused by infection with trypanosome parasites (*Trypanosoma* spp.). These are transmitted by infected tsetse flies (*Glossina* spp.) and cause a similar disease in animals, known as African animal trypanosomiasis (AAT). AAT is one of the largest constraints to livestock production in sub-Saharan Africa and causes a financial burden of approximately $4.5 billion annually. Some African *Bos taurus* cattle populations have an important evolutionary adaptation known as trypanotolerance, a genetically determined tolerance of infection by trypanosome parasites (*Trypanosoma* spp.). Trypanotolerant African *B. taurus* N’Dama and trypanosusceptible *Bos indicus* Boran cattle responded in largely similar ways during trypanosome infection when gene expression was examined using blood, liver, lymph node, and spleen samples with peaks and troughs of gene expression differences following the cyclic pattern of parasitaemia exhibited during trypanosome infection. However, differences in response to infection between the two breeds were reflected in differential expression of genes related to the immune system such as those encoding antimicrobial peptides and cytokines, including, for example, the antimicrobial peptide encoding genes *LEAP2*, *CATHL3*, *DEFB4A*, and *S100A7* and the cytokine genes *CCL20*, *CXCL11*, *CXCL13*, *CXCL16*, *CXCL17*, *IL33*, and *TNFSF13B*. In addition, transcriptional profiling of peripheral blood identified expression differences in genes relating to coagulation and iron homeostasis, which supports the hypothesis that the dual control of parasitaemia and the anaemia resulting from the innate immune response to trypanosome parasites is key to trypanotolerance and provide new insights into the molecular mechanisms underlying this phenomenon.

**Author Summary:** Trypanosome parasites are transmitted by infected tsetse flies and cause the neglected tropical disease human African trypanosomiasis (HAT) and the similar African animal trypanosomiasis (AAT), which is one of the largest impediments to livestock production in sub-Saharan Africa. Taurine (*Bos taurus*) and indicine (*Bos indicus*) cattle shared a common ancestor more than 150,000 years ago, and in the intervening period significant genomic differences have evolved between the two groups. Importantly, several African *B. taurus* populations have evolved an adaptation known as trypanotolerance, a genetically determined tolerance of infection by trypanosome parasites. Trypanotolerant African taurine and trypanosusceptible indicine cattle responded in largely similar ways during trypanosome infection when gene expression was examined using blood, liver, lymph node, and spleen samples with peaks and troughs of gene expression differences following the cyclic pattern of the number of trypanosome parasites in the blood. Transcriptional profiling of these tissues highlighted genes related to the multiple facets of the immune system; notably, for peripheral blood, differences observed for genes relating to coagulation and iron homeostasis support the existing hypothesis that control of both parasite number and anaemia is an important feature of the trypanotolerance trait and provide new insights into the molecular mechanisms underlying this phenomenon.

## Introduction

Trypanosomiasis is prevalent in humid and semi-humid regions of Africa and is a wasting disease caused by parasitic protozoa of the genus *Trypanosoma* which are transmitted by biting insect vectors such as tsetse flies (*Glossina* spp.) [1]. Human African trypanosomiasis (HAT), caused by *T. brucei gambiense* or *T. brucei rhodesiense* infection, is also known as sleeping sickness and is classed as a neglected tropical disease (NTD) by the World Health Organisation [2]. Trypanosomes also infect animals and cause particularly severe disease in domesticated animals—African animal trypanosomiasis (AAT)—for which the symptoms, as in HAT, include fever, severe weight loss, and anaemia [1, 3, 4]. As the disease progresses, animals weaken and become paralysed and unfit for work [1]. AAT is caused by a wider range of trypanosome parasites, most commonly *T. congolense, T. vivax*, and *T. brucei* ssp., although the prevalence and distribution of trypanosome species varies and co-infections are possible [5, 6]. In addition, domestic animals can also be infected with the human-infective trypanosome species, for which they act as a reservoir [5, 7].

Animal models, including mice, have long been used to study human-infective trypanosomes leading to important insights into host-trypanosome interactions [8–10]. The usefulness of these animal models has been improved by the differential host tolerance of, or susceptibility to, trypanosome infection that has been found between different cattle breeds, mouse strains, and human populations, although the underlying genes differ between host species [11–14]. Conversely, while it is much less common, humans may also be infected with trypanosome species more frequently found in animals, resulting in atypical human trypanosomiasis [5, 15]. With climate change and increasing globalisation there is growing concern that this atypical human trypanosomiasis may represent an emerging neglected zoonotic disease that underscores the importance of a One Health approach to both human and animal trypanosomiasis [16].Trypanosomiasis has a limiting effect on sub-Saharan livestock production, because even with application of trypanocidal drugs, the susceptibility of most cattle to trypanosomiasis makes production economically unsustainable in many regions on the African continent [17].

Cattle are significant components of rural economies and livelihoods in Africa. They provide milk, meat, fertiliser and traction and represent mobile assets that serve as a financial buffer for poor families, particularly for pastoralists and women [18, 19]. There are approximately 150 breeds of indigenous cattle in sub-Saharan Africa and African cattle represent a complex mosaic of African *Bos taurus*, European *B. taurus*, *B. indicus*, and various hybrid populations [20–22]. African cattle form a gradient of *B. taurus* and *B. indicus* ancestry across the continent [20]. This is a result of multiple domestications and the subsequent spread of distinct populations of cattle across Africa [23]. The spread of agriculture and livestock herding has also shaped African human genetics and linguistic variation, illustrating the importance of domesticated species in influencing recent human evolution [24, 25]. Zebu or indicine breeds, which have primarily *B. indicus* ancestry, are favoured by many farmers due to their larger size and higher production yields [26]. However, cattle populations in West Africa tend to have higher level of taurine (*B. taurus*) ancestry [20, 27]. This is because some indigenous *B. taurus* breeds have an advantage in western sub-Saharan Africa due to their tolerance of trypanosomes, a trait termed “trypanotolerance” [26, 28]. These cattle exhibit a greater ability to control parasitaemia and anaemia, making them more productive than *B. indicus* cattle or other *B. taurus* breeds in areas infested with tsetse flies and trypanosomes [26, 29]. These trypanotolerant breeds, which include the longhorn N’Dama and shorthorn Baoule, Lagune, and Somba breeds, are therefore an important genetic resource as they are uniquely suited to livestock production in these areas [30, 31]. Trypanotolerant breeds only make up 6% of the total cattle population of Africa and but represent 17% of the cattle in the tsetse infested areas [32]. Although trypanotolerance is a valuable trait in these areas, trypanotolerant breeds such as the N’Dama, are not more widely used because of their relatively inferior production characteristics, unpredictable temperament and smaller size, which make them unsuitable for draft purposes [32, 33].

Trypanotolerance has been shown to be a heritable trait, although there is variability in tolerance between animals within breeds [13, 29, 34, 35]. Trypanotolerant breeds are also less susceptible to other infectious diseases such as helminthiasis, ticks and tick-borne-diseases and genes associated with the immune response have been found to be under selection in west African cattle populations [36, 37]. Trypanotolerant and trypanosusceptible breeds have been shown to vary in their immune responses and exhibit gene expression differences in various tissues at key points during infection after infection with trypanosomes [33, 38–42]. They also show differences in ability to control anaemia over two years [43]. There are also *B. taurus*/*B. indicus* hybrid breeds that have been observed to be trypanotolerant [44]. However, trypanotolerant breeds with high levels of African *B. taurus* ancestry are better able to control anaemia while hybrid animals exhibit intermediate levels of control when compared to susceptible *B. indicus* breeds [45]. The genes, genomic regulatory elements (GREs), and genomic regulatory networks (GRNs) underpinning trypanotolerance remain largely unknown, although several candidate genes and regulatory mechanisms have been suggested based on population genomics and functional genomics studies [33, 40, 42, 43, 46–49]. It has been proposed that a better understanding of the trypanotolerance trait will facilitate genome-informed breeding programmes that could also leverage transgenesis to enhance the trait, thereby increasing the productivity of livestock production in sub-Saharan Africa [26, 33].

In this study we analysed Affymetrix^®^ Bovine Genome Array gene expression data across multiple tissues from trypanotolerant N’Dama and trypanosusceptible Boran cattle that had been infected with *T. congolense* across an infection time course [33, 40, 41]. We used previously published data from liver, lymph node, and spleen tissue samples and combined these data with new peripheral blood mononuclear cell (PBMC) data from the same animals. Differentially expressed genes (DEGs) were detected across the infection time course for each breed and between the two breeds for each time point. The sets of DEGs were then used for functional enrichment analyses focused on gene ontology (GO) to catalogue and interrogate biological processes associated with disease pathogenesis and trypanotolerance.

## Materials and methods

### Data sources

#### New microarray gene expression data

Affymetrix^®^ Bovine Genome Array data sets were generated for 40 peripheral blood mononuclear cell (PBMC) samples from 10 African cattle (5 trypanotolerant N’Dama and 5 trypanosusceptible Boran) collected at four time points for a *T. congolense* infection experiment performed in 2003 [39–41]. The cattle, all female and aged between 19–28 months, were reared together at the International Livestock Research Institute (ILRI) ranch at Kapiti Plains Estate, Kenya which is located in an area free from tsetse flies and trypanosomiasis [41]. The cattle were experimentally infected with the *T. congolense* clone IL1180 [50, 51] delivered via the bites of eight infected tsetse flies (*Glossina morsitans morsitans*) [52, 53] at the ILRI laboratories in Nairobi, Kenya. The flies were allowed to feed on the shaved flanks of the animals until engorgement. The infections were carried out in three batches which took place two weeks apart and were confirmed by microscopy. Parasitaemia and anaemia measured by packed cell volume (PCV) were monitored throughout the course of the infection [41].

Peripheral blood (200 ml) was collected in heparinised syringes before infection and at 14, 25, and 34 days post-infection (DPI), these time points were selected to align with the first wave of parasitaemia [41]. This animal work was completed prior to the requirement for formal Institutional Permission in Ireland, which is based on European Union Directive 2010/63/EU; however, all animal procedures were approved by the Institutional Animal Use and Care (IAUC) Committee of ILRI in accordance with the UK Animals (Scientific Procedures) Act 1986 and all efforts were made to ensure high welfare standards and ethical handling of animal subjects.

Peripheral blood mononuclear cell (PBMC) samples were isolated from the blood samples with Percoll™ gradients (GE Healthcare, Buckinghamshire, UK) and total RNA was extracted from the PBMCs in TriReagent (Molecular Research Center, Inc., Cincinnati, OH, USA) and the RNA samples were then DNase-treated and purified using an RNeasy mini kit (Qiagen Ltd., Crawley, UK). RNA quality and quantity was assessed using the 18S/28S ratio and RNA integrity number (RIN) on an Agilent Bioanalyzer with the RNA 6000 Nano LabChip^®^ kit (Agilent Technologies, Inc., Santa Clara, CA, USA). Following this, cDNA labelling, hybridisation and scanning for the microarray experiments were performed by Almac Diagnostic Services (Craigavon, Northern Ireland) using a one-cycle amplification/labelling protocol. Gene expression data in the form of cell intensity files (.CEL) were generated using the Affymetrix GeneChip^®^ Operating Software (GCOS) package. The Affymetrix^®^ GeneChip^®^ Bovine Genome Array data sets generated for this study have been deposited in the European Molecular Biology Laboratory European Bioinformatics Institute ArrayExpress data repository under accession number E-MTAB-14517 (temporary manuscript reviewer link: https://www.ebi.ac.uk/biostudies/arrayexpress/studies/E-MTAB-14517?key=b9fd37fb-644d-442b-b443-53506a43a8df).

#### Previously published microarray gene expression data

Additional Affymetrix^®^ Bovine Genome Array data sets were obtained from a published study using solid tissues samples from the same animal infection time course [33] resulting in a total of 220 biological samples across 12 infection time points and four tissues (for both N’Dama and Boran) before filtering (**Table 1**). **Fig 1** illustrates the experimental design and study workflow. The computer code required to repeat and reproduce the analyses is available at doi.org/10.5281/zenodo.11502109.

**Fig 1.**
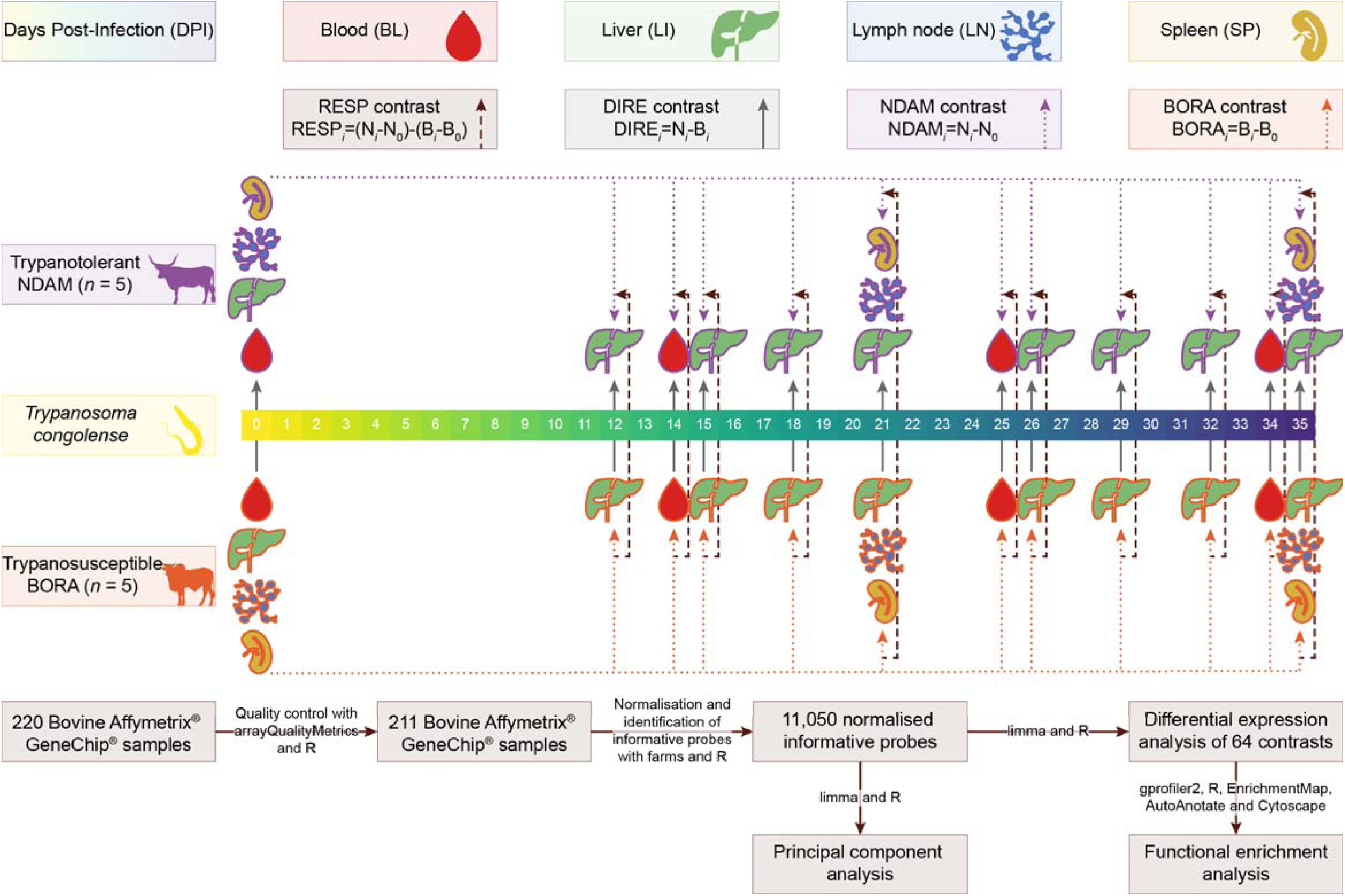
Diagram showing the experimental design and study workflow. Trypanosome image by Matus Valach and cattle images by Tracy A. Heath and T. Michael Keesey via phylopic.org and tissue images via healthicons.org. Colours were generated from the khroma (v. 1.10.0) [54] and viridis (v. 0.6.3) [55] R packages.

**Table 1.**
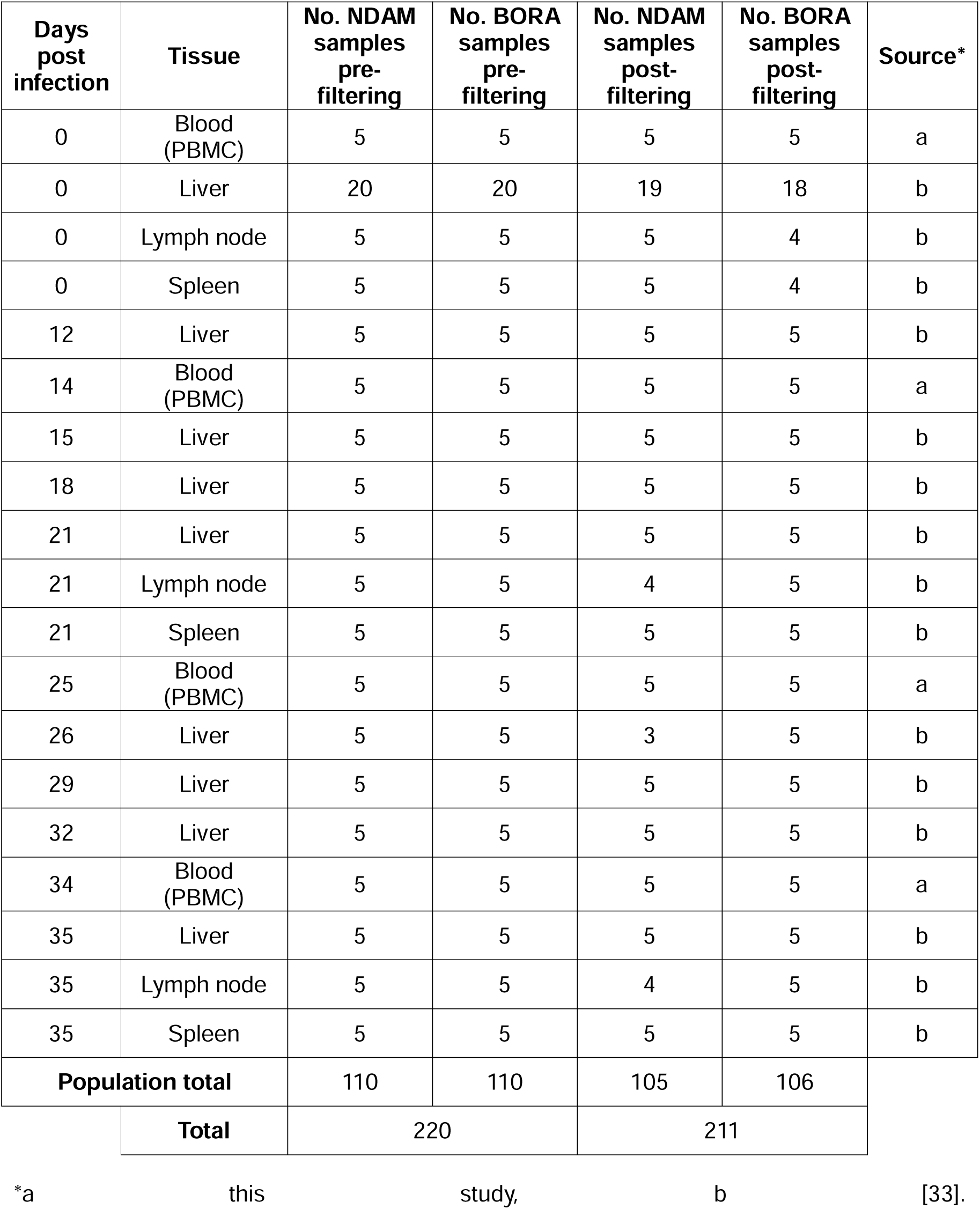
Days post infection, tissues, numbers of N’Dama (NDAM) and Boran (BORA) samples, and sources of microarray data used in this study before and after sample filtering.

| Days post infection | Tissue | No. NDAM samples pre-filtering | No. BORA samples pre-filtering | No. NDAM samples post-filtering | No. BORA samples post-filtering | Source* |
| --- | --- | --- | --- | --- | --- | --- |
| 0 | Blood (PBMC) | 5 | 5 | 5 | 5 | a |
| 0 | Liver | 20 | 20 | 19 | 18 | b |
| 0 | Lymph node | 5 | 5 | 5 | 4 | b |
| 0 | Spleen | 5 | 5 | 5 | 4 | b |
| 12 | Liver | 5 | 5 | 5 | 5 | b |
| 14 | Blood (PBMC) | 5 | 5 | 5 | 5 | a |
| 15 | Liver | 5 | 5 | 5 | 5 | b |
| 18 | Liver | 5 | 5 | 5 | 5 | b |
| 21 | Liver | 5 | 5 | 5 | 5 | b |
| 21 | Lymph node | 5 | 5 | 4 | 5 | b |
| 21 | Spleen | 5 | 5 | 5 | 5 | b |
| 25 | Blood (PBMC) | 5 | 5 | 5 | 5 | a |
| 26 | Liver | 5 | 5 | 3 | 5 | b |
| 29 | Liver | 5 | 5 | 5 | 5 | b |
| 32 | Liver | 5 | 5 | 5 | 5 | b |
| 34 | Blood (PBMC) | 5 | 5 | 5 | 5 | a |
| 35 | Liver | 5 | 5 | 5 | 5 | b |
| 35 | Lymph node | 5 | 5 | 4 | 5 | b |
| 35 | Spleen | 5 | 5 | 5 | 5 | b |
| **Population total** |  | 110 | 110 | 105 | 106 |  |
| **Total** |  | 220 |  | 211 |  |  |

\*a this study, b [33].

### Data filtering and normalisation

Quality control with log transformation was performed using affy (v. 1.80.0) [56], Biobase (v. 2.62.0) [57], and arrayQualityMetrics (v. 3.58.0) [58] with R (v. 4.3.2) [59]. As in previous microarray studies, any individual samples that were identified as outliers in two or more quality control tests were discarded from the analysis [60]. Normalisation was performed using farms (v. 1.25.0) [61] with R (v. 4.3.2).

The intensity of expression for each sample was visualised both with the raw data and after normalisation using affy (v. 1.80.0), Biobase (v. 2.62.0), dplyr (v. 1.1.2) [62], ggh4x (v. 0.2.4) [63], ggplot2 (v. 3.4.2) [64], ggtext (v. 0.1.2) [65], magrittr (v. 2.0.3) [66], readr (v. 2.1.4) [67], reshape2 (v. 1.4.4) [68], and stringr (v. 1.5.0) [69] with R (v. 4.3.2). Colours were generated from khroma (v. 1.10.0) [54] and viridis (v. 0.6.3) [55]. Informative probe sets were identified and extracted using farms (v. 1.25.0) with R (v. 4.3.2).

### Principal component analysis

Principal component analysis (PCA) was performed using limma (v. 3.58.1) [70] with R (v. 4.3.2). The results were visualised using dplyr (v. 1.1.2), ggplot2 (v. 3.4.2), patchwork (v. 1.2.0) [71], and stringr (v. 1.5.0), with R (v. 4.3.2). Colours were generated from khroma (v. 1.10.0) and viridis (v. 0.6.3).

### Differential expression analysis

Differential expression analysis was performed using limma (v. 3.58.1) [70] with R (v. 4.3.2). The correlation between samples from the same animal was estimated and included in the linear model [70, 72]. The contrast matrix contained 64 contrasts across the two populations, four tissues and 12 time points which included direct (DIRE) contrasts to identify changes in expression in the N’Dama samples relative to the Boran samples (DIRE*_i_* = N*_i_* – B*_i_*, where N represents the N’Dama population and B represents the Boran population), N’Dama (NDAM) and Boran (BORA) contrasts to identify changes in expression within the populations relative to time 0 (NDAM*_i_* = N*_i_* – N_0_; BORA*_i_* = B*_i_* – B_0_) and response (RESP) contrasts to identify changes in expression over time in the N’Dama samples relative to the Boran (RESP*_i_* = [N*_i_* – N_0_] – [B*_i_* – B_0_]) (**Fig 1**, **Table S1**) [33]. Moderated *t*-statistics, moderated *F*-statistic, and log-odds of differential expression were calculated for the informative probe sets by empirical Bayes moderation of the standard errors towards a global value [70, 73]. Global Benjamini-Hochberg correction for multiple testing was applied across all contrasts [70, 74]. Probe sets with an adjusted *P*-value of ≤ 0.05 (B-H *P*_adj._ ≤ 0.05) were determined to be significantly differentially expressed [70]. The microarray probe sets were converted to genes using gprofiler2 (v. 0.2.2) [75] with R (v. 4.3.2).

The results of the differential expression analysis were visualised using ComplexUpset (v. 1.3.3) [76], dplyr (v. 1.1.2), ggh4x (v. 0.2.4), ggplot2 (v. 3.4.2), ggrepel (v. 0.9.3) [77], patchwork (v. 1.2.0), readr (v. 2.1.4), rlang (v. 1.1.1) [78], stringr (v. 1.5.0), and tidyr (v 1.3.0) [79] with R (v. 4.3.2). Colours were generated from khroma (v. 1.10.0) and viridis (v. 0.6.3).

### Functional enrichment analysis

Functional enrichment was performed and visualised using dplyr (v. 1.1.2), ggh4x (v. 0.2.4), ggplot2 (v. 3.4.2), ggrepel (v. 0.9.3), gprofiler2 (v. 0.2.2), magick (v. 2.8.1) [80] with ImageMagick (v. 6.9.12.96) [81], magrittr (v. 2.0.3), purrr (v. 1.0.1) [82], readr (v. 2.1.4), rlang (v. 1.1.1), scales (v. 1.2.1) [83], and stringr (v. 1.5.0) with R (v. 4.3.2). Colours were generated from khroma (v. 1.10.0) and viridis (v. 0.6.3). The background set was the set of informative probe sets with valid Ensembl IDs. The query sets included those that were significantly differentially expression (adjusted *P*-value ≤ 0.05) for each of the 64 contrasts. The analysis was restricted to Gene Ontology (GO) terms and driver GO terms were highlighted. Generic Enrichment Map (GEM) files were generated and used as input for EnrichmentMap (v. 3.3.6) [84] with Cytoscape (v. 3.8.0) [75, 85, 86]. AutoAnotate (v. 1.3.5) [87] was used to create clusters of GO terms with genes in common with clusterMaker2 (v. 2.0) [88] and annotate them with appropriate names using WordCloud (v. 3.1.4) [89]. The yFiles Layout Algorithms (v. 1.1.3) method [90] was used to remove overlaps in the networks.

## Results

### Data filtering and normalisation

Nine samples were indicated as outliers by two or more of the quality control tests and were therefore removed, leaving 211 samples for analysis (**Fig S1**, **Table 1**). After normalisation the data were filtered to retain only informative probe sets, which resulted in a data set of 11,050 probe sets or 45.89% of the initial 24,128 probe sets.

### Principal component analysis

Based on the top ten principal components (PCs), the first PC explained 82.86% of the total variation in the data for PC1–10 and separated the liver from the blood, lymph node, and spleen samples (**Fig 2**). The second PC explained a further 9.10% of the total variation for PC1–10 and separated the blood from the lymph node and spleen samples (**Fig 2**). The third and fourth PCs explained 2.24% and 1.78% of the total variation for PC1–10, respectively, and separated the samples according to days post infection (dpi) while the fifth PC, which explained 1.32% of the total variation in the data for PC1–10 separated the lymph node and spleen samples (**Fig S2**□**S4**).

**Fig 2.**
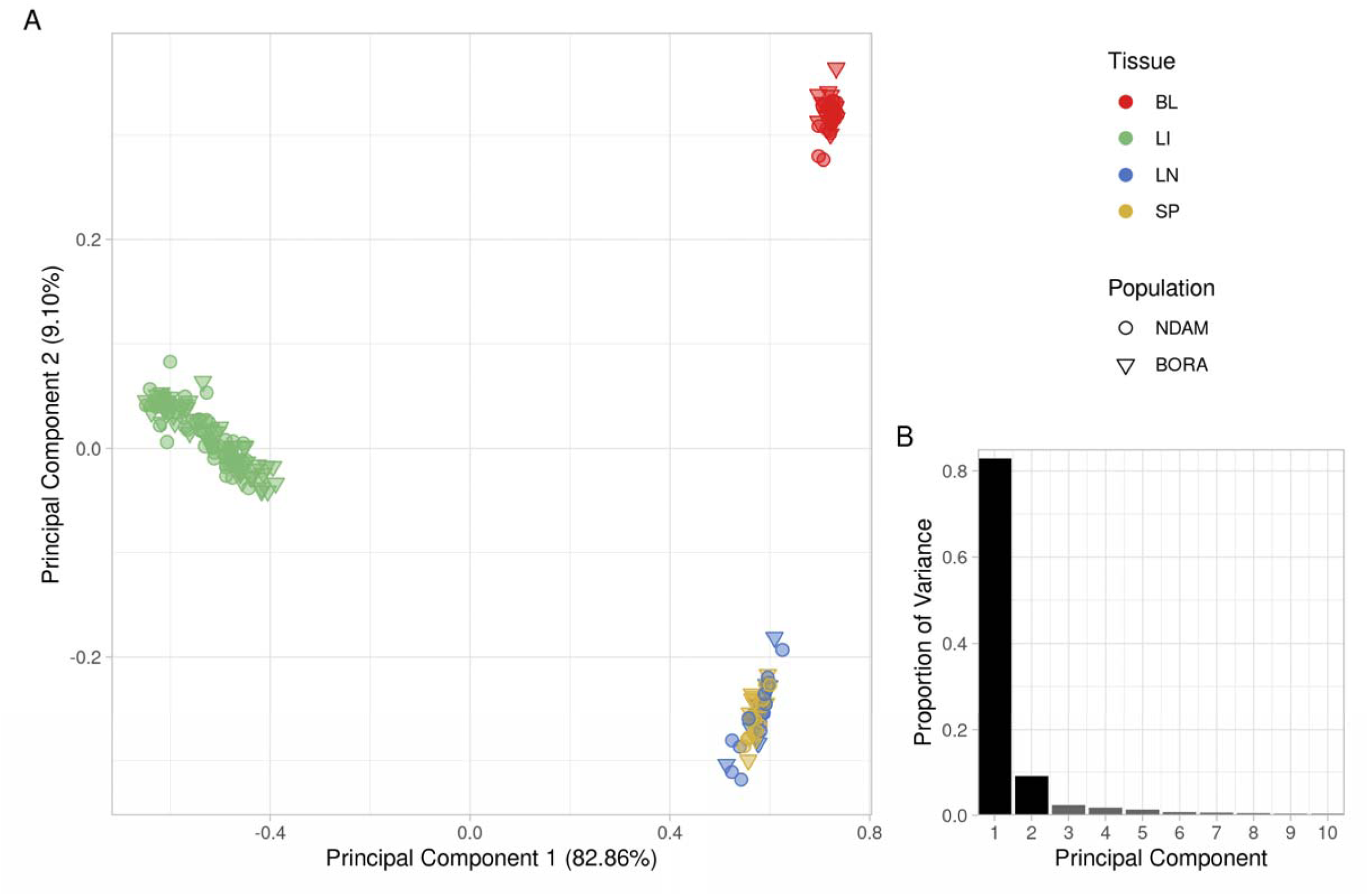
**A.** Principal component analysis (PCA) of the microarray data set with samples coloured according to tissue and with the shape indicating the population. The first two principal components (PC1 and PC2) are shown, and **B.** bar chart of proportion of variance of the top ten PCs.

### Differential expression analysis

The results of the differential expression analysis are detailed in **Table S2** and the number of significant (B-H *P*_adj._ ≤ 0.05) differentially expressed genes (DEGs) generally increased over time across all contrast types and tissues until 21 dpi when they began to decrease slightly before rising again to their peak at 34 or 35 dpi (**Fig 3**, **S5**□**S8**). The highest numbers of significant DEGs were found in the N’Dama and Boran contrasts, followed by the direct contrasts (**Fig 3**). Within the response contrasts the highest number of significant DEGs were found in the lymph node samples at 35 dpi (**Fig 3**, **S5**) while the blood samples at 34 dpi had the highest number of significant DEGs in the direct contrasts (**Fig 3**, **S6**). The highest number of significant DEGs in the N’Dama and Boran contrasts were at 35 dpi in the lymph node and liver samples, respectively (**Fig 3**, **S7**, **S8**). The numbers of significant DEGs were generally higher in the N’Dama contrasts than in the Boran contrasts early in the time course (**Fig 3**).

**Fig 3.**
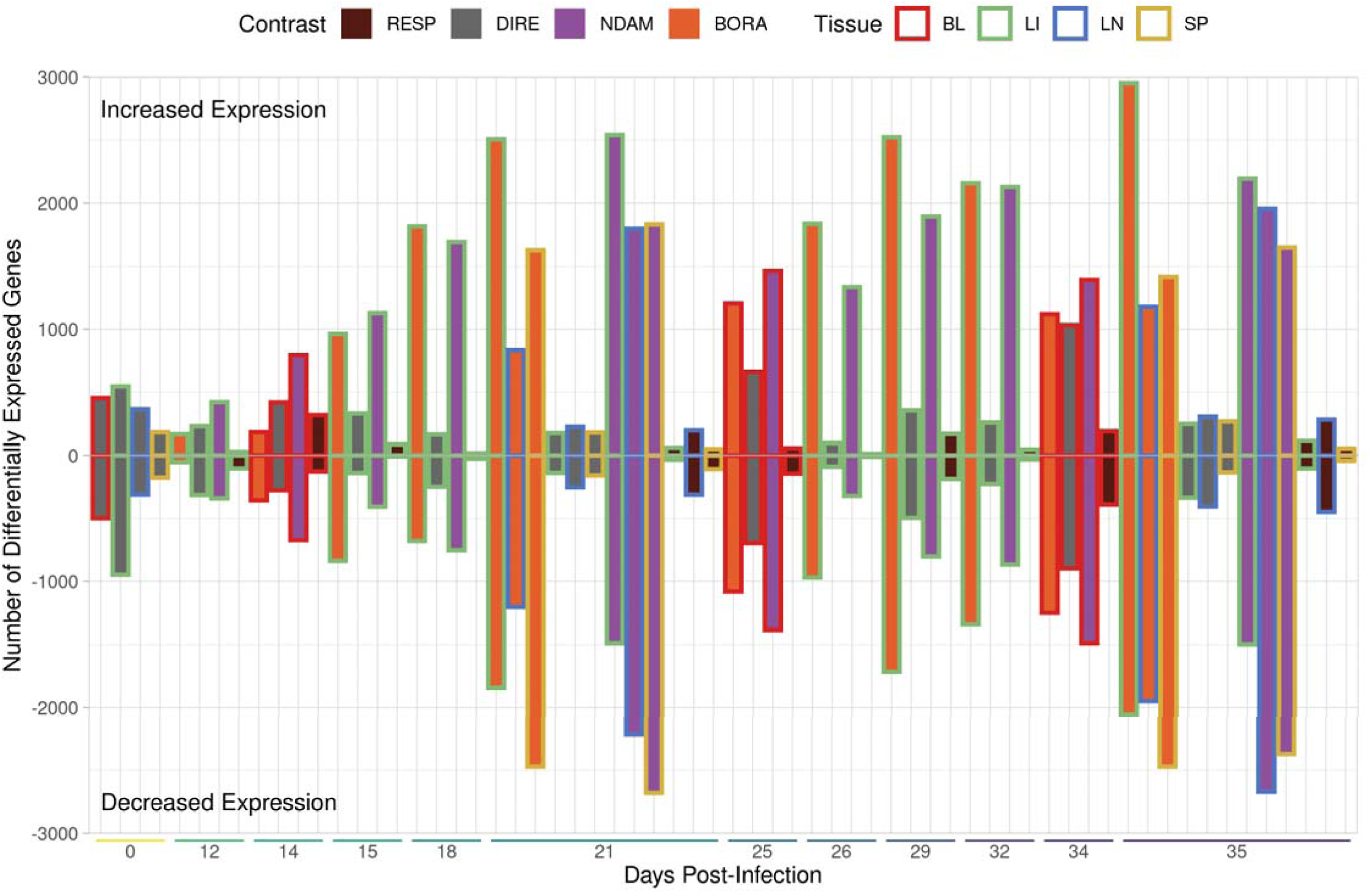
Bar chart showing the numbers of significant differentially expressed genes (DEGs) for all 64 contrasts (B-H *P*_adj_. ≤ 0.05). The extent of the bar above and below 0 on the *y*-axis indicates the numbers of significant DEGs with increased and decreased expression, respectively. The position on the *x*-axis indicates the number of days post infection (dpi). The bar colours represent the contrast type while the bar outline colours represent the tissue.

The N’Dama and Boran contrasts also had a higher number of significant DEGs in common between them when overlaps among all 64 contrasts were examined (**Fig S9**). Within the response and direct contrasts any overlaps in significant DEGs were generally confined to those between samples from the same tissue type at different time points (**Fig S10**, **S11**) while the N’Dama and Boran contrasts included more overlaps between tissues and time points (**Fig S12**, **S13**).

There were also many overlaps when the top 10 most significant genes with increased and decreased expression were examined for each of the contrasts (**Table 2**, **S3**□**S5**, **Fig 4**, **S14 S27**). The set of 1,267 genes identified as being in the top 10 most significant for increased or decreased expression for each of the 64 contrasts contained just 602 unique genes (**Table 2**, **S3**□**S5**). Of these, 328 were in the top 10 most significant for only one contrast, with the remaining 274 in the top 10 most significant for an average of 3.43 contrasts (**Table 2**, **S3**□**S5**). The response contrasts had the most unique genes in the top 10 most significant with 129 of the 328 unique top genes, while the direct, N’Dama and Boran contrasts had 75, 58 and 66 unique top genes, respectively (**Table 2**, **S3**□**S5**). The most common genes in the top 10 most significant differentially expressed for all the contrasts included *TMSB10,* which was in the top 10 most significant genes for 16 of the 64 contrasts, followed by *CYRIB*, *PTPRC*, *SPI1*, and *TTLL1*, which were in the top 10 most significant genes for 14 of the 64 contrasts (**Table 2**, **S3**□**S5**). The *TMSB10* gene was in the top 10 most significant genes with increased expression for the liver samples at all time points for the N’Dama and Boran contrasts (**Table S4**, **S5**). The *CYRIB*, *PTPRC*, and *SPI1* genes showed similar patterns with the exclusion of 12 dpi (**Table S4**, **S5**). *TTLL1* was in the top 10 most significant genes with increased expression in the blood samples at 25 and 34 dpi and the liver samples at 18 dpi for the response contrasts (**Table 2**). It was also in the top 10 most significant genes with increased expression for the blood samples at 0, 14, 25, and 34 dpi, the liver samples at 18 dpi, and the lymph node and spleen samples at 0, 21, and 35 dpi for the direct contrasts (**Table S3**).

**Fig 4.**
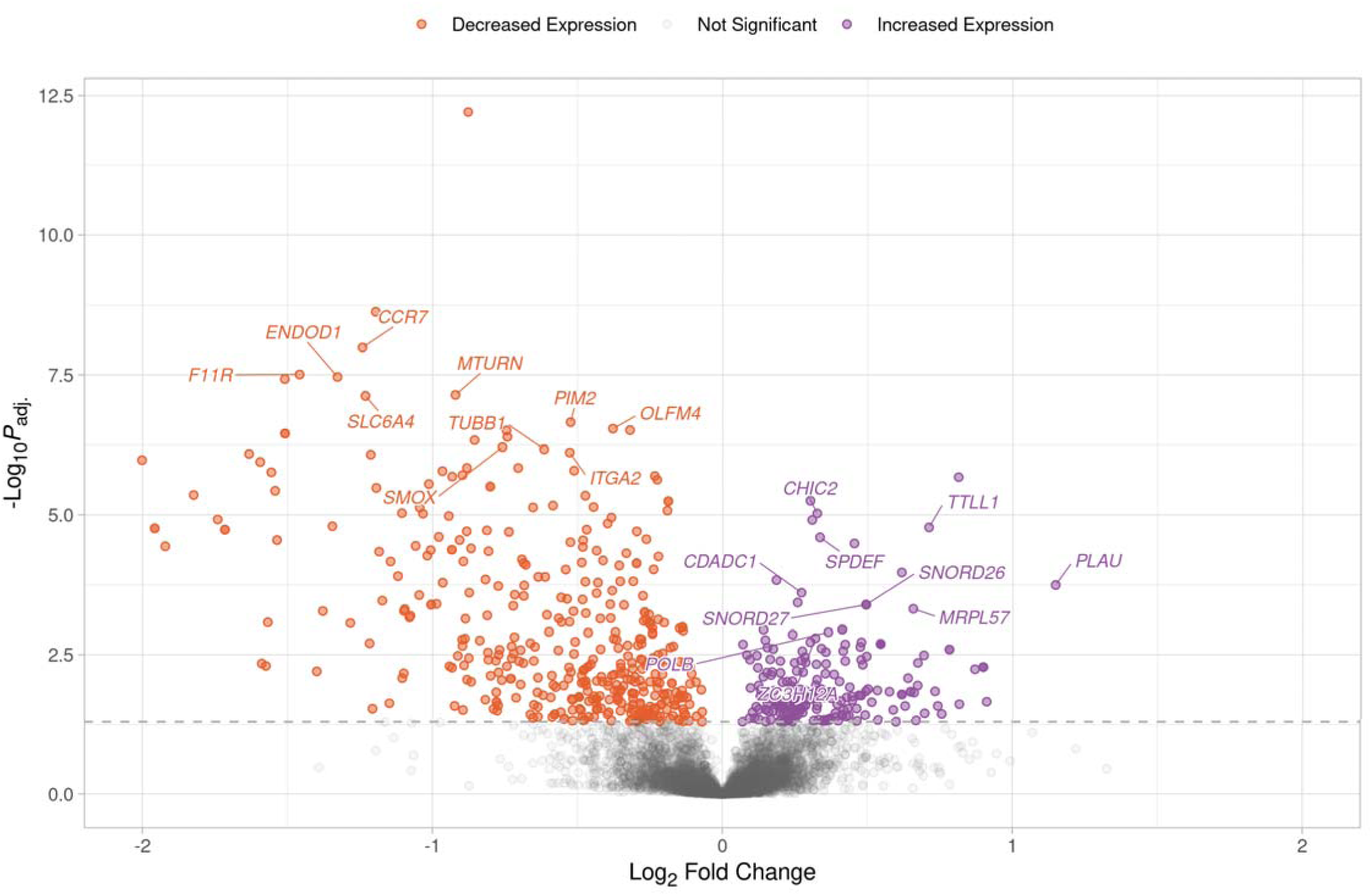
Volcano plot showing the results of the RESP contrast for the peripheral blood mononuclear cell (PBMC) samples at 34 days post infection (dpi). Each data point represents a gene with the position on the *x*- and *y*-axes indicating the log_2_ fold change and -log_10_*P*_adj._, respectively. Genes above the horizontal dashed line are significantly differentially expressed with the colours representing the change in expression. The top 10 mos significant genes for increased and decreased expression with gene symbols are labelled.

**Table 2.**
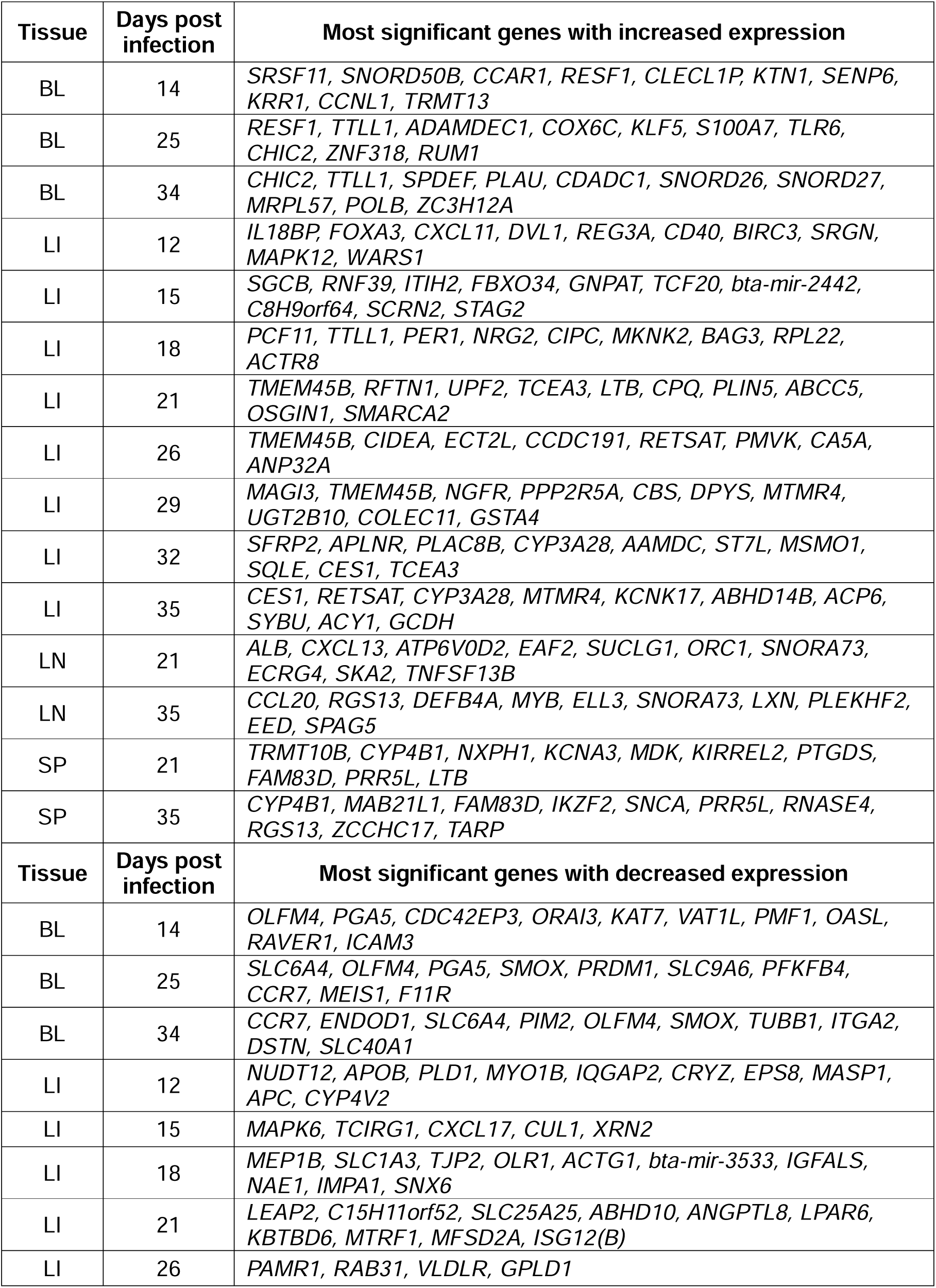
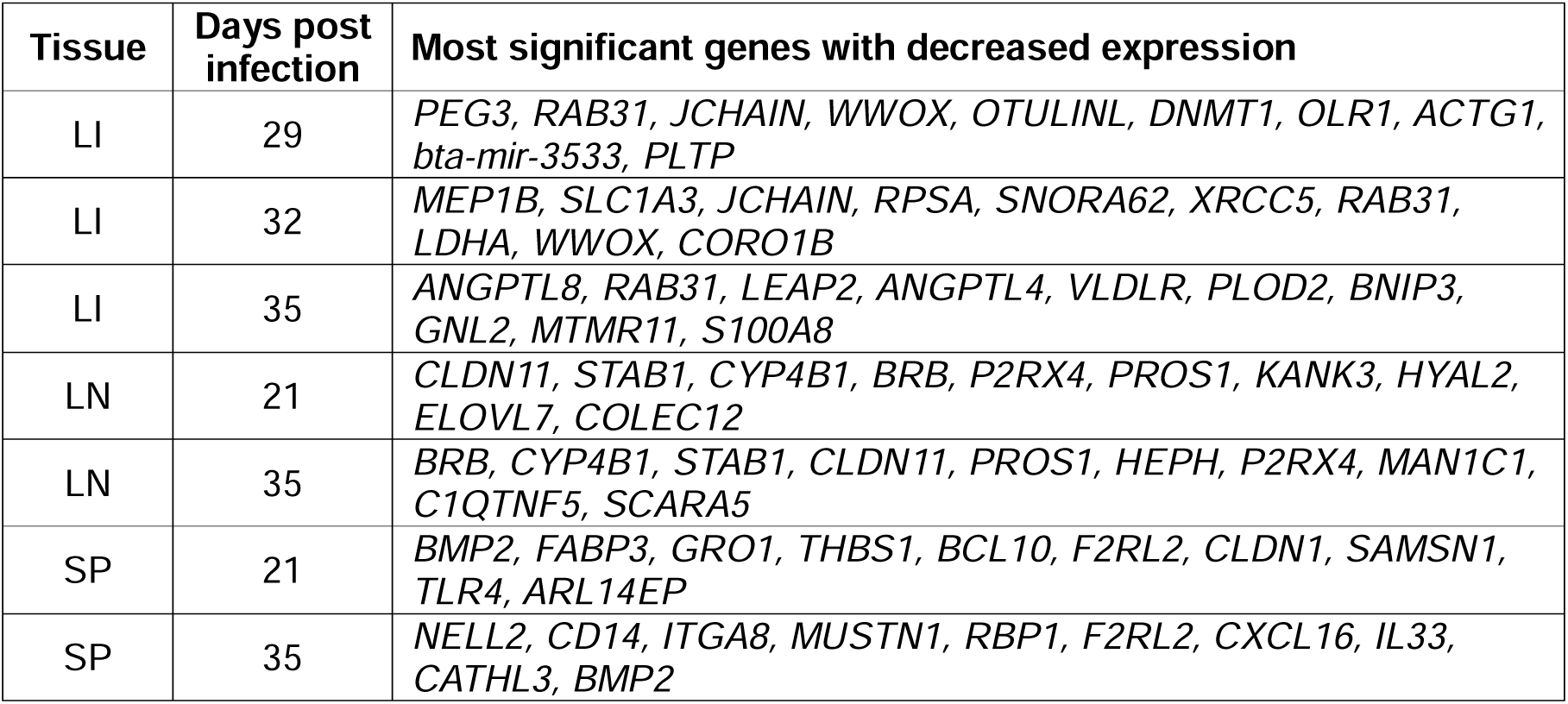
Tissue, days post infection (dpi) and the top 10 most significant genes with increased and decreased expression with valid gene symbols for the response contrasts.

The most common genes in the top 10 most significant differentially expressed for the response contrasts included *CYP4B1* and *RAB31,* which were in the top 10 most significant genes for four of the 15 response contrasts, followed by *OLFM4*, *TMEM45B*, and *TTLL1,* which were in the top 10 most significant genes for three of the 15 response contrasts (**Table 2**). For the response contrasts, *CYP4B1* was in the top 10 most significant genes with increased expression for the spleen samples at 21 and 35 dpi and the top 10 most significant genes with decreased expression for the lymph node samples at the same time point contrasts (**Table 2**). It was also in the top 10 most significant genes with increased expression for the spleen samples at 0 dpi and the top 10 most significant genes with decreased expression for the lymph node samples at the same time point for the direct contrasts (**Table S3**). *RAB31* was only in the top 10 most significant genes with decreased expression for the liver samples at 26, 29, 32, and 35 dpi for the response contrasts (**Table 2**). For the response contrasts, *OLFM4* was in the top 10 most significant genes with decreased expression for the blood samples at 14, 25, and 34 dpi (**Fig 4**, **Table 2**). It was also in the top 10 most significant genes with decreased expression for the blood samples at 0 dpi for the direct contrasts, and at 14, 25, and 34 dpi for the N’Dama contrasts (**Table S3**, **S4**). For the response contrasts, *TMEM45B* was in the top 10 most significant genes with increased expression for the liver samples at 21, 26, and 29 dpi (**Table 2**). It was also in the top 10 most significant genes with decreased expression for the liver samples at 0, 12, and 15 dpi for the direct contrasts (**Table S3**).

### Functional enrichment analysis

GO terms related to regulation of the mitotic cell cycle were significantly enriched for the genes with significantly increased expression in the blood and lymph node samples for the response contrasts (**Fig 5**). The top driver GO terms for the blood samples included *GO:0006396 RNA processing* and *GO:0005730 nucleolus* (**Fig S28**) while the top driver GO terms for the lymph node samples included *GO:0007059 chromosome segregation*, *GO:0005694 chromosome*, *GO:0006259 DNA metabolic process*, *GO:0006260 DNA replication*, *GO:0051301 cell division*, *GO:0003677 DNA binding*, and *GO:0008017 microtubule binding* (**Fig S30**). GO terms related to steroid metabolic processes (*GO:0008202 steroid metabolic process*) and oxidoreductase activity (*GO:0016705 oxidoreductase activity, acting on paired donors, with incorporation or reduction of molecular oxygen*, *GO:0016712 oxidoreductase activity, acting on paired donors, with incorporation or reduction of molecular oxygen, reduced flavin or flavoprotein as one donor, and incorporation of one atom of oxygen*) were significantly enriched for the genes with significantly increased expression in the liver samples for the response contrasts (**Fig 5**, **S29**). No GO terms were significantly enriched for genes with significantly increased expression in the spleen samples for the response contrasts (**Fig 5**).

**Fig 5.**
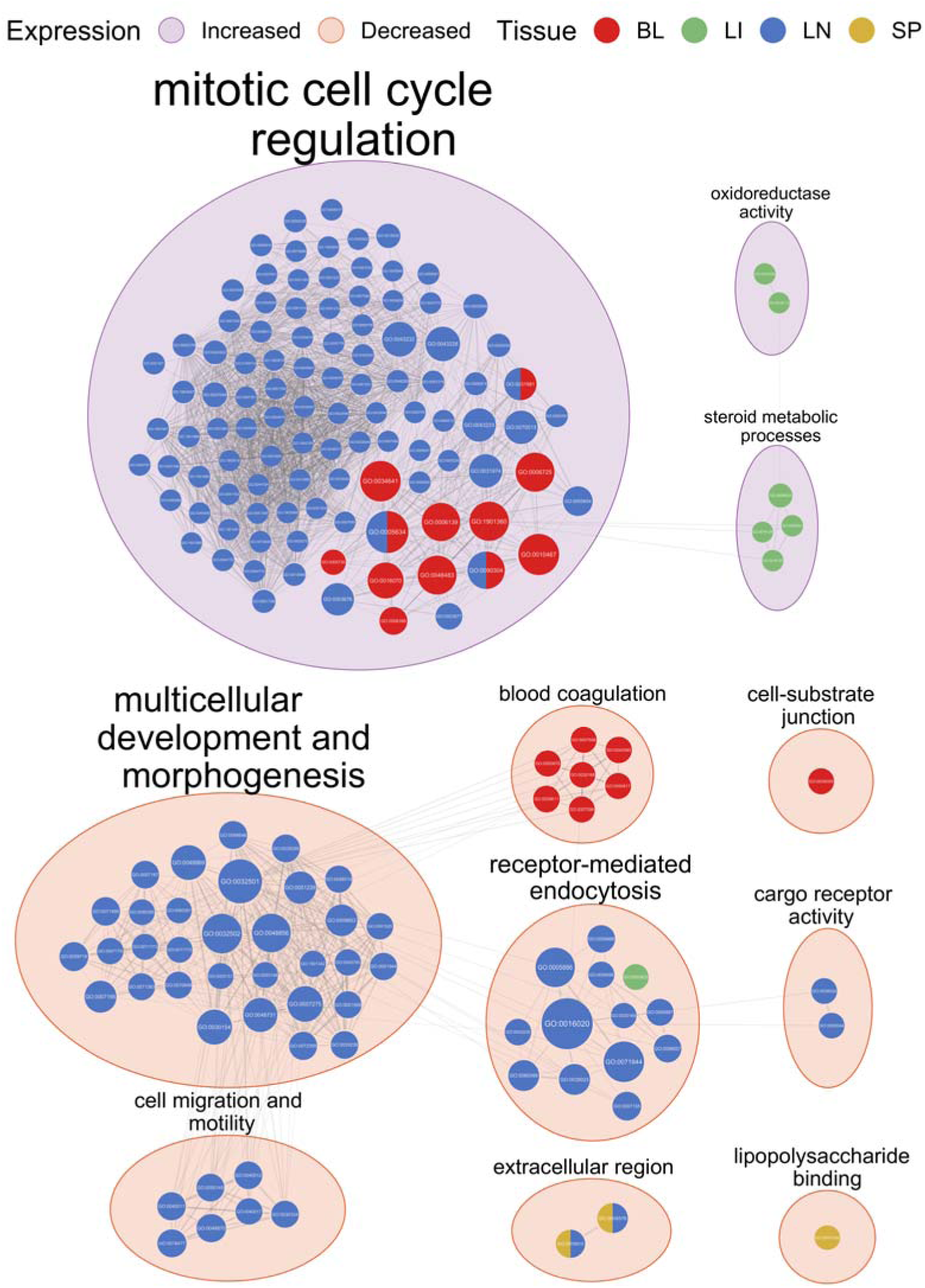
EnrichmentMap network of significantly enriched gene ontology (GO) terms identified from g:Profiler functional enrichment of significantly differentially expressed genes (DEGs) for the response contrasts. Each node represents a GO term with the colour of the node representing the tissue and the size representing the number of genes in the GO term. The edges indicate overlap between the GO terms with the width of the edges representing the similarity coefficient for the connected GO terms. The GO terms are clustered by AutoAnotate with the background colour of the clusters representing the direction of expression. The clusters are labelled with the size of the label scaling with the number of GO terms in the cluster.

GO terms related to blood coagulation (*GO:0007596 blood coagulation*) and the cell-substrate junction (*GO:0030055 cell-substrate junction*) were significantly enriched for the genes with significantly decreased expression in the blood samples for the response contrasts (**Fig 5**, **S28**). GO terms related to receptor-mediated endocytosis were significantly enriched for the genes with significantly decreased expression in the liver and lymph node samples for the response contrasts (**Fig 5**). The top driver GO terms included *GO:0003823 antigen binding* for the liver samples (**Fig S29**), and *GO:0071944 cell periphery*, *GO:0009986 cell surface*, *GO:0007155 cell adhesion*, and *GO:0006897 endocytosis* for the lymph node samples (**Fig S30**). GO terms related to multicellular development and morphogenesis including the driver GO terms *GO:0007167 enzyme-linked receptor protein signaling pathway*, *GO:0032501 multicellular organismal process*, and *GO:0071495 cellular response to endogenous stimulus* were also significantly enriched for the genes with significantly decreased expression in the lymph node samples for the response contrasts (**Fig 5**, **S30**). GO terms related to cell migration and motility (*GO:0016477 cell migration*, *GO:0048870 cell motility*) and cargo receptor activity (*GO:0038024 cargo receptor activity*) were also enriched for these genes (**Fig 5**, **S30**). GO terms related to the extracellular region (*GO:0005576 extracellular region*) were significantly enriched for the genes with significantly decreased expression in both the lymph node and spleen samples for the response contrasts (**Fig 5**, **S30**, **S31**), while GO terms related to lipopolysaccharide binding (*GO:0001530 lipopolysaccharide binding*) were also significantly enriched for the genes with significantly decreased expression in the spleen samples for the response contrasts (**Fig 5**, **S31**).

For the direct contrasts, GO terms related to organelle metabolic processes were significantly enriched for the genes with significantly increased expression in the blood, liver, and lymph node samples (**Fig S32**). The GO term for leukocyte proliferation was significantly enriched for genes with significantly increased expression in the blood samples, while GO terms related to scavenger receptor activity and cell periphery were also significantly enriched for genes with significantly increased expression in the lymph node samples (**Fig S32**). GO terms related to receptor signalling pathways were significantly enriched for genes with significantly decreased expression in the liver and lymph node samples, while GO terms related to coagulation were significantly enriched for genes with significantly decreased expression in the blood samples (**Fig S32**). GO terms related to actin binding and cytoskeleton, immune response and antigen binding, small molecule and pyruvate metabolic processes, and the cytosolic ribosome were also significantly enriched for genes with significantly decreased expression in the liver samples (**Fig S32**). GO terms related to tube development, cell migration, extracellular region, cell adhesion, and calcium ion binding were significantly enriched for genes with significantly decreased expression in the lymph node samples (**Fig S32**). GO terms related to metaphase chromosome alignment, the chromosome centromeric region, and the microtubule cytoskeleton were significantly enriched for genes with significantly decreased expression in the spleen samples (**Fig S32**).

For the N’Dama contrasts, GO terms related to cell process regulation were significantly enriched for genes with significantly increased expression in all tissues (**Fig S33**). GO terms related to GTPase activator activity, phagocytosis, and protein kinase binding were also significantly enriched for genes with significantly increased expression in the liver samples (**Fig S33**). GO terms related to cell development and regulation were significantly enriched for genes with significantly decreased expression in all tissues (**Fig S33**). GO terms related to metabolic processes, peroxisome organization, electron transfer activity, oxidoreductase activity, and the endoplasmic reticulum membrane were enriched for genes with significantly decreased expression in the liver samples (**Fig S33**). GO terms related to actin cytoskeleton organization, extracellular matrix, cell projection, and vesicle-mediated transport were significantly enriched for genes with significantly decreased expression in the lymph node and spleen samples (**Fig S33**). GO terms related to the extracellular region were also significantly enriched for genes with significantly decreased expression in the lymph node samples, while GO terms related to extracellular matrix organization were significantly enriched for genes with significantly decreased expression in the spleen samples (**Fig S33**). For the Boran contrasts, GO terms related to cell process regulation were significantly enriched for genes with significantly increased expression in all tissues and GO terms related to cell response regulation were significantly enriched for genes with significantly decreased expression in all tissues, while GO terms related to metabolic processes were also significantly enriched for genes with significantly decreased expression in the liver and spleen samples (**Fig S34**).

## Discussion

The data filtering and normalisation steps resulted in a similar number of samples and probe sets for analysis as previous studies that used Affymetrix^®^ Bovine Genome Array data sets to examine host responses to trypanosomiasis and other infectious diseases of cattle, including the previous analysis of the liver, lymph node, and spleen samples [33, 91–93]. The clear separation of the liver samples from the other tissues that is evident for PC1, which also explained the majority of the total variation in the data set for PC1–10, may be due to several factors (**Fig 2**). The first is biological differentiation between the liver samples and those of the other tissues. This is supported by a previous gene expression microarray study using rat (*Rattus norvegicus*) tissues, which also observed that PC1 in a PCA separated liver samples from those taken from blood, lymph node, spleen, and other tissues [94]. This biological difference may have been compounded by the higher number of liver samples in the data set from Noyes and colleagues [33] due to the experimental design, which encompassed a larger number of time points for the liver samples as well as a greater number of animals from which the liver samples were taken to prevent the collection of multiple liver biopsies from the same animal at consecutive time points [33]. This experimental design consideration also resulted in more control liver samples for the pre-infection time point [33]. In total, 115 of the 211 samples that passed the quality control filters or 54.50% of the filtered data set were liver samples (**Table 1**). Because an unequal number of samples is known to increase the distance between groups with larger sample sizes in a PCA, it is possible that the disproportionately large number of liver samples in the present study has given rise to the effect seen in PC1 [95–97]. It is therefore unsurprising that PC2, which explained a further 9.10% of the total variation in the data set for PC1–10, separated the blood samples from the lymph node and spleen samples (**Fig 2**) as the blood samples were the second most numerous in the data set with 40 samples or 18.96% of the filtered data set (**Table 1**). It is not until PC5, which explained 1.32% of the total variation in the data set for PC1–10, that the lymph node and spleen samples are separated (**Fig S4**). The samples from these tissues are the least numerous in the data set with 29 spleen and 27 lymph node samples, making up 13.74% and 12.80% of the filtered data set, respectively (**Table 1**). The spleen and lymph nodes are also both key components of the lymphatic and immune systems and their functional and morphological similarities as hemopoietic organs that filter bodily fluids have been long established [98–100]. It is logical, therefore, that these tissues would show similar patterns of gene expression both before and after infection. The third and fourth PCs (PC3 and PC4), which explained 2.24% and 1.78% of the total variation for PC1–10, respectively, separated the samples temporally in order of dpi (**Fig S2**, **S3**). This illustrates the overall similarity in response to infection across the populations and is in agreement with RNA-seq data from whole blood samples taken during a similar trypanosome infection time course experiment [42]. It also shows that the systemic response to trypanosome infection in peripheral blood is distinct from that in the liver and other tissues and that blood sampling alone may not be sufficient to capture all the tissue specific responses to infection.

Gene expression in bovine tissues can be measured using a variety of methods, one of which is the Affymetrix^®^ Bovine Genome Array based on GeneChip^®^ technology [101] and which contains more than 24,000 probe sets representing over 23,000 gene transcripts. The Affymetrix^®^ Bovine Genome Array platform has been previously used to study infectious disease in cattle, including trypanosomiasis [33], and for mycobacterial infections that cause tuberculosis and paratuberculosis [91–93]. Importantly, although expression microarrays have been largely supplanted by RNA sequencing (RNA-seq), when the same biological samples were analysed using both the Bovine Genome Array and RNA-seq, gene expression data obtained using the two platforms were extremely well correlated in peripheral blood and alveolar macrophages [102–104]. In addition, it has recently been shown that peripheral blood gene expression data generated using RNA-seq, a different microarray technology (long oligonucleotide microarrays), and reverse transcription quantitative real-time PCR (RT-qPCR) were well correlated in independent studies of trypanosome infection in cattle [42].

The pattern of an increase in the numbers of significant DEGs until 21 days post infection followed by a decrease before rising to a final peak at 35 days post infection across contrast types and tissues (**Fig 3**, **S5**□**S8**) is consistent with known information about the cyclic nature of parasitaemia during trypanosome infection, which has been shown to have an initial peak at 20 dpi [38]. It is also in agreement with the measures of parasitaemia in the blood samples during this infection time course experiment, which showed peaks of parasitaemia from 15 to 22 dpi [41]. The final peak of significant DEGs at the end of the infection time course is consistent with analysis of the same blood samples using a gene expression platform with less gene content—a bovine long oligonucleotide (BLO) array— which detected the highest number of significant DEGs between the two populations at 34 dpi [40]. It is also in agreement with a study using RNA-seq data from whole blood samples that found the highest number of DEGs in N’Dama samples at the end of a similar trypanosome infection time course experiment [42]. The higher numbers of significant DEGs in the N’Dama contrasts when compared to the Boran contrasts is in agreement with previous results and supports the hypothesis that, compared to trypanosusceptible Boran, trypanotolerant N’Dama cattle exhibit an earlier proinflammatory response, which is evident in both PBMC and whole peripheral blood [40–42]. That the response contrasts had the lowest number of significant DEGs is to be expected given that these contrasts are examining the data set for differences in the rate of change of gene expression between the populations [33]. In addition, the numbers of significant DEGs for the response contrasts were similar to those found in the original analysis by Noyes and colleagues [33], which employed a similar approach. The numbers of genes in common between the contrasts is to be expected due to the shared underlying data set while the numbers of genes in common within the contrasts is likely due to similar patterns of gene expression found across the populations, tissues, and time points during the infection (**Fig S9**□**S13**). This is in agreement with RNA-seq data that showed large overlaps in DEGs in response to trypanosome infection between five different cattle populations with varying levels of trypanotolerance [42].

The similarities in gene expression are also illustrated by the high level of overlap in the top 10 most significant DEGs with increased and decreased expression between the contrasts (**Table 2**, **S3**□**S5**). The most common genes in the top 10 most significant DEGs for all the contrasts included those related to the immune system, such as the most common significant DEG—the thymosin beta 10 gene (*TMSB10*), which has been identified as a hub gene in the response to Rift Valley fever (RVF), an important viral disease in African cattle [105]. Other immune-related genes in the set of the most common genes in the top 10 most significant DEGs for all the contrasts included the CYFIP related Rac1 interactor B gene (*CYRIB*), which encodes a regulator of phagocytosis [106], and the protein tyrosine phosphatase receptor type C gene (*PTPRC*), which is an essential regulator of T and B cell antigen receptor-mediated activation and has been highlighted by multiple studies of trypanosome infection in mice [107–112]. Similarly, the Spi-1 proto-oncogene gene (*SPI1*), which encodes a transcription factor that activates gene expression during myeloid and B-lymphoid cell development, has also been implicated as an important host gene in trypanosome infection in the mouse [111, 113]. The presence of immune genes in the top 10 most significant DEGs is to be expected given the nature of the infection time course experiment and is in agreement with previous results from this and other infection experiments, illustrating the similarity in immune responses between the N’Dama and Boran cattle during trypanosome infection [33, 40, 42].

The most common genes in the top 10 most significant DEGs for the response contrasts included genes which are involved in in inflammation and immunity such as the olfactomedin 4 gene (*OLFM4*) and the transmembrane protein 45B gene (*TMEM45B*) [114, 115]. Other common genes in the top 10 most significant DEGs for the response contrasts are more varied in function and included the cytochrome P450 family 4 subfamily B member 1 gene (*CYP4B1*), which is involved in drug metabolism [116]; the RAB31, member RAS oncogene family gene (*RAB31*), which is a small GTPase [117]; and the TTL family tubulin polyglutamylase complex subunit L1 gene (*TTLL1*), which is involved in microtubule cytoskeleton organisation [118].

The top 10 most significant DEGs for the response contrasts also included genes encoding antimicrobial peptides (AMPs), which are key components of the innate immune system that have huge therapeutic potential and are important for the healthy function of a variety of bovine tissues [119–122]. This is in agreement with previous RT-qPCR results using blood samples from the same time course infection experiment; in this regard, it has previously been hypothesised that trypanotolerance may be partly due to inherited regulatory sequence variation in genes encoding antimicrobial peptides [39]. For example, the liver enriched antimicrobial peptide 2 gene (*LEAP2*) [123], which was in the top 10 most significant DEGs with decreased expression for the liver samples at 21 and 35 dpi for the response contrasts (**Table 2**). The previous RT-qPCR results showed that *LEAP2* exhibited significantly decreased expression in N’Dama samples at 34 dpi relative to day 0 while the Boran animals showed no such change [39]. Other AMP genes in the top 10 most significant DEGs for the response contrasts which examine the N’Dama samples relative to the Boran samples included the cathelicidin antimicrobial peptide gene (*CATHL3*) [124] with decreased expression in the spleen samples at 35 dpi, which is notable since cathelicidins were found to be the most effective of the three classes of AMP at killing both insect and bloodstream forms of *T. brucei* in mice [125]; the defensin beta 4A gene (*DEFB4A*) [126] with increased expression in the lymph node samples at 35 dpi; and S100 calcium binding protein A7 gene (*S100A7*) [127] with increased expressed in peripheral blood at 25 dpi (**Table 2**) [39, 124, 126, 127].

Another group of genes represented in the top 10 most significant DEGs for the response contrasts were cytokine genes (**Table 2**). This is consistent with both RT-qPCR and BLO microarray results from the same infection time course experiment and previous studies using the same animals [33, 40, 41]. This observation agrees with more recent transcriptomics studies from trypanotolerant and trypanosusceptible cattle and studies of trypanosome infection in the mouse [42, 128]. These genes included the C-C motif chemokine ligand 20 gene (*CCL20*) with increased expression in the lymph node samples at 35 dpi; the C-X-C motif chemokine ligand 11 (*CXCL11*) with increased expression in the liver samples at 12 dpi; the C-X-C motif chemokine ligand 13 gene (*CXCL13*) with increased expression in the lymph node samples at 21 dpi; the C-X-C motif chemokine ligand 16 gene (*CXCL16*) with decreased expression in the spleen samples 35 dpi; the C-X-C motif chemokine ligand 17 gene (*CXCL17*) with decreased expression in the liver samples at 15 dpi; the interleukin 33 gene (*IL33*) with decreased expression in the lymph node samples at 35 dpi; and the TNF superfamily member 13b gene (*TNFSF13B*) with increased expression in the lymph node samples at 21 dpi (**Table 2**).

Related genes such as cytokine receptors and inhibitors were also in the top 10 most significant DEGs for the response contrasts (**Table 2**). These included the C-C motif chemokine receptor 7 gene (*CCR7*) with decreased expression in the blood samples at 25 and 34 dpi; the C1q and TNF related 5 gene (*C1QTNF5*) with decreased expression in lymph node samples at 35 dpi; and the interleukin 18 binding protein gene (*IL18BP*) with increased expression in the liver samples at 12 dpi (**Table 2**). Additionally, mitogen-activated protein kinase (MAPK) genes were present in the top 10 most significant DEGs, which is in agreement with previous work using the same animals and also a more recent study of trypanosome infection in cattle [33, 40, 42]. In this regard, it is notable that MAPK signalling pathways are involved in the response to proinflammatory cytokines and are known to be subverted by *T. cruzi* and other trypanosomatid parasites to evade the host immune response [129–131].

A final notable group of genes present in the top 10 most significant DEGs were those encoding apolipoproteins, including apolipoprotein B (*APOB*) with decreased expression for the response contrast in the liver samples at 12 dpi (**Table 2**); apolipoprotein L3 (*APOL3*) with increased expression for the direct contrast in the blood samples at 25 dpi (**Table S3**); and apolipoprotein M (*APOM*) with decreased expression for the Boran contrast in the liver samples at 15 dpi (**Table S5**). These genes are related to apolipoprotein 1 (*APOL1*) which encodes the protein that makes up the trypanosome lytic factor (TLF) that is present in human and western lowland gorilla (*Gorilla gorilla gorilla*) serum [132]. TLF is taken up by susceptible trypanosomes where it interferes with their lysosomes and mitochondria, thereby conferring host resistance to most trypanosome species [132]. It is therefore interesting that related genes are among the top 10 most differentially expressed in this study.

Gene ontology terms related to regulation of the mitotic cell cycle, steroid metabolic processes and oxidoreductase activity (**Fig 5**) were significantly enriched for the genes with significantly increased expression for the response contrasts, which is in agreement with results from a previous study using the same animals [33]. The overlap and grouping of the GO terms between the blood and lymph node samples illustrates the similarity of responses to infection for these tissues. The lack of GO terms significantly enriched for genes with significantly increased expression for the response contrasts in the spleen samples (**Fig 5**) can be explained by the lower numbers of significant DEGs with increased expression for these samples (**Fig 3**, **S5**). A larger range of GO terms were significantly enriched for the genes with significantly decreased expression for the response contrasts (**Fig 5**). This is likely due to the higher number of significant DEGs with decreased expression, which would allow more GO terms to be significantly enriched for these genes (**Fig 3**). The GO terms significantly enriched for the genes with significantly decreased expression for the response contrasts included those related to multicellular development and morphogenesis, receptor-mediated endocytosis, cell migration and motility, cargo receptor activity, extracellular region, and lipopolysaccharide binding (**Fig 5**). These observations are also in agreement with results of the previous analysis using the same animals [33].

It is logical that the GO terms that are significantly enriched for genes with significantly decreased expression for the response contrasts are related to blood coagulation in the PBMC samples (**Fig 5**, **S28**). The effects of blood-borne trypanosome parasites on the blood of the host have been long studied in both cattle and humans [133–135]. Notably, anaemia is the main cause of death due trypanosomiasis and the ability to control this anaemia is consider critical to trypanotolerance in cattle [136, 137]. While trypanosomes do obtain iron from their environment in the blood of the host and that iron homeostasis and metabolism in the trypanosome parasite are known to be essential for infection, it has been observed that anaemia does not correlate with parasitaemia [138–141]. Additionally, trypanosomes require much less iron than a mammalian cell [142, 143]. The anaemia caused by trypanosome infection is therefore considered to be an immune response, which may be driven by cytokines [137, 144, 145].

During trypanosome infection in cattle, control of anaemia and parasitaemia are key components of trypanotolerance but are considered to represent separate genetically determined traits [146]. This is exemplified by genes related to iron ion homeostasis such as the solute carrier family 40 member 1 gene (*SLC40A1*) [147], which was noted to be among the most divergent between trypanotolerant N’Dama and trypanosusceptible Boran as a result of a marked reduction in expression in the N’Dama population over the course of the infection experiment relative to the pre-infection level in PBMC [40]. *SLC40A1* was also in the top 10 most significant DEGs with decreased expression in the blood samples at 34 dpi for the response contrast in this study, highlighting the divergent nature of the expression of this gene between the N’Dama and Boran populations during trypanosome infection (**Table 2**). The reduction of cellular iron export by the SLC40A1 protein is thought to be a component of an innate immune-driven strategy to prevent bloodstream pathogens from accessing iron, with anaemia as a side effect for the host [137, 140, 144]. A related gene, the solute carrier family 11 member 1 gene (*SLC11A1*), which regulates iron homeostasis in macrophages was found to be differentially expressed between N’Dama and African indicine cattle during trypanosome infection using RNA-seq data [42] and variants of this gene are associated with susceptibility to several infectious diseases, including tuberculosis in cattle [148, 149]. Finally, it is noteworthy that the previous work by Noyes and colleagues [33], which used a different functional enrichment approach also highlighted biological pathways related to iron ion homeostasis.

The significantly enriched GO terms showed similar patterns for the direct contrasts with those enriched for the genes with significantly increased expression related to organelle metabolic processes for multiple tissues, while those enriched for the genes with significantly decreased expression showed a greater range of processes (**Fig S32**). Gene ontology terms related to coagulation were also significantly enriched for genes with significantly decreased expression in the PBMC samples (**Fig S32**). For the N’Dama and Boran contrasts, the significantly enriched GO terms were related to cell regulation across all tissues, time points, and direction of expression (**Fig S33**, **S34**). This is likely a result of the higher numbers of significant DEGs for these contrasts, which led to higher numbers of significantly enriched GO terms that were too varied for the EnrichmentMap method to summarise effectively (**Fig 3**) [84].

In conclusion, trypanotolerant N’Dama and trypanosusceptible Boran cattle responded in largely similar ways during trypanosome infection when gene expression was examined using PBMC, liver, lymph node, and spleen samples with peaks and troughs of gene expression following the cyclic pattern of parasitaemia exhibited during trypanosome infection. Differences in response to infection between the two populations include genes related to the immune system such as those encoding antimicrobial peptides and cytokines. Within the PBMC samples, differences in genes relating to coagulation and iron homeostasis support the hypothesis that the dual abilities to control both parasitaemia and the anaemia resulting from the innate immune response to trypanosome parasites are key to trypanotolerance. This work adds to our understanding of host-trypanosome interactions across multiple tissues and time points as well as to our knowledge of how genetic variation underlying the host response can lead to differential host tolerance of, or susceptibility to, trypanosome infection. This improves our general understanding of mammalian trypanosome infection which may also be applicable to human African trypanosomiasis.

## Acknowledgements

We thank Morris Agaba, Olivier Hanotte, Stephen J. Kemp, and Stephen V. Gordon for assistance with sample resources and for useful scientific discussion.

