## Supporting information (supplemental tables and figures) for "Functional genomics of trypanotolerant and trypanosusceptible cattle infected with *Trypanosoma congolense* across multiple time points and tissues"

**Table S1.** Contrast ID, contrast type, tissue, days post infection (dpi), and formula for each of the 64 contrasts.

| Contrast ID | Contrast | Tissue | dpi | Contrast formula |
|---|---|---|---|---|
| RESP BL 14 | RESP | BL | 14 | (NDAM BL 14 - NDAM BL 00) - (BORA BL 14 - BORA BL 00) |
| RESP BL 25 | RESP | BL | 25 | (NDAM BL 25 - NDAM BL 00) - (BORA BL 25 - BORA BL 00) |
| RESP BL 34 | RESP | BL | 34 | (NDAM BL 34 - NDAM BL 00) - (BORA BL 34 - BORA BL 00) |
| RESP LI 12 | RESP | LI | 12 | (NDAM LI 12 - NDAM LI 00) - (BORA LI 12 - BORA LI 00) |
| RESP LI 15 | RESP | LI | 15 | (NDAM LI 15 - NDAM LI 00) - (BORA LI 15 - BORA LI 00) |
| RESP LI 18 | RESP | LI | 18 | (NDAM LI 18 - NDAM LI 00) - (BORA LI 18 - BORA LI 00) |
| RESP LI 21 | RESP | LI | 21 | (NDAM LI 21 - NDAM LI 00) - (BORA LI 21 - BORA LI 00) |
| RESP LI 26 | RESP | LI | 26 | (NDAM LI 26 - NDAM LI 00) - (BORA LI 26 - BORA LI 00) |
| RESP LI 29 | RESP | LI | 29 | (NDAM LI 29 - NDAM LI 00) - (BORA LI 29 - BORA LI 00) |
| RESP LI 32 | RESP | LI | 32 | (NDAM LI 32 - NDAM LI 00) - (BORA LI 32 - BORA LI 00) |
| RESP LI 35 | RESP | LI | 35 | (NDAM LI 35 - NDAM LI 00) - (BORA LI 35 - BORA LI 00) |
| RESP LN 21 | RESP | LN | 21 | (NDAM LN 21 - NDAM LN 00) - (BORA LN 21 - BORA LN 00) |
| RESP LN 35 | RESP | LN | 35 | (NDAM LN 35 - NDAM LN 00) - (BORA LN 35 - BORA LN 00) |
| RESP SP 21 | RESP | SP | 21 | (NDAM SP 21 - NDAM SP 00) - (BORA SP 21 - BORA SP 00) |
| RESP SP 35 | RESP | SP | 35 | (NDAM SP 35 - NDAM SP 00) - (BORA SP 35 - BORA SP 00) |
| DIRE BL 00 | DIRE | BL | 0 | NDAM BL 00 - BORA BL 00 |
| DIRE BL 14 | DIRE | BL | 14 | NDAM BL 14 - BORA BL 14 |
| DIRE BL 25 | DIRE | BL | 25 | NDAM BL 25 - BORA BL 25 |
| DIRE BL 34 | DIRE | BL | 34 | NDAM BL 34 - BORA BL 34 |
| DIRE LI 00 | DIRE | LI | 0 | NDAM LI 00 - BORA LI 00 |
| DIRE LI 12 | DIRE | LI | 12 | NDAM LI 12 - BORA LI 12 |
| DIRE LI 15 | DIRE | LI | 15 | NDAM LI 15 - BORA LI 15 |
| DIRE LI 18 | DIRE | LI | 18 | NDAM LI 18 - BORA LI 18 |
| DIRE LI 21 | DIRE | LI | 21 | NDAM LI 21 - BORA LI 21 |
| DIRE LI 26 | DIRE | LI | 26 | NDAM LI 26 - BORA LI 26 |
| DIRE LI 29 | DIRE | LI | 29 | NDAM LI 29 - BORA LI 29 |
| DIRE LI 32 | DIRE | LI | 32 | NDAM LI 32 - BORA LI 32 |
| DIRE LI 35 | DIRE | LI | 35 | NDAM LI 35 - BORA LI 35 |

**Table S1 continued.**

| Contrast ID | Contrast | Tissue | dpi | Contrast formula |
|---|---|---|---|---|
| DIRE LN 00 | DIRE | LN | 0 | NDAM LN 00 - BORA LN 00 |
| DIRE LN 21 | DIRE | LN | 21 | NDAM LN 21 - BORA LN 21 |
| DIRE LN 35 | DIRE | LN | 35 | NDAM LN 35 - BORA LN 35 |
| DIRE SP 21 | DIRE | SP | 21 | NDAM SP 21 - BORA SP 21 |
| DIRE SP 35 | DIRE | SP | 35 | NDAM SP 35 - BORA SP 35 |
| NDAM BL 14 | NDAM | BL | 14 | NDAM BL 14 - NDAM BL 00 |
| NDAM BL 25 | NDAM | BL | 25 | NDAM BL 25 - NDAM BL 00 |
| NDAM BL 34 | NDAM | BL | 34 | NDAM BL 34 - NDAM BL 00 |
| NDAM LI 12 | NDAM | LI | 12 | NDAM LI 12 - NDAM LI 00 |
| NDAM LI 15 | NDAM | LI | 15 | NDAM LI 15 - NDAM LI 00 |
| NDAM LI 18 | NDAM | LI | 18 | NDAM LI 18 - NDAM LI 00 |
| NDAM LI 21 | NDAM | LI | 21 | NDAM LI 21 - NDAM LI 00 |
| NDAM LI 26 | NDAM | LI | 26 | NDAM LI 26 - NDAM LI 00 |
| NDAM LI 29 | NDAM | LI | 29 | NDAM LI 29 - NDAM LI 00 |
| NDAM LI 32 | NDAM | LI | 32 | NDAM LI 32 - NDAM LI 00 |
| NDAM LI 35 | NDAM | LI | 35 | NDAM LI 35 - NDAM LI 00 |
| NDAM LN 21 | NDAM | LN | 21 | NDAM LN 21 - NDAM LN 00 |
| NDAM LN 35 | NDAM | LN | 35 | NDAM LN 35 - NDAM LN 00 |
| NDAM SP 21 | NDAM | SP | 21 | NDAM SP 21 - NDAM SP 00 |
| NDAM SP 35 | NDAM | SP | 35 | NDAM SP 35 - NDAM SP 00 |
| BORA BL 14 | BORA | BL | 14 | BORA BL 14 - BORA BL 00 |
| BORA BL 25 | BORA | BL | 25 | BORA BL 25 - BORA BL 00 |
| BORA BL 34 | BORA | BL | 34 | BORA BL 34 - BORA BL 00 |
| BORA LI 12 | BORA | LI | 12 | BORA LI 12 - BORA LI 00 |
| BORA LI 15 | BORA | LI | 15 | BORA LI 15 - BORA LI 00 |
| BORA LI 18 | BORA | LI | 18 | BORA LI 18 - BORA LI 00 |
| BORA LI 21 | BORA | LI | 21 | BORA LI 21 - BORA LI 00 |
| BORA LI 26 | BORA | LI | 26 | BORA LI 26 - BORA LI 00 |
| BORA LI 29 | BORA | LI | 29 | BORA LI 29 - BORA LI 00 |
| BORA LI 32 | BORA | LI | 32 | BORA LI 32 - BORA LI 00 |
| BORA LI 35 | BORA | LI | 35 | BORA LI 35 - BORA LI 00 |
| BORA LN 21 | BORA | LN | 21 | BORA LN 21 - BORA LN 00 |
| BORA LN 35 | BORA | LN | 35 | BORA LN 35 - BORA LN 00 |
| BORA SP 21 | BORA | SP | 21 | BORA SP 21 - BORA SP 00 |
| BORA SP 35 | BORA | SP | 35 | BORA SP 35 - BORA SP 00 |

**Table S3.** Tissue, days post infection (dpi), and the top 10 most significant genes with increased and decreased expression with valid gene symbols for the direct contrasts.

| Tissue | dpi | Genes with increased expression |
|---|---|---|
| BL | 0 | *TTLL1, PGA5, USP42, OLFM4, GPR132, MED27, WLS, SH3GLB2, PMF1, EIF2B2* |
| BL | 14 | *TTLL1, USP42, GPR132, UPK3B, TM9SF5, WLS, KLC1, MRPS6, TIPIN, BLK* |
| BL | 25 | *TTLL1, TM9SF5, TBXA2R, ADRA2A, RUM1, DHPS, ARL6IP4, ZNF318, KLHDC8A, APOL3* |
| BL | 34 | *TTLL1, SPDEF, KLHDC8A, TM9SF5, DHPS, KCNN4, RUM1, ARL6IP4, CDK20, FCRL3* |
| LI | 0 | *DOCK11, MAGIX, AGA, MEP1B, TMLHE, TCIRG1, SETD9, IRAK4, TRIM13, GSTM1* |
| LI | 12 | *DOCK11, AGA, MAGIX, CYP4F47, SETD9, IRAK4, IL18BP, TCIRG1, CHIA, TMEM41A* |
| LI | 15 | *MAGIX, DOCK11, ALDH7A1, AGA, TMLHE, CYP4A59, CYP4A11, INTS8, PON3, CYP4F47* |
| LI | 18 | *DOCK11, MAGIX, AGA, EPCAM, PDLIM4, ARL4D, ADTRP, TTLL1, SETD9, LTF* |
| LI | 21 | *AGA, CYP2D43, CYP2D14, COL12A1, EPPK1, DOCK11, MAGIX, TRIM13, IRAK4, ALDH7A1* |
| LI | 26 | *UGT2B10, DOCK11, TSPAN6, CCDC191, MAGIX, RTN2, SETD9, AGA, MMAA, DBI* |
| LI | 29 | *UGT2B10, DOCK11, CYP4A59, CYP4A11, AGA, DPYS, ATPAF1, QPRT, CYP4F47, MAGIX* |
| LI | 32 | *DOCK11, AGA, ADTRP, LTF, MAGIX, UGT2B10, APLNR, bta-mir-30f, GSTM1, EPCAM* |
| LI | 35 | *DOCK11, AGA, CYP2D43, CYP2D14, CYP4F47, CHIA, TSPAN6, PTPRM, RETSAT, CTDSP2* |
| LN | 0 | *CLDN11, TTLL1, BRB, CYP4B1, STAB1, PROS1, HEPH, CTSK, C1QTNF5, MT3* |
| LN | 21 | *TTLL1, SLC38A11, SUCLG1, SNORA73, EIF2B2, TXNDC11, MZB1, SLC25A26, PDIA5, ORC1* |
| LN | 35 | *TTLL1, SNORA73, LXN, MYB, DEFB4A, SLC38A11, CHD1L, TRNAU1AP, bta-mir-221, ORC1* |
| SP | 0 | *TTLL1, F2RL2, FABP3, RGS5, KIF3A, BMP2, TNFAIP6, NDN, SYT1, ITGA8* |
| SP | 21 | *TTLL1, ECRG4, TRMT10B, PTGDS, STMN2, KCNA3, CD38, CD2, CFI, MSN* |
| SP | 35 | *TTLL1, SNCA, LXN, RNASE4, MSN, ECRG4, STMN2, MAB21L1, TIPIN, TM9SF5* |
| **Tissue** | **dpi** | **Genes with decreased expression** |
| BL | 0 | *PHF12, VCAN, S100A7, PPP1R15A, CXCL5, ADAMDEC1, PLAU, CLECL1P, SAP18, CRYGS* |
| BL | 14 | *PHF12, GABARAPL2, OASL, MORN4, CRYGS, UCHL3, SEL1L3, MED16, SAP18, ERAP1* |
| BL | 25 | *VCAN, PFKFB4, PROCR, MORN4, JAM3, SELENOP, PHF12, DHRS7, SLC25A17, MTURN* |
| BL | 34 | *PROCR, MTURN, SLC25A17, MORN4, DHRS7, SMOX, TUBB1, ISCU, MFSD2B, ITGA2* |

**Table S3 continued.**

| Tissue | dpi | Genes with decreased expression |
|---|---|---|
| LI | 0 | *RFTN1, CYP4F2, TMEM45B, TPD52L1, MANEA, SPTSSB, PSMC5, PLBD1, SAP18, MED16* |
| LI | 12 | *SPTSSB, RFTN1, CYP4F2, MANEA, DNAJC22, TPD52L1, TMEM45B, TMLHE, BAMBI, TBC1D7* |
| LI | 15 | *SPTSSB, CYP4F2, RFTN1, PLBD1, TMEM45B, RAC1, TPD52L1, MANEA, ERAP1, GPX1* |
| LI | 18 | *CYP4F2, BAMBI, TPD52L1, RFTN1, SPTSSB, DNAJC22, PTER, MANEA, DNASE1L3, DCXR* |
| LI | 21 | *TPD52L1, PTER, KBTBD6, CYP4F2, MANEA, CRCP, RCBTB2, SLC5A6, TMLHE, JSP.1* |
| LI | 26 | *RFTN1, CYP4F2, ARSB, COX10, MANEA, ARPC1B, PAMR1, TMLHE, LCP1, NAGK* |
| LI | 29 | *JCHAIN, WWOX, PEG3, RFTN1, DNMT1, TPD52L1, PTER, ARSB, UBR7, OTULINL* |
| LI | 32 | *RFTN1, TPD52L1, CYP4F2, MANEA, PTER, SLC38A11, PIGR, JCHAIN, VPS41, HTATSF1* |
| LI | 35 | *RFTN1, TPD52L1, PARP11, TMLHE, MANEA, AVPR1A, CRCP, TBC1D7, C29H11orf86, JCHAIN* |
| LN | 0 | *SELE, CCL20, SEL1L3, EFHD1, ATP4B, SPP1, CXCL13, ALB, PLOD2, MME* |
| LN | 21 | *FOXO1, SEL1L3, COL16A1, ADAMDEC1, DDT, MOSMO, ATP4B, SELE, EFHD1, LTBP2* |
| LN | 35 | *LTBP2, DNER, IGFBP3, FOXO1, EFEMP1, RARRES2, DNASE1L3, DEFB10, VPS41, TBC1D7* |
| SP | 0 | *CFH, CYP4B1, EFHD1, LTBP2, EXOC7, FAM83D, MCPH1, TM4SF18, DDT, HCLS1* |
| SP | 21 | *CFH, FOXO1, DDT, SCG2, THBS1, BCAT2, PTER, RARRES2, MORN4, C6* |
| SP | 35 | *NELL2, CFH, RARRES2, SCG2, GDPD2, PLAU, SAP18, JSP.1, CD14, FOXO1* |

**Table S4.** Tissue, days post infection (dpi), and the top 10 most significant genes with increased and decreased expression with valid gene symbols for the N'Dama contrasts.

| Tissue | dpi | Genes with increased expression |
|---|---|---|
| BL | 14 | *SRSF11, HNRNPH1, UPK3B, CEP95, LAP3, NUPR1, PRPF40A, CCAR1, SNORD24, ABCC10* |
| BL | 25 | *NEB, ADORA2B, CMTM3, CPLANE1, CERS4, UPK3B, RHBDF2, ADRA2A, SRSF11, KCNN4* |
| BL | 34 | *NEB, CERS4, ADRA2A, UPK3B, FCRL3, KCNN4, WDR73, CMTM3, RHBDF2, LCAT* |
| LI | 12 | *CXCL11, UBD, TMSB10, WARS1, TAP1, PSMB9, BCL2A1, IRF1, BIRC3, PSMB10* |
| LI | 15 | *TMSB10, ADA, CYRIB, CD48, HCK, SPI1, PTPRC, JCHAIN, SLA, TMSB4X* |
| LI | 18 | *TMSB10, CYRIB, PTPRC, SPI1, CD48, TMSB4X, ADA, HCK, ARHGDIB, CD53* |
| LI | 21 | *TMSB10, CYRIB, PTPRC, TMSB4X, ARHGDIB, SPI1, CD48, CD53, RAC2, ADA* |
| LI | 26 | *TMSB10, TMSB4X, CYRIB, PTPRC, SPI1, CD53, CD48, CCDC191, CD55, SLA* |
| LI | 29 | *TMSB10, TMSB4X, PTPRC, CYRIB, SPI1, CD53, CD55, CD48, TMSB4, HTRA4* |
| LI | 32 | *TMSB10, TMSB4X, CYRIB, PTPRC, CD53, SPI1, CD55, CD48, RAP1B, TMSB4* |
| LI | 35 | *TMSB10, TMSB4X, PTPRC, CYRIB, CD53, SPI1, ARHGDIB, SLA, CD48, RAP1B* |
| LN | 21 | *ORC1, MZB1, CDC6, SKA3, UBE2S, SPAG5, SLC25A5, SPDL1, NDUFA4, CDC20* |
| LN | 35 | *SPAG5, ORC1, MKI67, TOP2A, TPX2, BUB1, AURKB, CDC6, CENPT, ASPM* |
| SP | 21 | *IRF4, CDC6, SPAG5, KIF2C, ESPL1, MKI67, CENPT, H2AC18, H2AC19, KRTCAP2* |
| SP | 35 | *MKI67, ORC1, AURKB, SPAG5, IRF4, ESPL1, BUB1, CDC6, ASPM, UBE2C* |
| **Tissue** | **dpi** | **Genes with decreased expression** |
| BL | 14 | *VAT1L, OLFM4, JAML, PGA5, PMF1, GZMB, ALOX15, FCER1A, PTGDR2, RAVER1, ICAM3* |
| BL | 25 | *JAML, CSF3R, OLFM4, RAVER1, ICAM3, VAT1L, PIP5K1B, GZMB, NFAM1, PTGDR2* |
| BL | 34 | *JAML, PIP5K1B, VAT1L, RAVER1, ICAM3, OLFM4, GZMB, PTGDR2, CSF3R, FCER1A* |
| LI | 12 | *MAPK6, SLC25A15, NUDT12, EEF1A1, CRYZ, GLCE, EPB41L5, PHKB, MYO1B, GNA14* |
| LI | 15 | *SIGLEC1, IGFBP6, GSTA2, CDK3, TEN1, ACSM5, GALT, MAPK6, SLC51B, JOSD2* |
| LI | 18 | *SIGLEC1, GCAT, AS3MT, PPOX, SLC51B, PMVK, IGFBP6, EPB41L5, NTN5, RORC* |
| LI | 21 | *RORC, GCAT, FAM83H, SIGLEC1, C15H11orf52, DVL1, CARD19, PTPRF, MOSPD1, NAA30* |
| LI | 26 | *SIGLEC1, GCAT, FOXA2, GUCD1, HPN, HOMER2, PTPRF, HSD17B14, THOP1, GPLD1* |

**Table S4 continued.**

| Tissue | dpi | Genes with decreased expression |
|---|---|---|
| LI | 29 | *SIGLEC1, GCAT, IYD, SHPK, NMRAL1, GRHPR, PPP2R2B, KHK, ACY1, FARP1* |
| LI | 32 | *GCAT, SIGLEC1, ACY1, FOXA2, MGC152281, PDK2, FAAH, GRHPR, GUCD1, SLC27A4* |
| LI | 35 | *SIGLEC1, NIT1, AS3MT, EPB41L5, GCAT, PDZK1, PLPP6, GSR, GRHPR, C15H11orf52* |
| LN | 21 | *CLDN11, ELOVL7, SNTB2, SMCHD1, BRB, SMAD5, CFLAR, NR2F1, ADGRF5, HEPH* |
| LN | 35 | *CLDN11, BRB, ELOVL7, SMYD2, HEPH, NR2F1, TJP1, SCARA5, MEIS2, ADGRF5* |
| SP | 21 | *SYT1, GDPD2, NTRK2, NTN4, IGFBP6, DNASE1L3, NDN, FGF2, PENK, F2RL2* |
| SP | 35 | *SYT1, GDPD2, IGFBP6, NTRK2, NTN4, NDN, FBLN5, PGM5, PENK, PYGM* |

**Table S5.** Tissue, days post infection (dpi), and the top 10 most significant genes with increased and decreased expression with valid gene symbols for the Boran contrasts.

| Tissue | dpi | Genes with increased expression |
|---|---|---|
| BL | 14 | *GZMB, LAP3, SLAMF8, IRF1, FCRL3, NUB1, NLRC5, OASL, WARS1, GBP5* |
| BL | 25 | *CPLANE1, CSTB, NEB, ADORA2B, CMTM3, SRSF11, PIK3C2B, SEC11A, INPP5B, UPK3B* |
| BL | 34 | *NEB, ADORA2B, CPLANE1, CMTM3, INPP5B, SLC25A12, CERS4, PIK3C2B, ADRA2A, PYGO1* |
| LI | 12 | *TAP1, UBD, IRF1, PSMB9, TMSB10, CXCL11, B2M, PSMB10, CXCL10, NLRC5* |
| LI | 15 | *TMSB10, CYRIB, ADA, CD48, SPI1, HCK, PTPRC, ARHGDIB, GNG2, CORO1A* |
| LI | 18 | *TMSB10, CYRIB, SPI1, PTPRC, CD48, ADA, ARHGDIB, HCK, PLTP, CD53* |
| LI | 21 | *TMSB10, CYRIB, CD53, SPI1, PTPRC, TMSB4X, ARHGDIB, ADA, CD48, ACTR3* |
| LI | 26 | *TMSB10, CYRIB, PTPRC, SPI1, CD53, TMSB4X, PLTP, CDH5, HCK, ARHGDIB* |
| LI | 29 | *CYRIB, TMSB10, SPI1, PTPRC, CD53, PLTP, TMSB4X, RAP1B, ARHGDIB, DOCK2* |
| LI | 32 | *TMSB10, CYRIB, PTPRC, SPI1, CD53, TMSB4X, PLTP, CD55, HCK, RAP1B* |
| LI | 35 | *TMSB10, CYRIB, CD53, PTPRC, SPI1, TMSB4X, PLTP, RAP1B, CD48, VAV1* |
| LN | 21 | *MEA1, ATP5ME, MRPL52, POLR2L, RNASEH2C, SLC25A5, MZB1, COX6B1, TXNDC5, FAM32A* |
| LN | 35 | *MEA1, NDUFA4, COX6B1, KIFC1, SPAG5, ATP5ME, AQP3, MKI67, TXNDC5, TOP2A* |
| SP | 21 | *CDC6, IRF4, KRTCAP2, RNASEH2C, CSTB, MYDGF, BUB1, TUBG1, CENPT, ESPL1* |
| SP | 35 | *CSTB, LAP, IRF4, CD14, CTLA4, CDC6, MYDGF, TUBG1, ACSL5, NOLC1* |
| **Tissue** | **dpi** | **Genes with decreased expression** |
| BL | 14 | *PIP5K1B, ADAMDEC1, IFT27, CA5B, S100A7, FCER1A, RGCC, JAML, FOSB, NR4A2* |
| BL | 25 | *JAML, PIP5K1B, CXCR1, CXCR2, IFT27, GZMB, ALOX15, CSF3R, VAT1L, PTGDR2* |
| BL | 34 | *JAML, IFT27, PIP5K1B, CA5B, S100A7, PARP8, CSF3R, CXCL8, NFAM1, PTGDR2* |
| LI | 12 | *RPS2, SNORA64, CA5A, TRIM6, PAF1, FGA, HSD17B14, 7SK, SLC7A9, HEXIM2* |
| LI | 15 | *COLEC11, TCEA3, EPB41L5, SIGLEC1, IMMP2L, PYURF, AS3MT, AUH, APOM, AGMAT* |
| LI | 18 | *SIGLEC1, PTPRF, PMVK, TEAD2, APLNR, TCEA3, DHRS11, EPB41L5, ANKS4B, CBS* |
| LI | 21 | *PHACTR4, APLNR, TCEA3, RORC, GCGR, GCAT, SIGLEC1, FAM149A, ST7L, CRYBG2* |
| LI | 26 | *PMVK, SIGLEC1, GCAT, CBS, HSD17B14, PTPRF, ACY1, HAGH, CIDEA, PAFAH2* |

**Table S5 continued.**

| Tissue | dpi | Genes with decreased expression |
|---|---|---|
| LI | 29 | *GCAT, GRHPR, FAAH, EPB41L5, DPYS, PMVK, CBS, NIPSNAP1, TCEA3, ACY1* |
| LI | 32 | *GCAT, PMVK, CES1, SIGLEC1, KCTD21, ACY1, FAAH, EBP, FOXA2, MMAB* |
| LI | 35 | *GCAT, GCGR, EBP, PMVK, ACY1, GRHPR, PTPRF, FAH, NTN5, HPN* |
| LN | 21 | *MSANTD2, MFSD4A, SMCHD1, NPNT, SHISA3, ZNF318, MDFIC, BPTF, KMT2C, TSC1* |
| LN | 35 | *MSANTD2, LUM, CLDN11, SMCHD1, MFSD4A, KMT2C, MINDY2, COL6A3, NR2F1, AGO3* |
| SP | 21 | *SYT1, NTN4, DNASE1L3, GDPD2, IGFBP6, NTRK2, PGM5, NXPH1, FBLN5, GPR34* |
| SP | 35 | *NTRK2, IGFBP6, GDPD2, SYT1, NTN4, FBLN5, PGM5, PYGM, DNASE1L3, TPM2* |

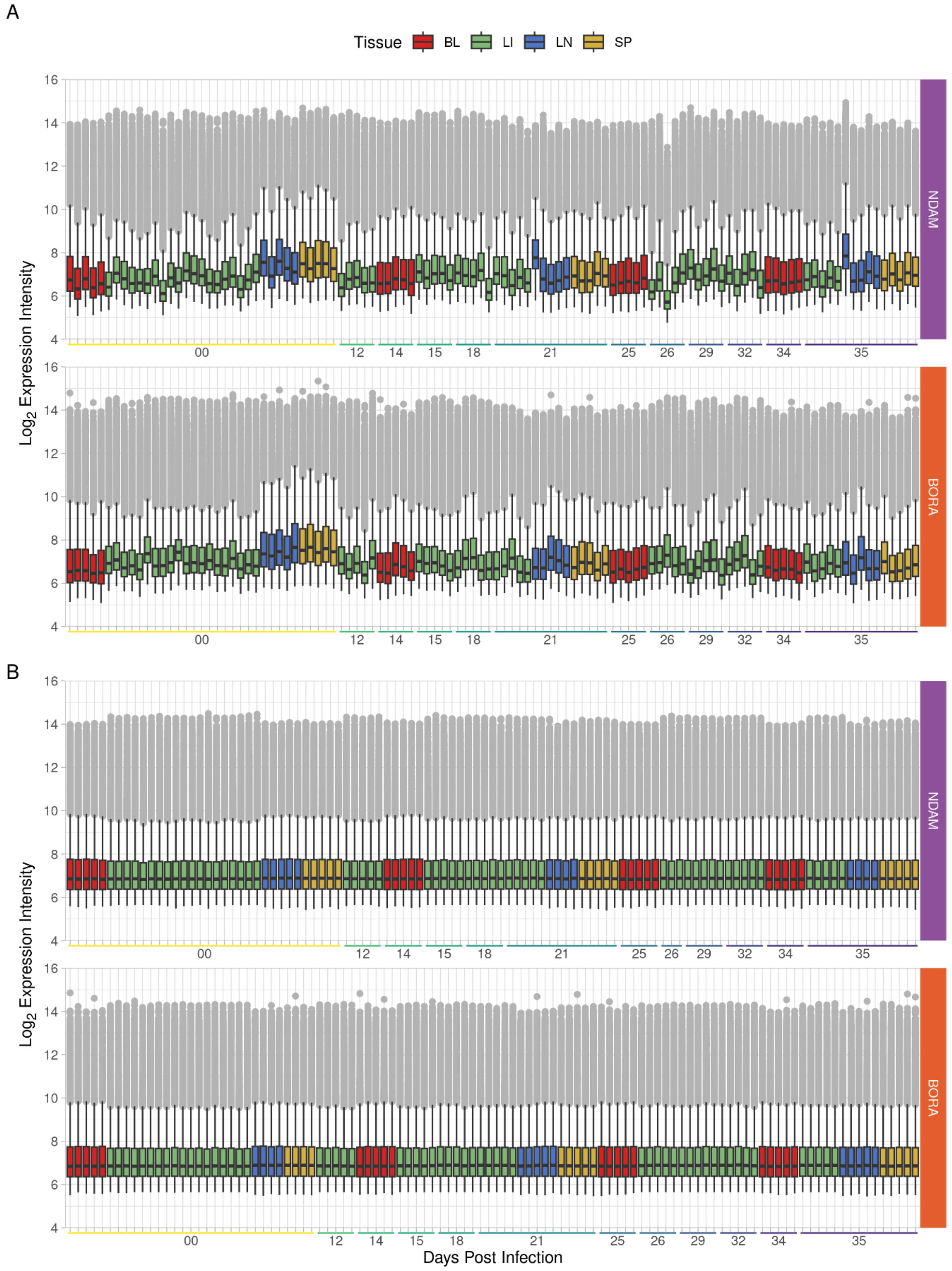


**Fig S1.** Boxplots showing the $\log_2$ expression intensity of the probe sets for each sample of the **A.** raw and **B.** normalised data after quality control filtering separated into NDAM and BORA populations and coloured according to tissue. The position along the horizontal axis indicates the days post infection. Outlier probe sets are shown as grey dots.

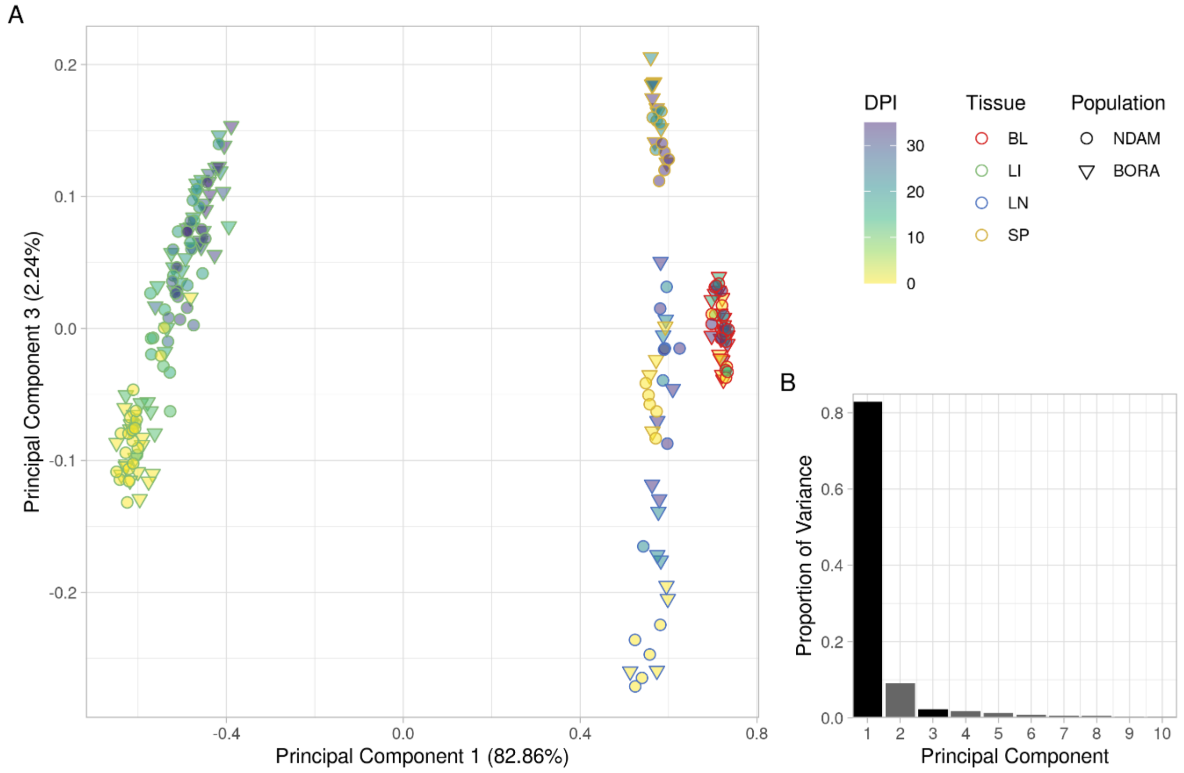


**Fig S2. A.** Principal component analysis (PCA) of the microarray data set with samples coloured according to days post infection (dpi) with the outer colour representing the tissue and shape indicating the population showing the first and third principal components (PC1 and PC3), and **B.** bar chart of proportion of variance of the top ten PCs.

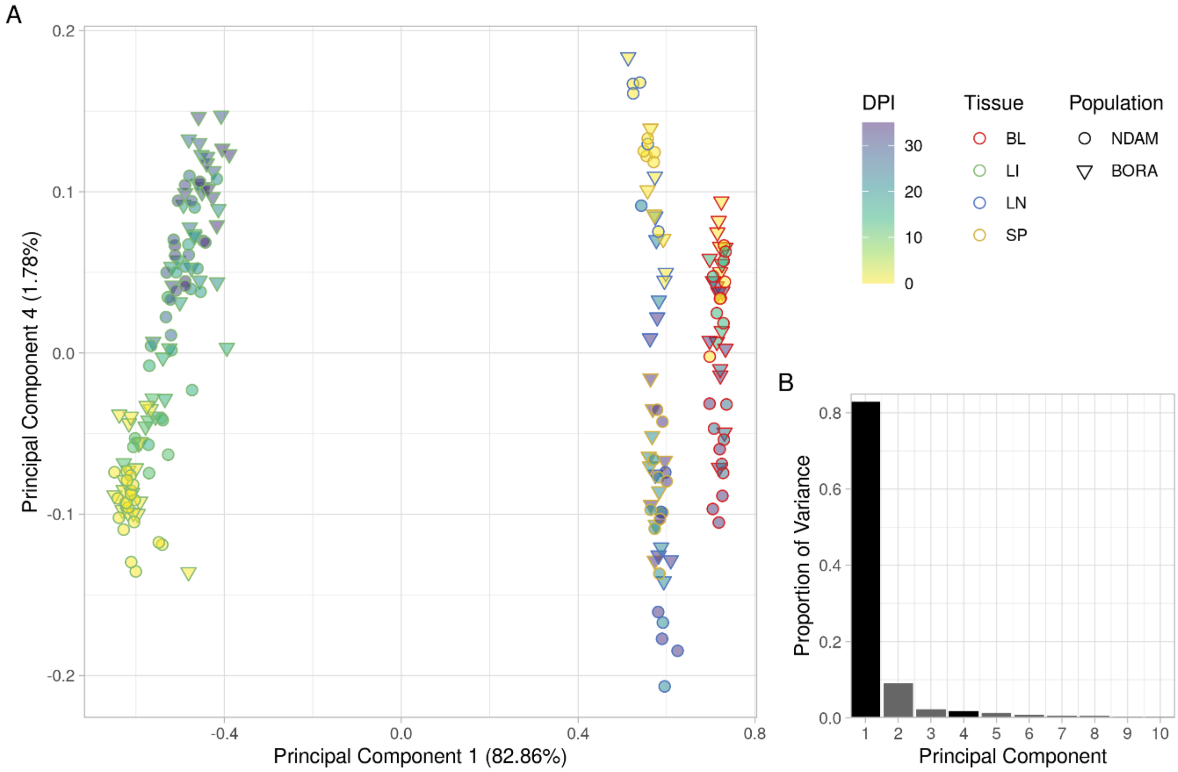


**Fig S3. A.** Principal component analysis (PCA) of the microarray data set with samples coloured according to days post infection (dpi) with the outer colour representing the tissue and shape indicating the population showing the first and fourth principal components (PC1 and PC4), and **B.** bar chart of proportion of variance of the top ten PCs.

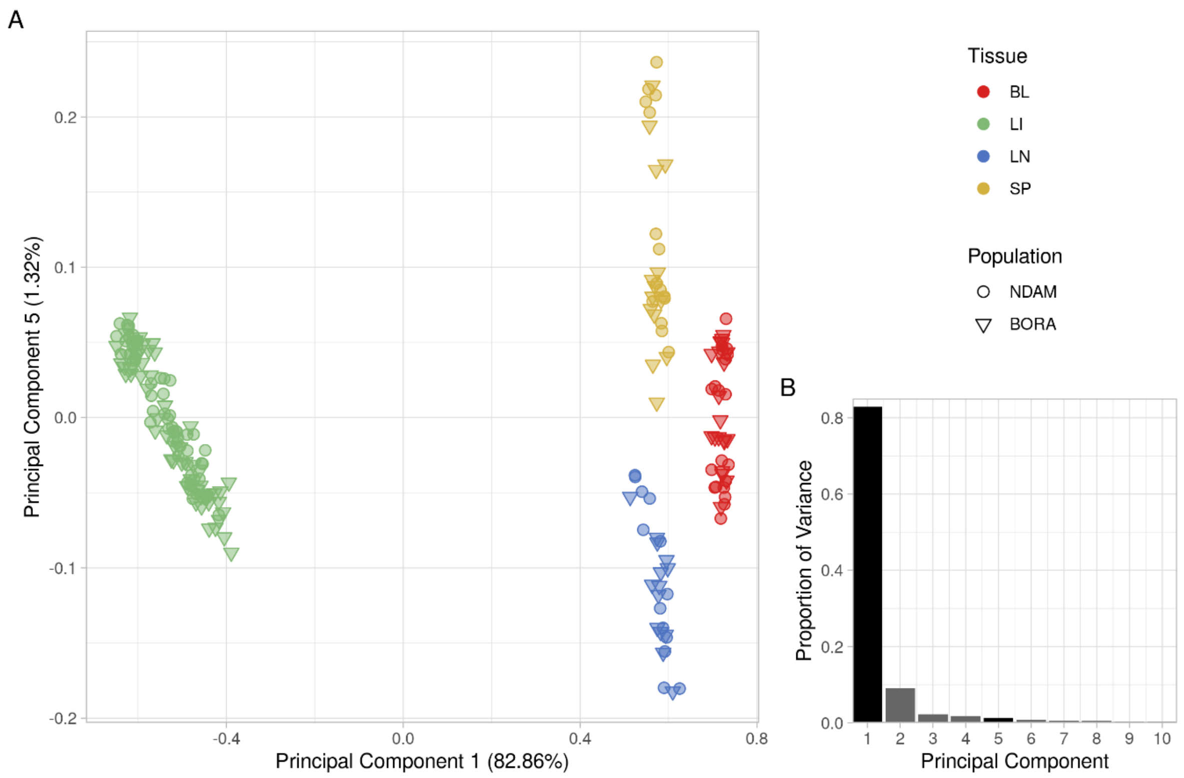


**Fig S4. A.** Principal component analysis (PCA) of the microarray data set with samples coloured according to tissue and shape indicating the population showing the first and fifth principal components (PC1 and PC5), and **B.** bar chart of proportion of variance of the top ten PCs.

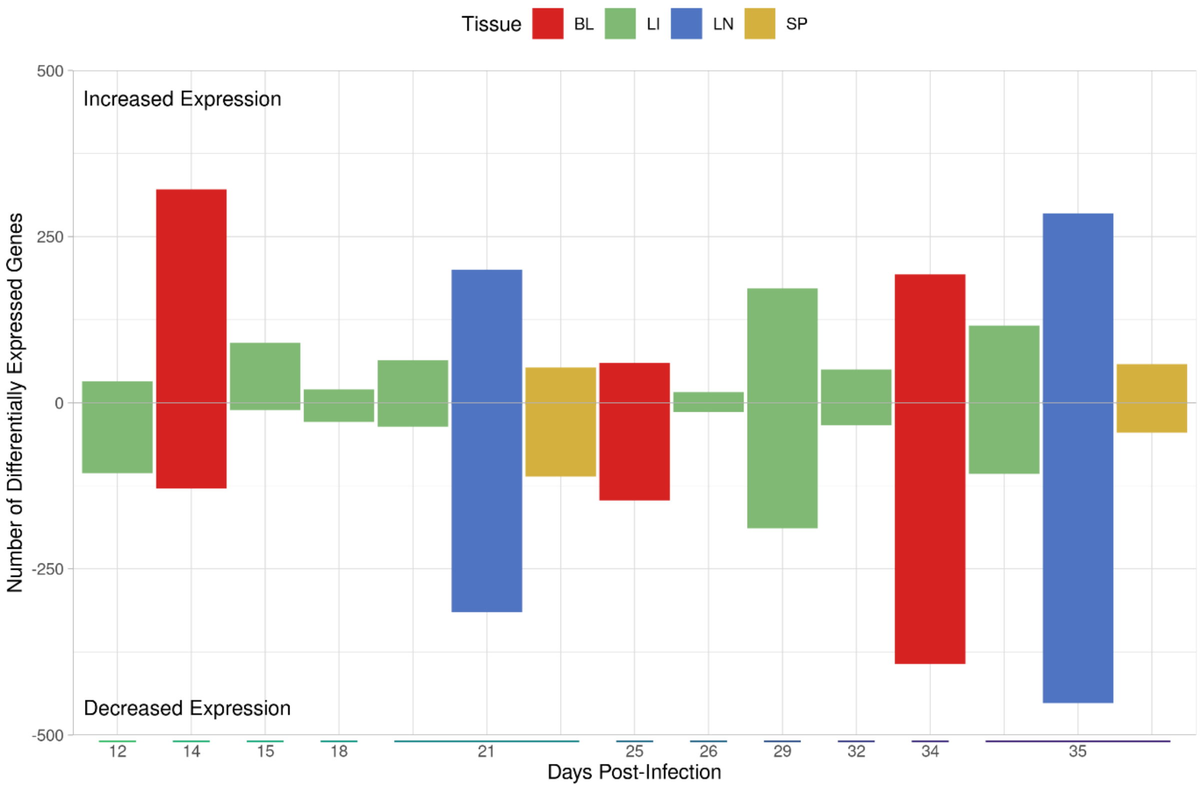


**Fig S5.** Bar chart showing the numbers of significantly differentially expressed genes for the RESP contrasts. The extent of the bar above and below 0 on the vertical axis indicates the numbers of significantly differentially expressed genes (DEGs) with increased and decreased expression, respectively. The position on the horizontal indicates the number of days post infection (dpi) and the colour of the bars represents the tissue.

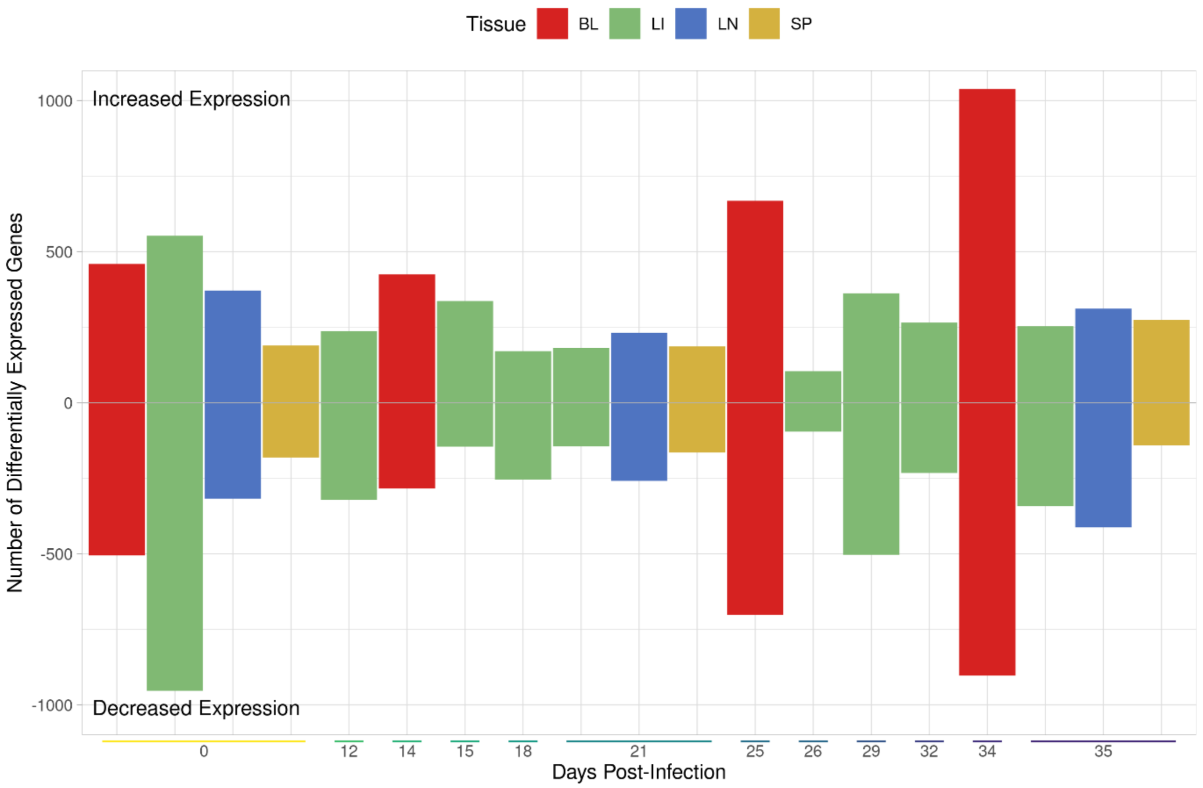


**Fig S6.** Bar chart showing the numbers of significantly differentially expressed genes for the DIRE contrasts. The extent of the bar above and below 0 on the vertical axis indicates the numbers of significantly differentially expressed genes (DEGs) with increased and decreased expression, respectively. The position on the horizontal indicates the number of days post infection (dpi) and the colour of the bars represents the tissue.

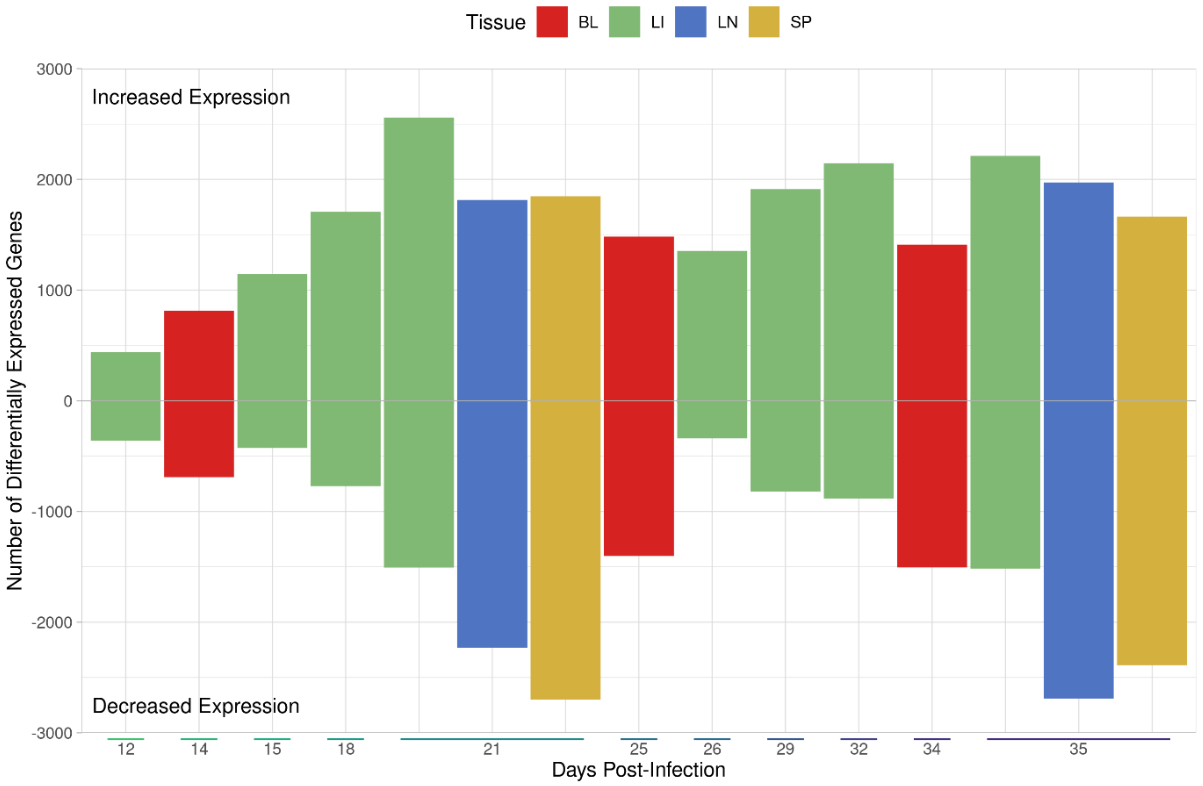


**Fig S7.** Bar chart showing the numbers of significantly differentially expressed genes for the NDAM contrasts. The extent of the bar above and below 0 on the vertical axis indicates the numbers of significantly differentially expressed genes (DEGs) with increased and decreased expression, respectively. The position on the horizontal indicates the number of days post infection (dpi) and the colour of the bars represents the tissue.

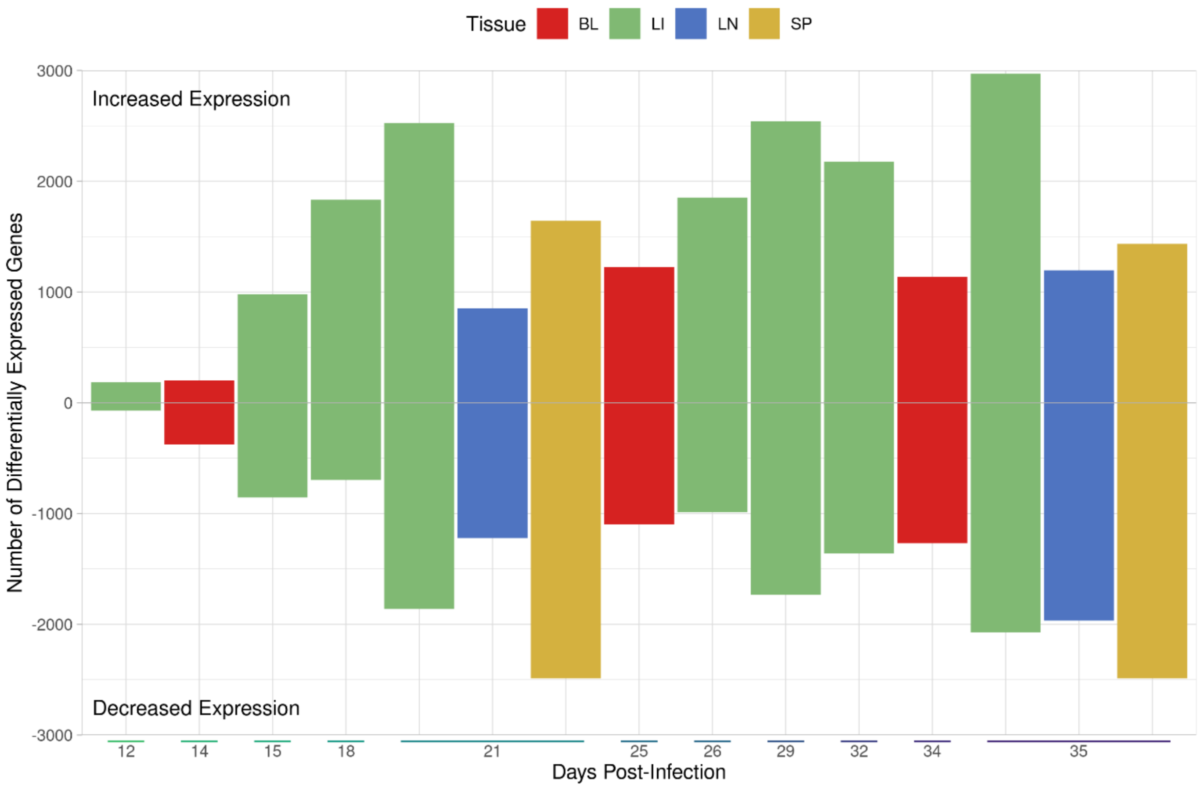


**Fig S8.** Bar chart showing the numbers of significantly differentially expressed genes for the BORA contrasts. The extent of the bar above and below 0 on the vertical axis indicates the numbers of significantly differentially expressed genes (DEGs) with increased and decreased expression, respectively. The position on the horizontal indicates the number of days post infection (dpi) and the colour of the bars represents the tissue.

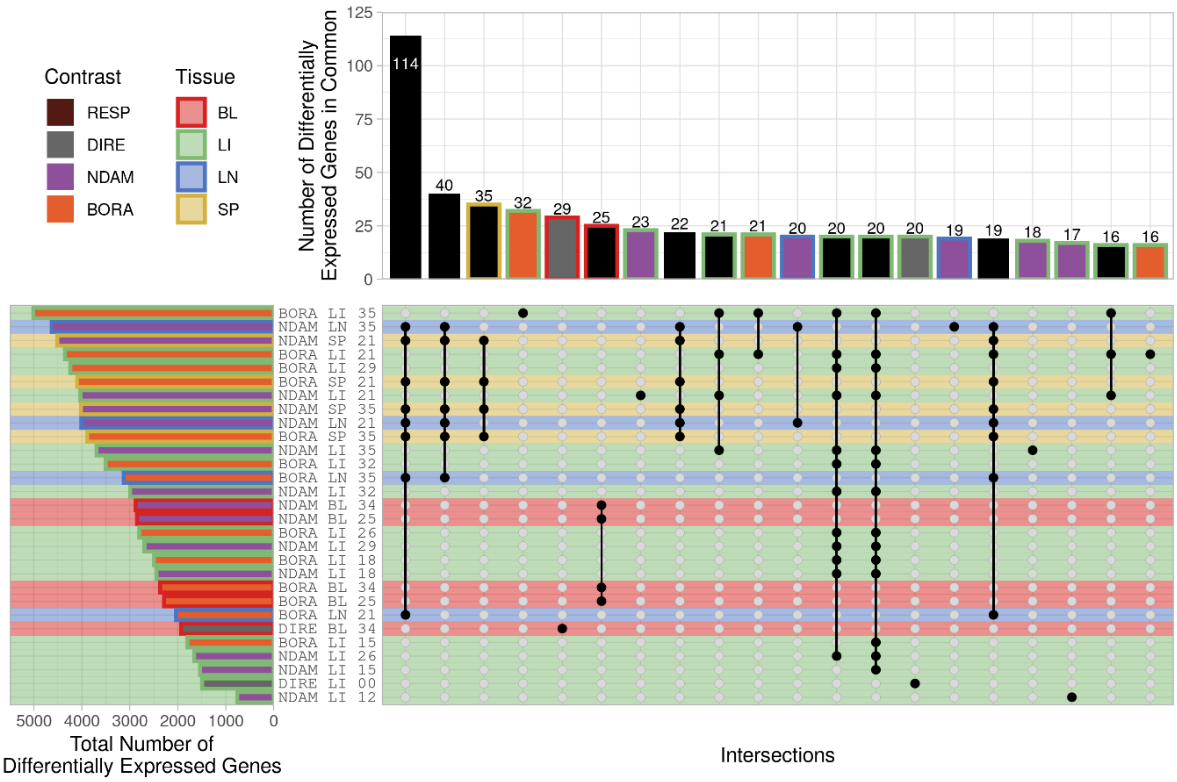


**Fig S9.** UpSet plot showing the top 20 intersections among all 64 contrasts. The horizontal bars indicate the total number of significant differentially expressed genes (DEGs) for each contrast while the vertical bars indicate the number of significant DEGs in common between the contrasts annotated with black dots connected by lines in the intersection matrix. The background colour of the stripes in the intersection matrix and horizontal bars represents the tissue. The colour of the bars represents the contrast type with black bars indicating an overlap between different contrast types. The outline colour of the bars also represents the tissue with no outline representing an overlap between different tissues.

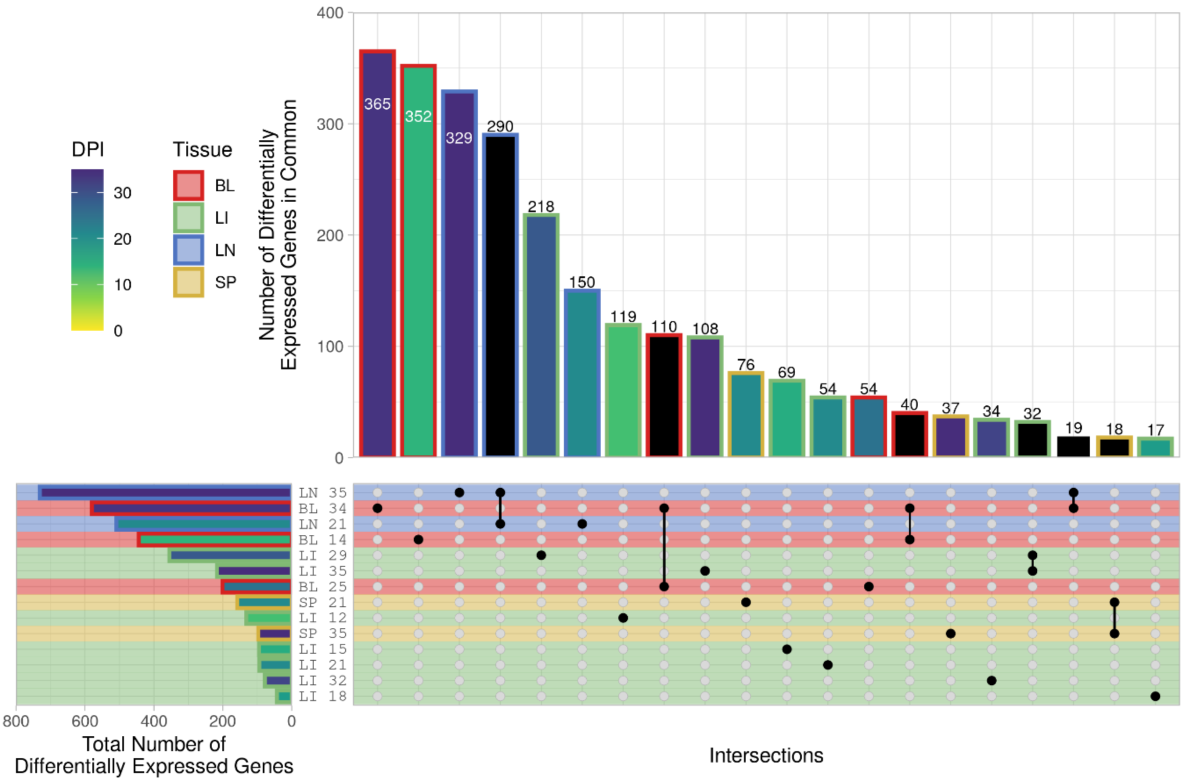


**Fig S10.** UpSet plot showing the top 20 intersections among the RESP contrasts. The horizontal bars indicate the total number of significant differentially expressed genes (DEGs) for each contrast while the vertical bars indicate the number of significant DEGs in common between the contrasts annotated with black dots connected by lines in the intersection matrix. The background colour of the stripes in the intersection matrix and horizontal bars represents the tissue. The colour of the bars represents the days post infection (dpi) with black bars indicating an overlap between different timepoints. The outline colour of the bars also represents the tissue with no outline representing an overlap between different tissues.

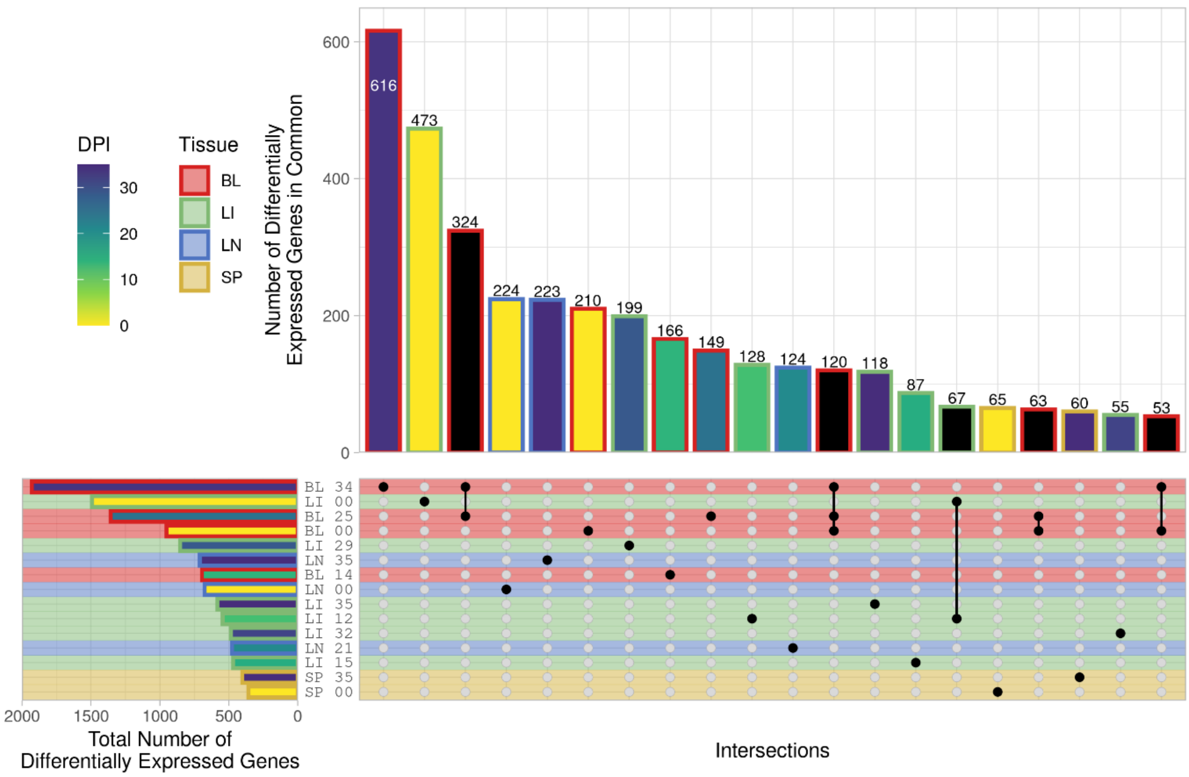


**Fig S11.** UpSet plot showing the top 20 intersections among the DIRE contrasts. The horizontal bars indicate the total number of significant differentially expressed genes (DEGs) for each contrast while the vertical bars indicate the number of significant DEGs in common between the contrasts annotated with black dots connected by lines in the intersection matrix. The background colour of the stripes in the intersection matrix and horizontal bars represents the tissue. The colour of the bars represents the days post infection (dpi) with black bars indicating an overlap between different timepoints. The outline colour of the bars also represents the tissue with no outline representing an overlap between different tissues.

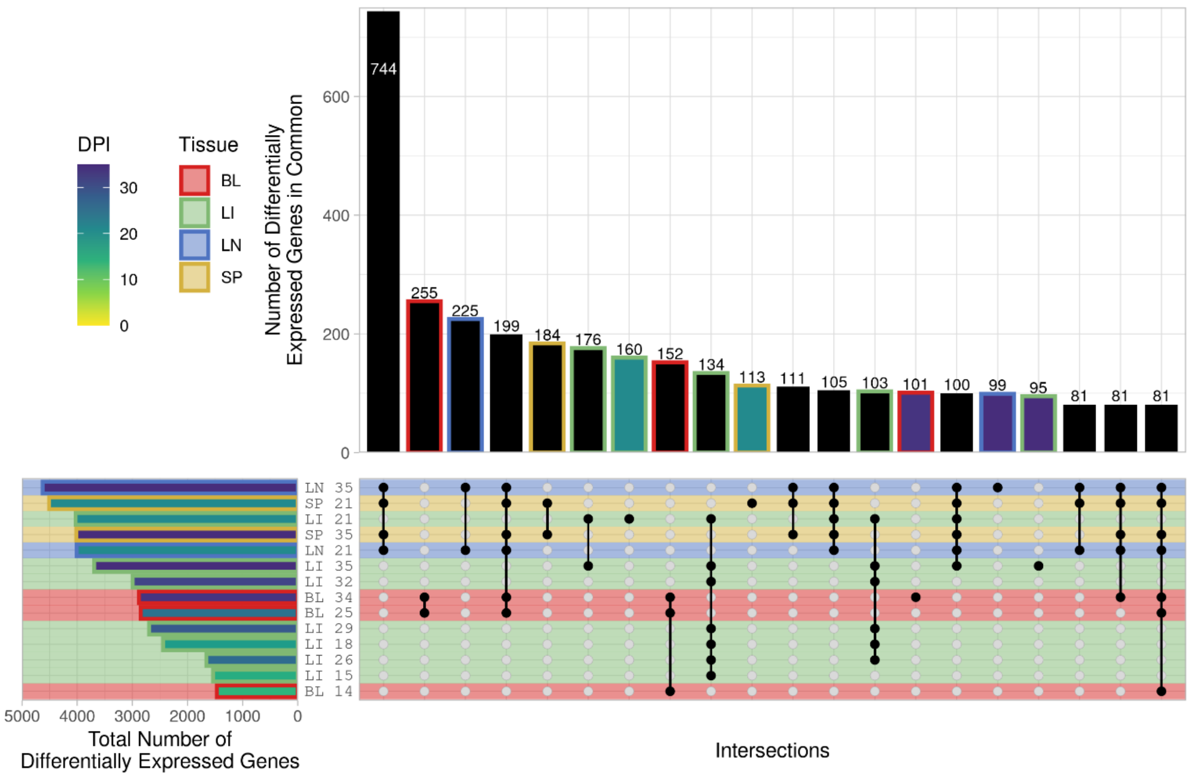


**Fig S12.** UpSet plot showing the top 20 intersections among the NDAM contrasts. The horizontal bars indicate the total number of significant differentially expressed genes (DEGs) for each contrast while the vertical bars indicate the number of significant DEGs in common between the contrasts annotated with black dots connected by lines in the intersection matrix. The background colour of the stripes in the intersection matrix and horizontal bars represents the tissue. The colour of the bars represents the days post infection (dpi) with black bars indicating an overlap between different timepoints. The outline colour of the bars also represents the tissue with no outline representing an overlap between different tissues.

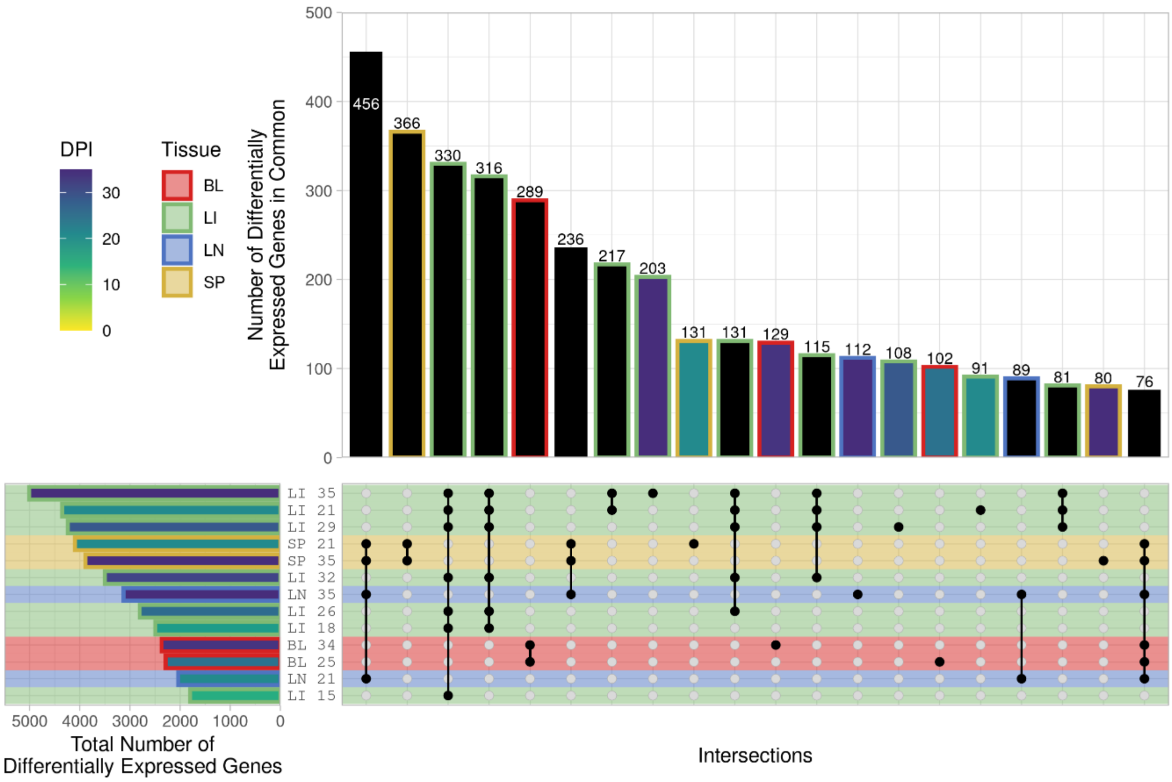


**Fig S13.** UpSet plot showing the top 20 intersections among the BORA contrasts. The horizontal bars indicate the total number of significant differentially expressed genes (DEGs) for each contrast while the vertical bars indicate the number of significant DEGs in common between the contrasts annotated with black dots connected by lines in the intersection matrix. The background colour of the stripes in the intersection matrix and horizontal bars represents the tissue. The colour of the bars represents the days post infection (dpi) with black bars indicating an overlap between different timepoints. The outline colour of the bars also represents the tissue with no outline representing an overlap between different tissues.

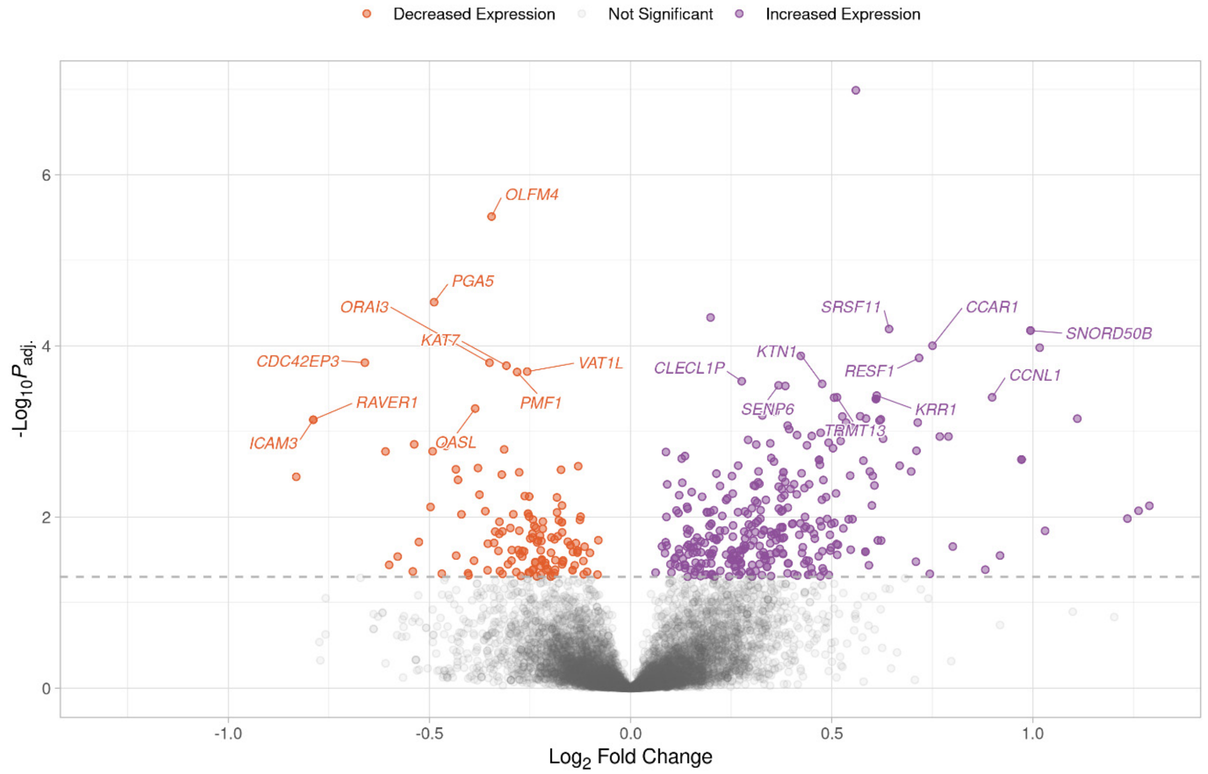


**Fig S14.** Volcano plot showing the results of the RESP contrast for the peripheral blood mononuclear cell (PBMC) samples at 14 days post infection (dpi). Each data point represents a gene with the position on the *x*- and *y*-axes indicating the $\log_2$ fold change and $-\log_{10}P_{adj.}$, respectively. Genes above the horizontal dashed line are significantly differentially expressed with the colours representing the change in expression. The top 10 most significant genes for increased and decreased expression with gene symbols are labelled.

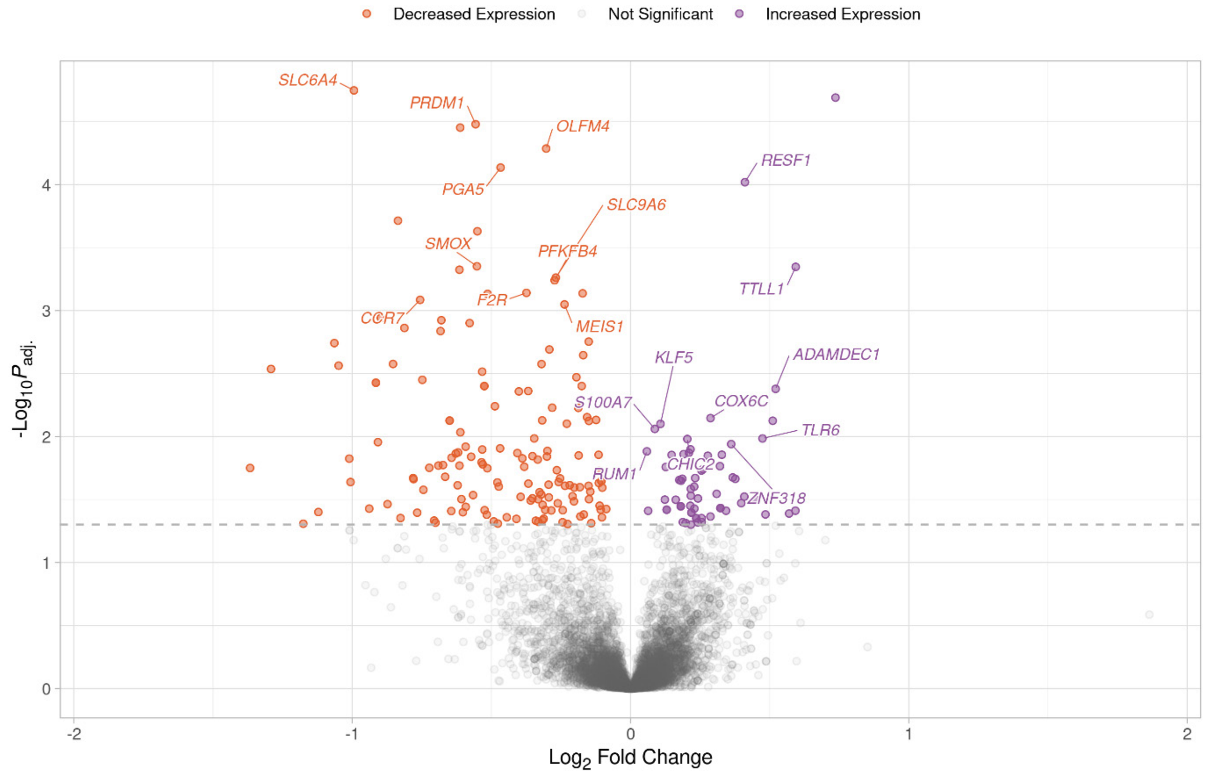


**Fig S15.** Volcano plot showing the results of the RESP contrast for the peripheral blood mononuclear cell (PBMC) samples at 25 days post infection (dpi). Each data point represents a gene with the position on the *x*- and *y*-axes indicating the $\log_2$ fold change and $-\log_{10}P_{adj.}$, respectively. Genes above the horizontal dashed line are significantly differentially expressed with the colours representing the change in expression. The top 10 most significant genes for increased and decreased expression with gene symbols are labelled.

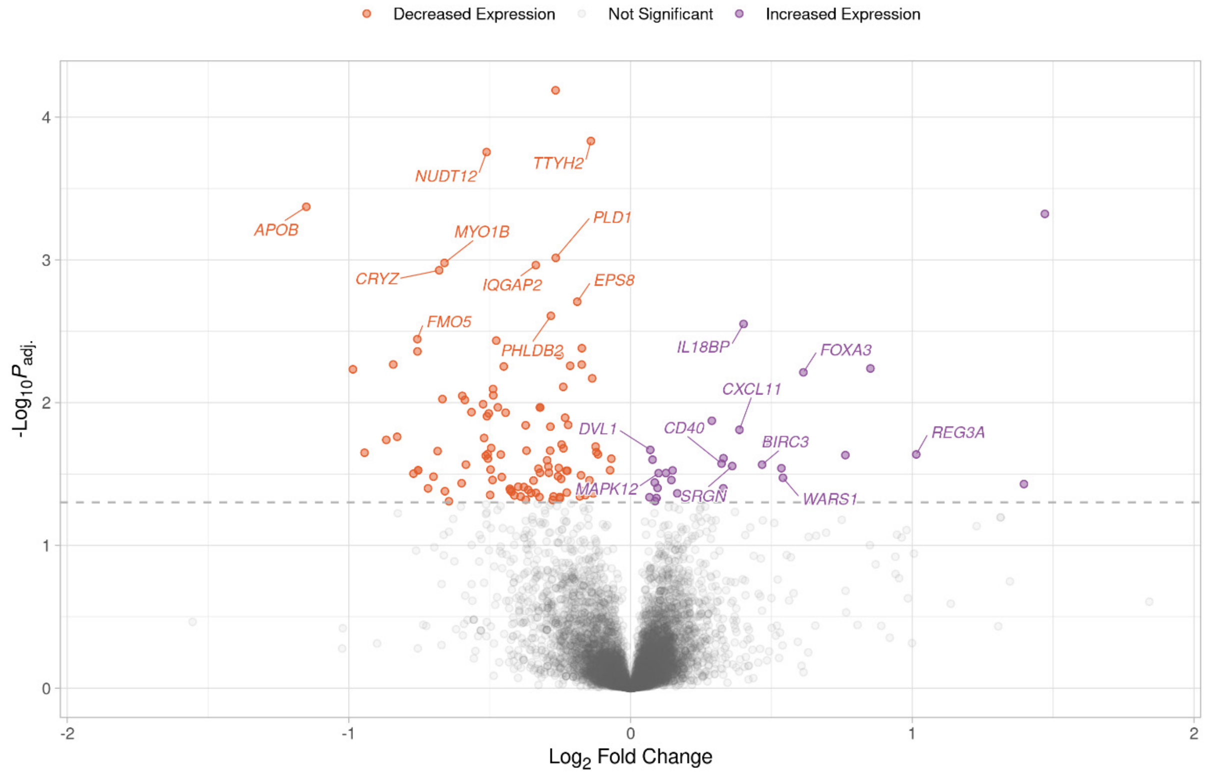


**Fig S16.** Volcano plot showing the results of the RESP contrast for the liver samples at 12 days post infection (dpi). Each data point represents a gene with the position on the *x*- and *y*-axes indicating the $\log_2$ fold change and $-\log_{10}P_{adj.}$, respectively. Genes above the horizontal dashed line are significantly differentially expressed with the colours representing the change in expression. The top 10 most significant genes for increased and decreased expression with gene symbols are labelled.

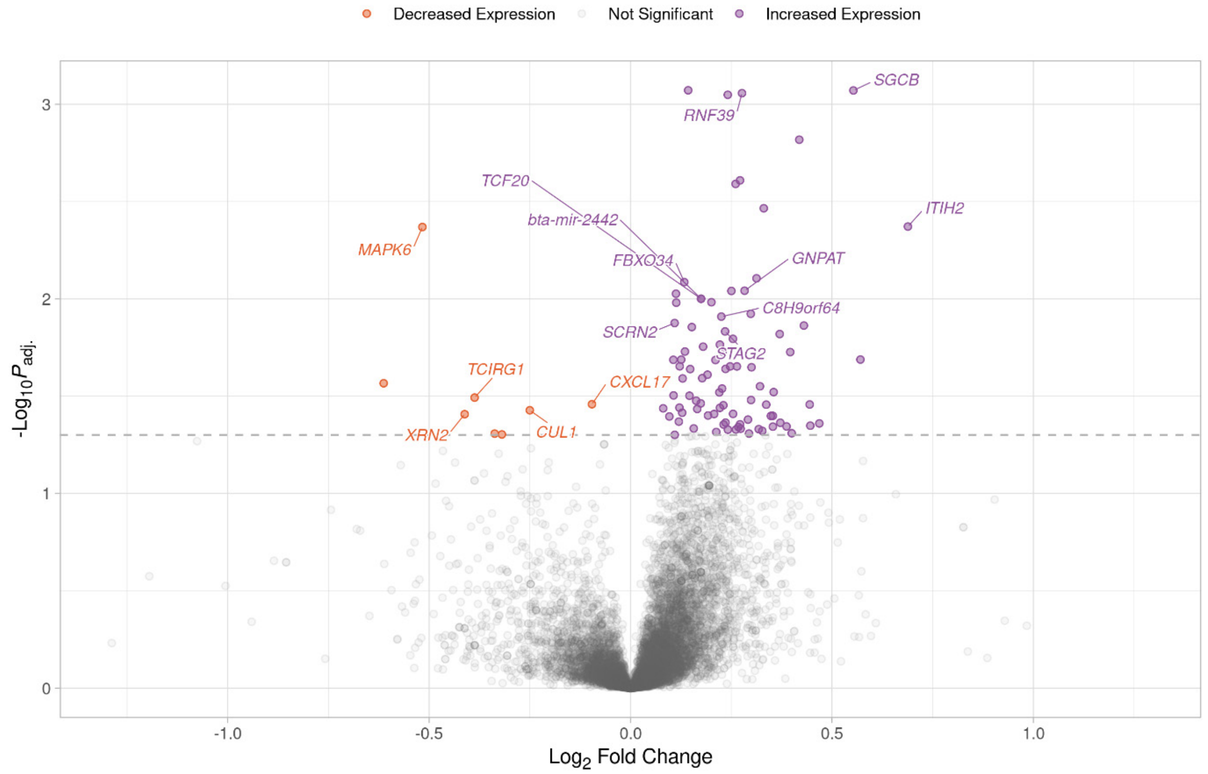


**Fig S17.** Volcano plot showing the results of the RESP contrast for the liver samples at 15 days post infection (dpi). Each data point represents a gene with the position on the *x*- and *y*-axes indicating the $\log_2$ fold change and $-\log_{10}P_{adj.}$, respectively. Genes above the horizontal dashed line are significantly differentially expressed with the colours representing the change in expression. The top 10 most significant genes for increased and decreased expression with gene symbols are labelled.

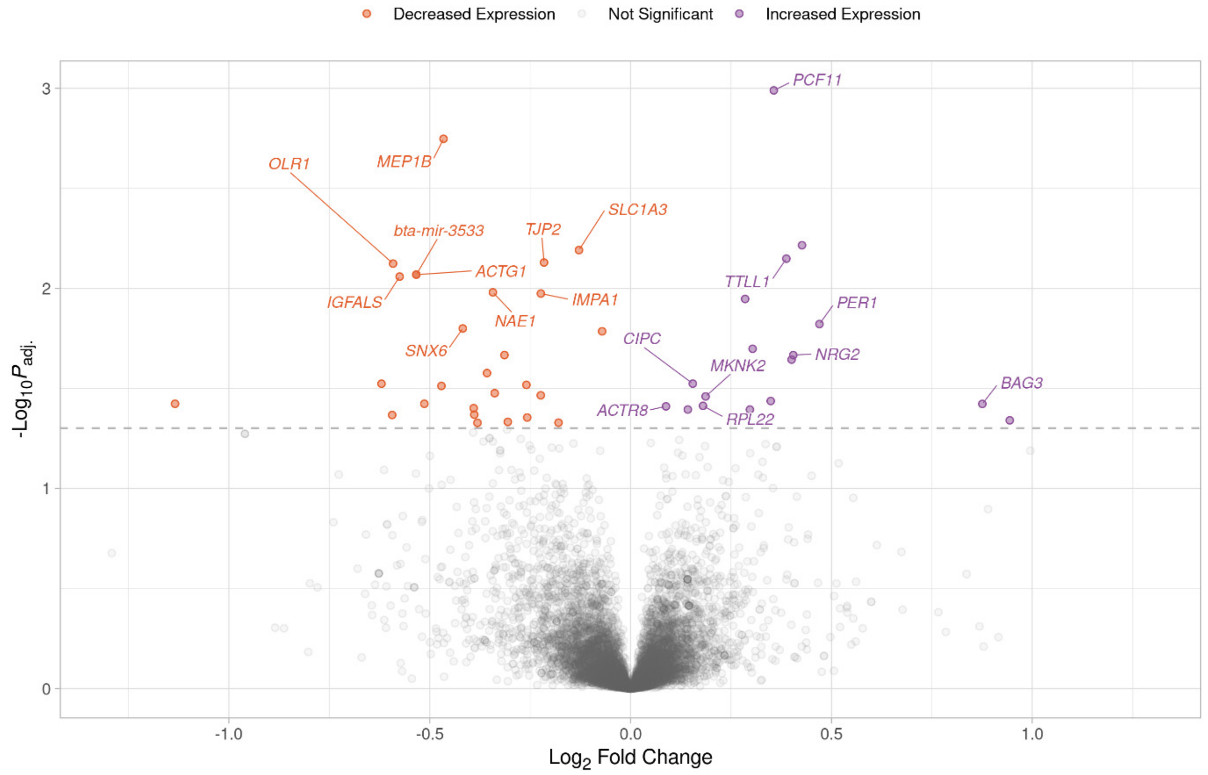


**Fig S18.** Volcano plot showing the results of the RESP contrast for the liver samples at 18 days post infection (dpi). Each data point represents a gene with the position on the *x*- and *y*-axes indicating the $\log_2$ fold change and $-\log_{10}P_{adj.}$, respectively. Genes above the horizontal dashed line are significantly differentially expressed with the colours representing the change in expression. The top 10 most significant genes for increased and decreased expression with gene symbols are labelled.

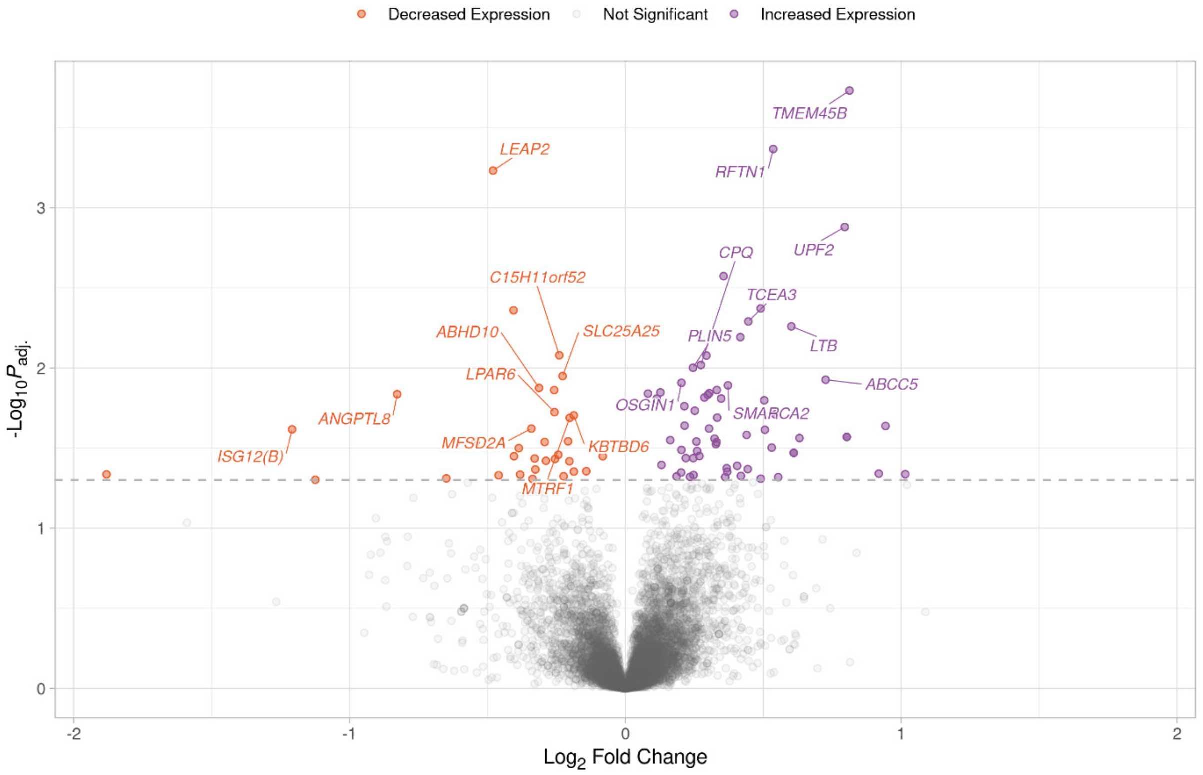


**Fig S19.** Volcano plot showing the results of the RESP contrast for the liver samples at 21 days post infection (dpi). Each data point represents a gene with the position on the *x*- and *y*-axes indicating the $\log_2$ fold change and $-\log_{10}P_{adj.}$, respectively. Genes above the horizontal dashed line are significantly differentially expressed with the colours representing the change in expression. The top 10 most significant genes for increased and decreased expression with gene symbols are labelled.

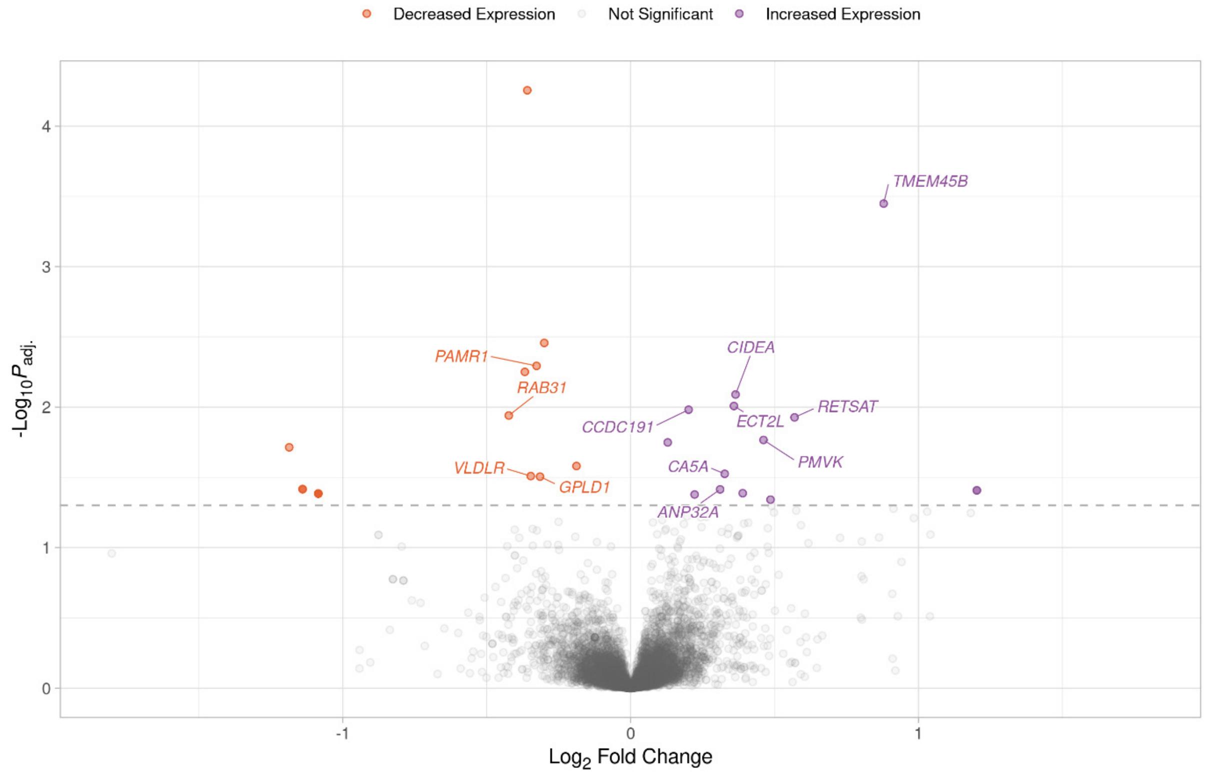


**Fig S20.** Volcano plot showing the results of the RESP contrast for the liver samples at 26 days post infection (dpi). Each data point represents a gene with the position on the *x*- and *y*-axes indicating the $\log_2$ fold change and $-\log_{10}P_{adj.}$, respectively. Genes above the horizontal dashed line are significantly differentially expressed with the colours representing the change in expression. The top 10 most significant genes for increased and decreased expression with gene symbols are labelled.

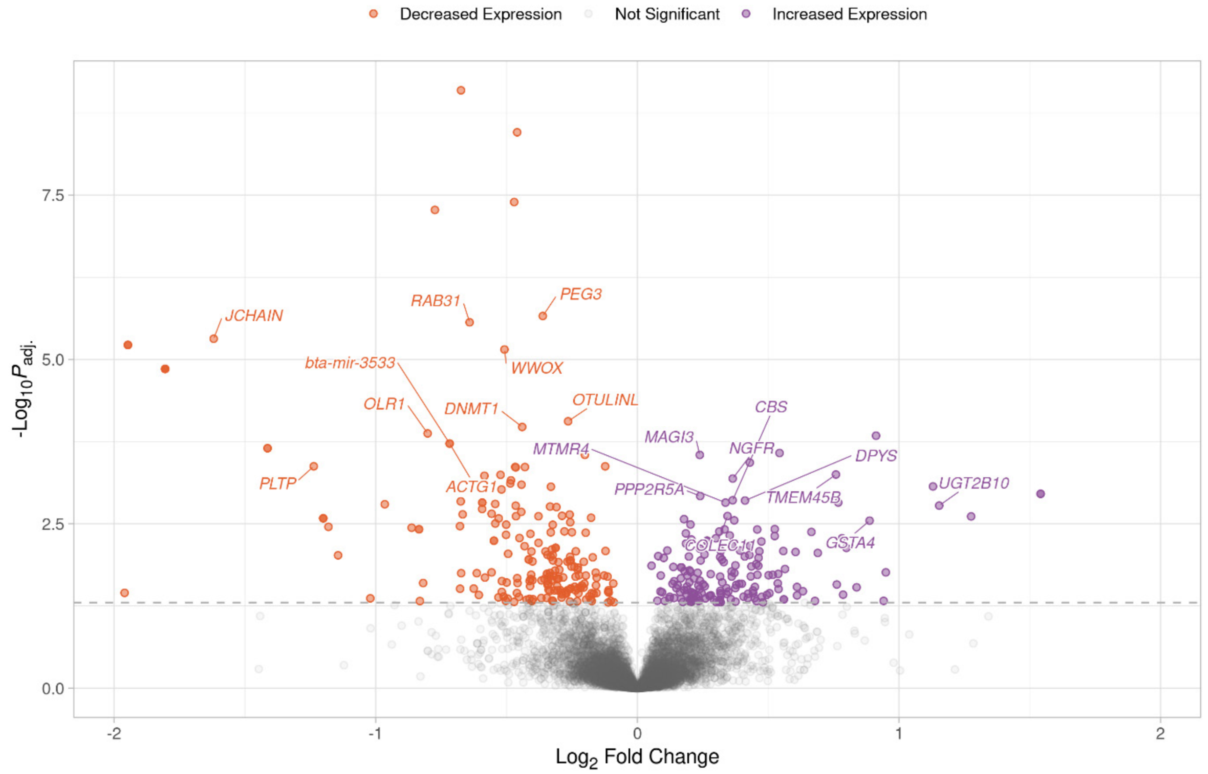


**Fig S21.** Volcano plot showing the results of the RESP contrast for the liver samples at 29 days post infection (dpi). Each data point represents a gene with the position on the *x*- and *y*-axes indicating the $\log_2$ fold change and $-\log_{10}P_{adj.}$, respectively. Genes above the horizontal dashed line are significantly differentially expressed with the colours representing the change in expression. The top 10 most significant genes for increased and decreased expression with gene symbols are labelled.

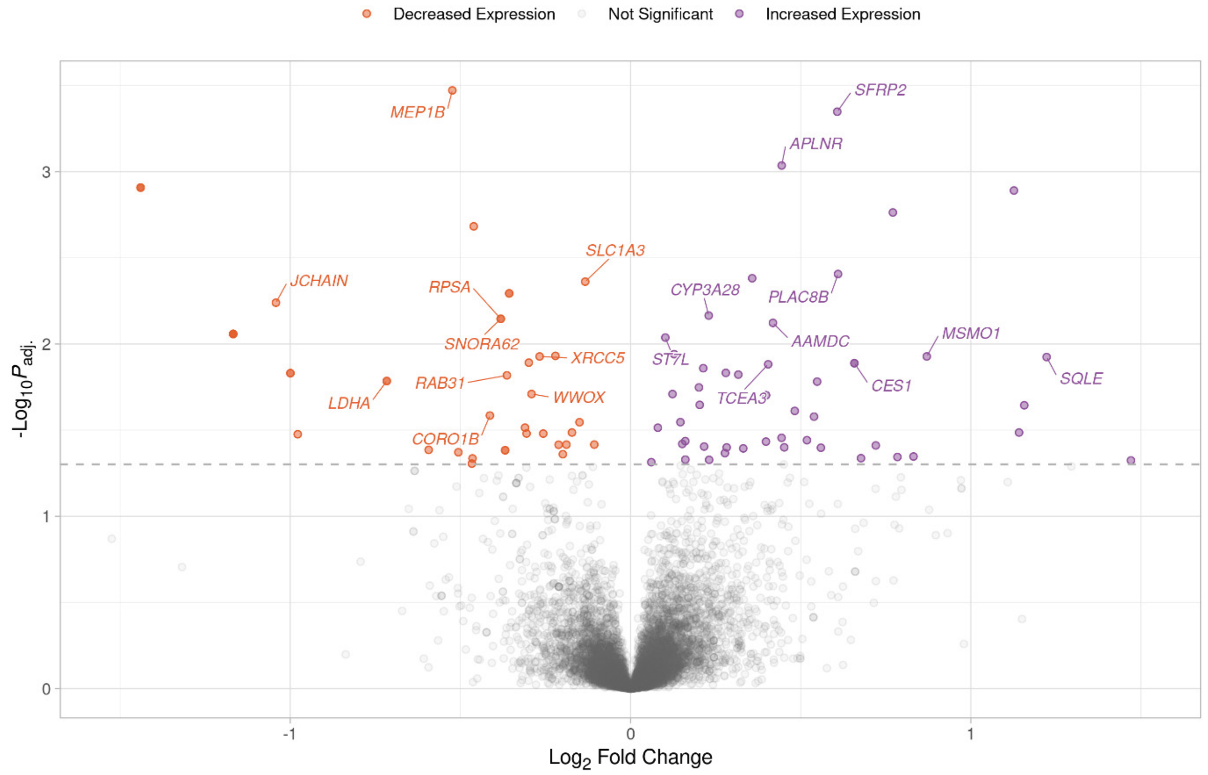


**Fig S22.** Volcano plot showing the results of the RESP contrast for the liver samples at 32 days post infection (dpi). Each data point represents a gene with the position on the *x*- and *y*-axes indicating the $log_2$ fold change and $-log_{10}P_{adj.}$, respectively. Genes above the horizontal dashed line are significantly differentially expressed with the colours representing the change in expression. The top 10 most significant genes for increased and decreased expression with gene symbols are labelled.

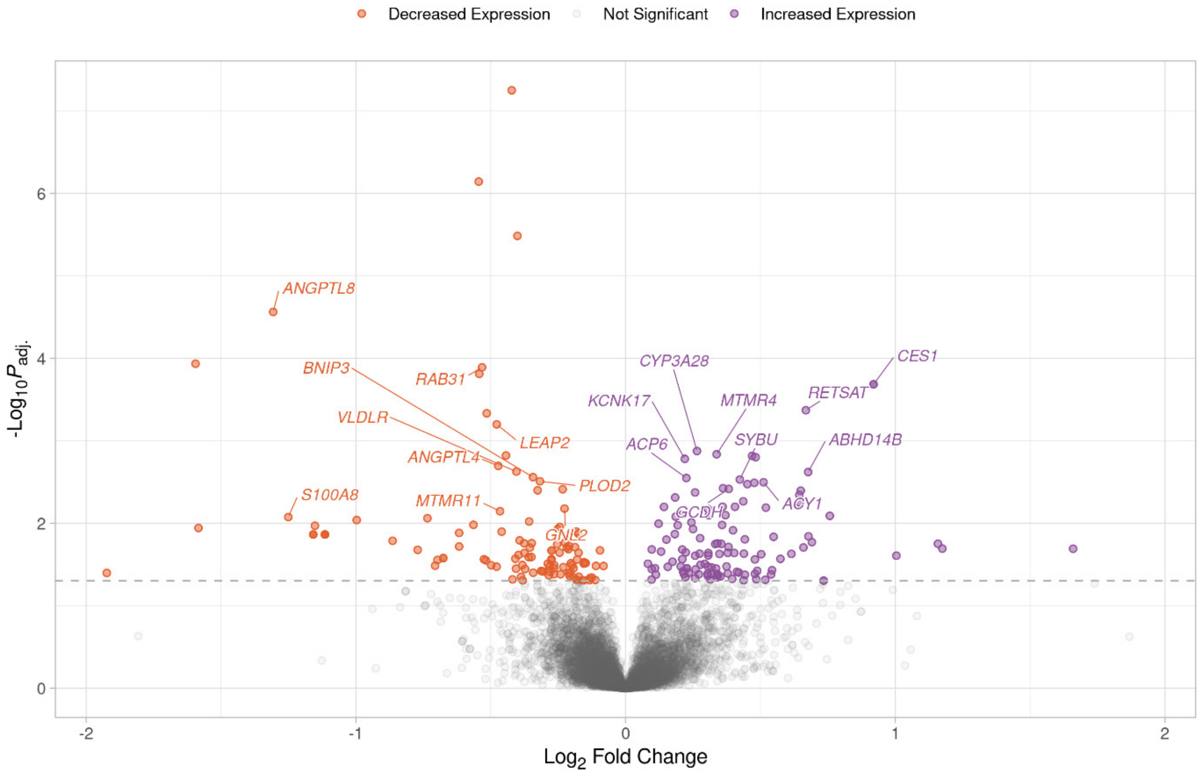


**Fig S23.** Volcano plot showing the results of the RESP contrast for the liver samples at 35 days post infection (dpi). Each data point represents a gene with the position on the *x*- and *y*-axes indicating the $\log_2$ fold change and $-\log_{10}P_{adj.}$, respectively. Genes above the horizontal dashed line are significantly differentially expressed with the colours representing the change in expression. The top 10 most significant genes for increased and decreased expression with gene symbols are labelled.

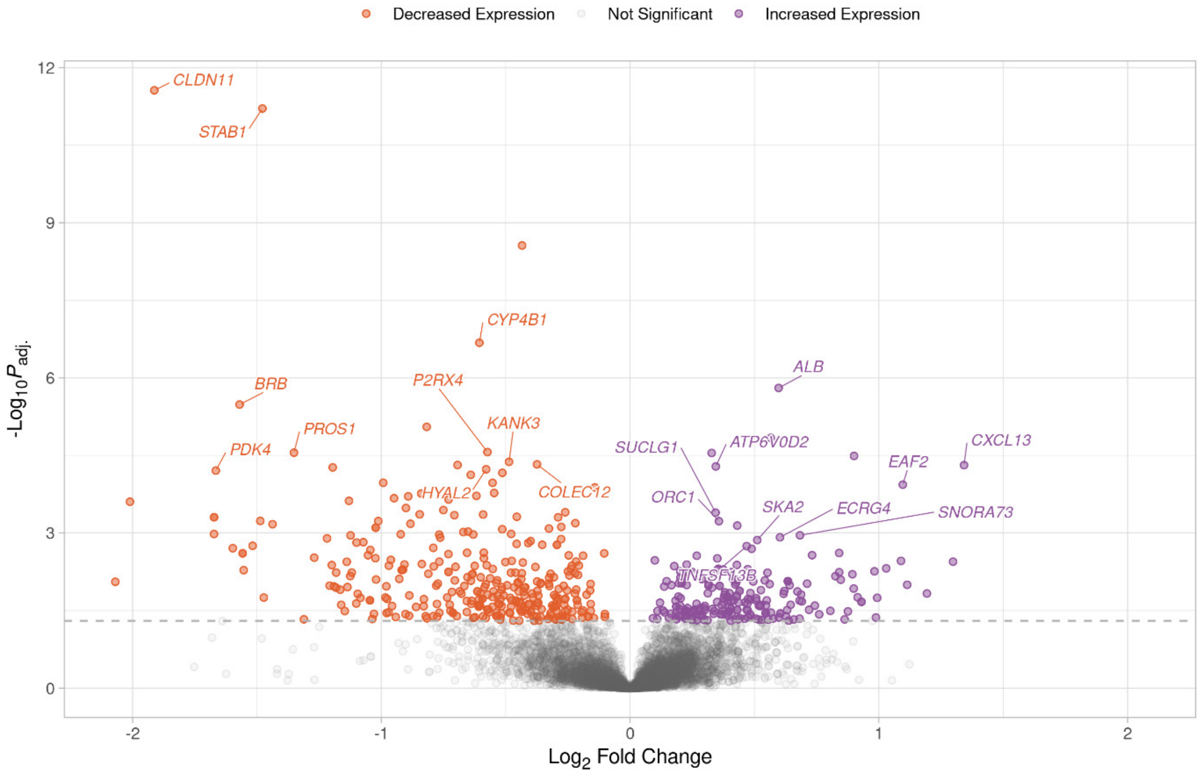


**Fig S24.** Volcano plot showing the results of the RESP contrast for the lymph node samples at 21 days post infection (dpi). Each data point represents a gene with the position on the *x*- and *y*-axes indicating the $\log_2$ fold change and $-\log_{10}P_{adj.}$, respectively. Genes above the horizontal dashed line are significantly differentially expressed with the colours representing the change in expression. The top 10 most significant genes for increased and decreased expression with gene symbols are labelled.

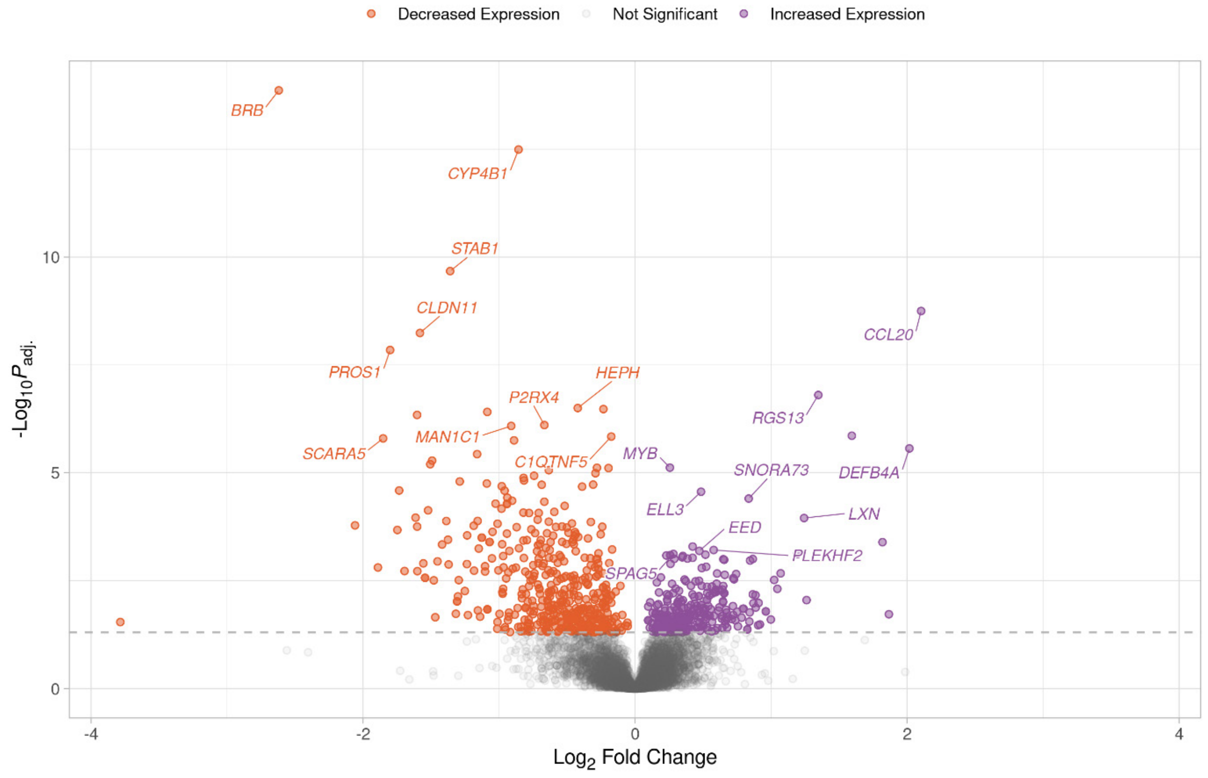


**Fig S25.** Volcano plot showing the results of the RESP contrast for the lymph node samples at 35 days post infection (dpi). Each data point represents a gene with the position on the *x*- and *y*-axes indicating the $\log_2$ fold change and $-\log_{10}P_{adj.}$, respectively. Genes above the horizontal dashed line are significantly differentially expressed with the colours representing the change in expression. The top 10 most significant genes for increased and decreased expression with gene symbols are labelled.

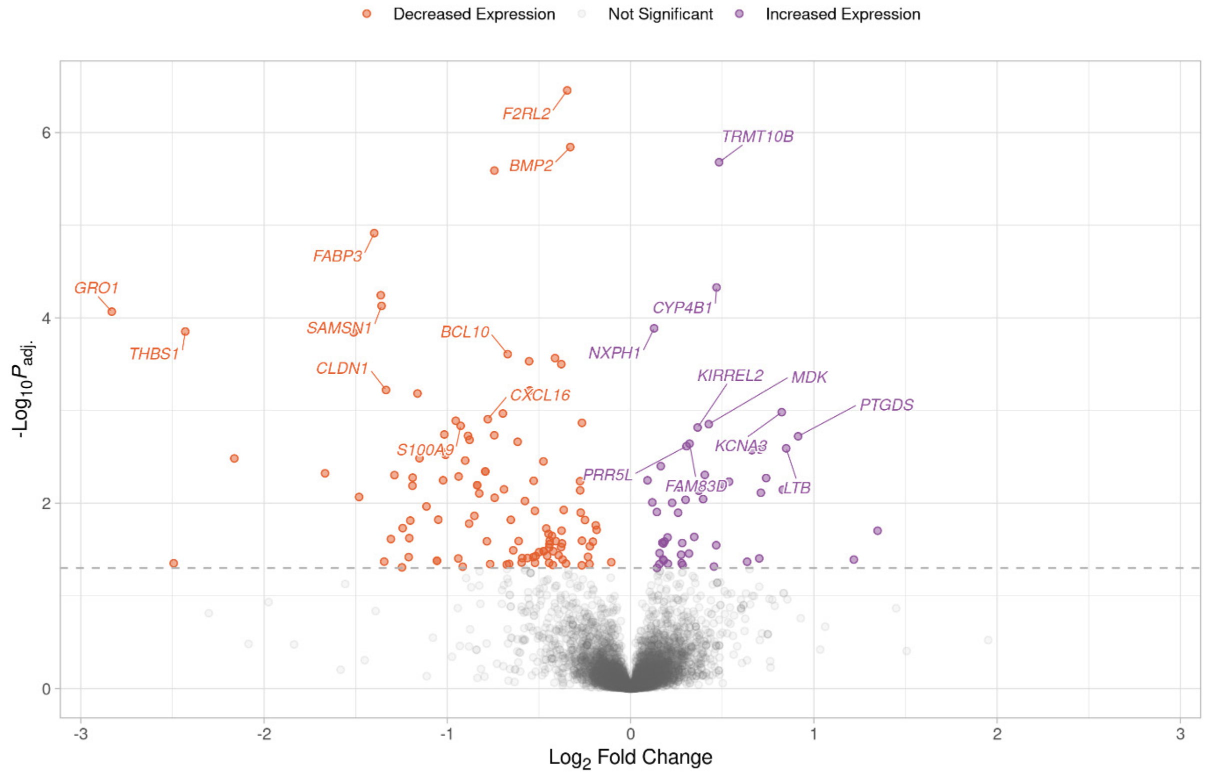


**Fig S26.** Volcano plot showing the results of the RESP contrast for the spleen samples at 21 days post infection (dpi). Each data point represents a gene with the position on the *x*- and *y*-axes indicating the $\log_2$ fold change and $-\log_{10}P_{adj.}$, respectively. Genes above the horizontal dashed line are significantly differentially expressed with the colours representing the change in expression. The top 10 most significant genes for increased and decreased expression with gene symbols are labelled.

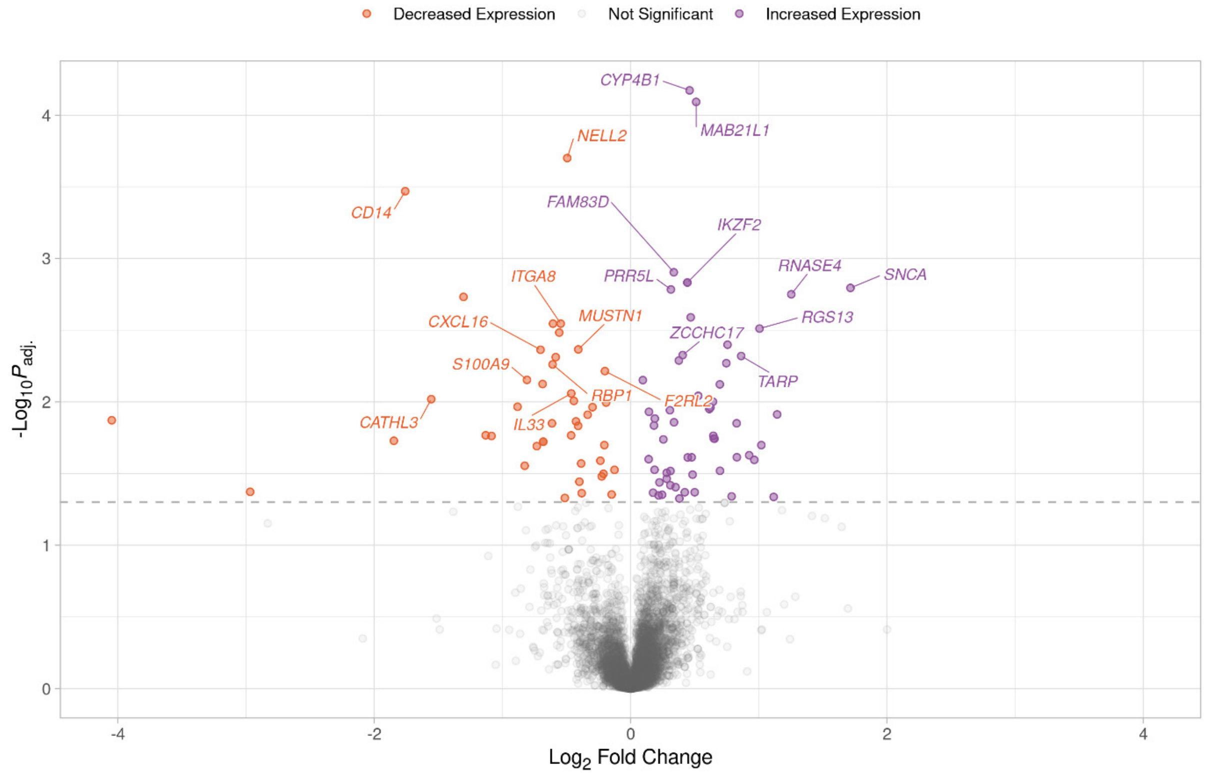


**Fig S27.** Volcano plot showing the results of the RESP contrast for the spleen samples at 35 days post infection (dpi). Each data point represents a gene with the position on the *x*- and *y*-axes indicating the $\log_2$ fold change and $-\log_{10}P_{adj.}$, respectively. Genes above the horizontal dashed line are significantly differentially expressed with the colours representing the change in expression. The top 10 most significant genes for increased and decreased expression with gene symbols are labelled.

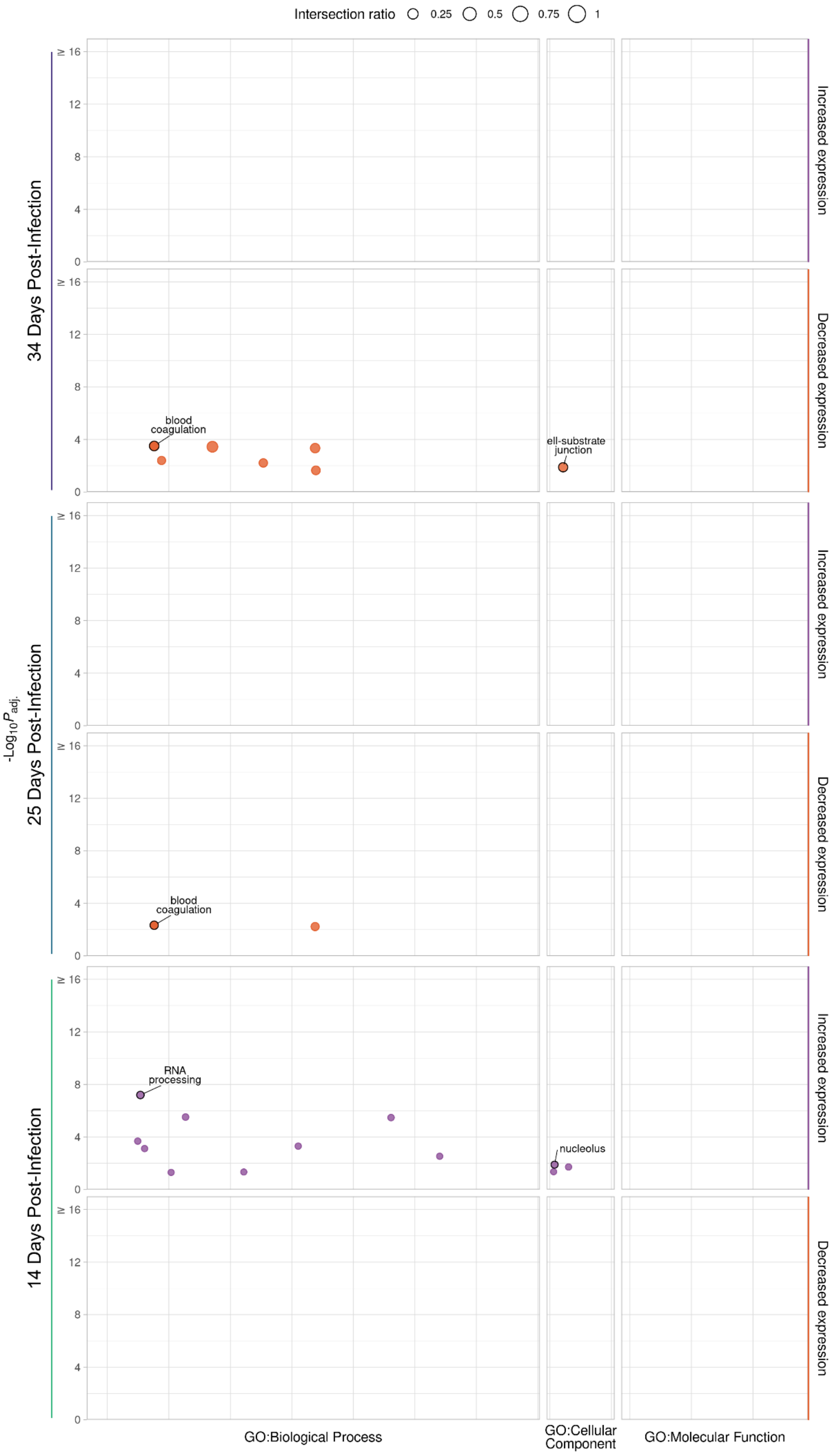


**Fig S28.** g:Profiler functional enrichment of significantly differentially expressed genes (DEGs) in the peripheral blood mononuclear cell (PBMC) sample RESP contrasts. Each circle represents a significantly enriched GO term with the size indicating the ratio of the intersection between the term and the DEGs. The vertical axis shows the $-\log_{10}P_{adj.}$ value and the vertical panels and colours indicate the direction of change in expression. The horizontal panels indicate the source of the term and the position within the panels groups terms from the same

GO subtree. The top driver GO terms (up to a maximum of 10) are indicated with a black outline and label.

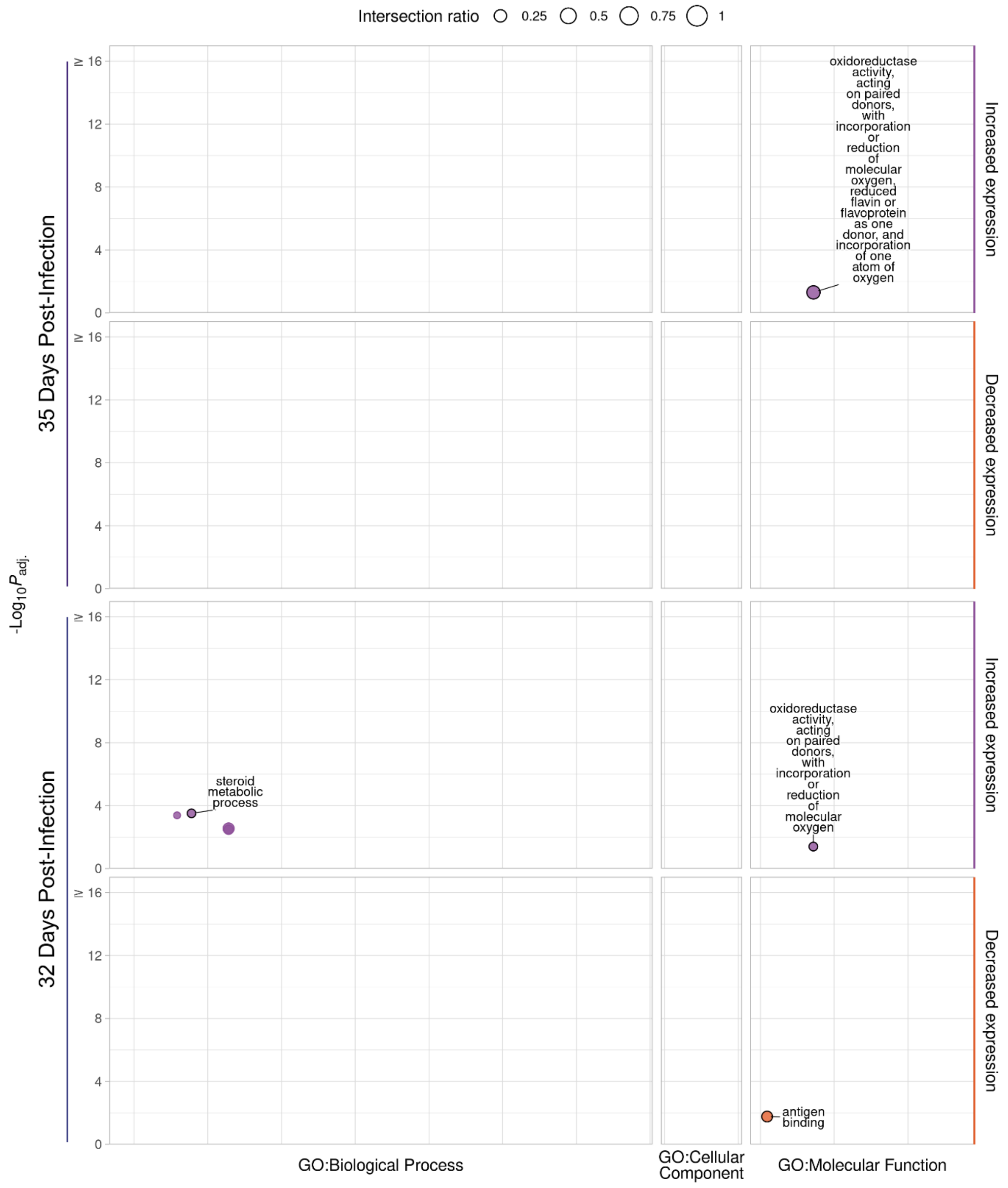


**Fig S29.** g:Profiler functional enrichment of significantly differentially expressed genes (DEGs) in the liver sample RESP contrasts. Each circle represents a significantly enriched GO term with the size indicating the ratio of the intersection between the term and the DEGs. The vertical axis shows the -log$_{10}P_{adj.}$ value and the vertical panels and colours indicate the direction of change in expression. The horizontal panels indicate the source of the term and the position within the panels groups terms from the same GO subtree. The top driver GO terms (up to a maximum of 10) are indicated with a black outline and label.

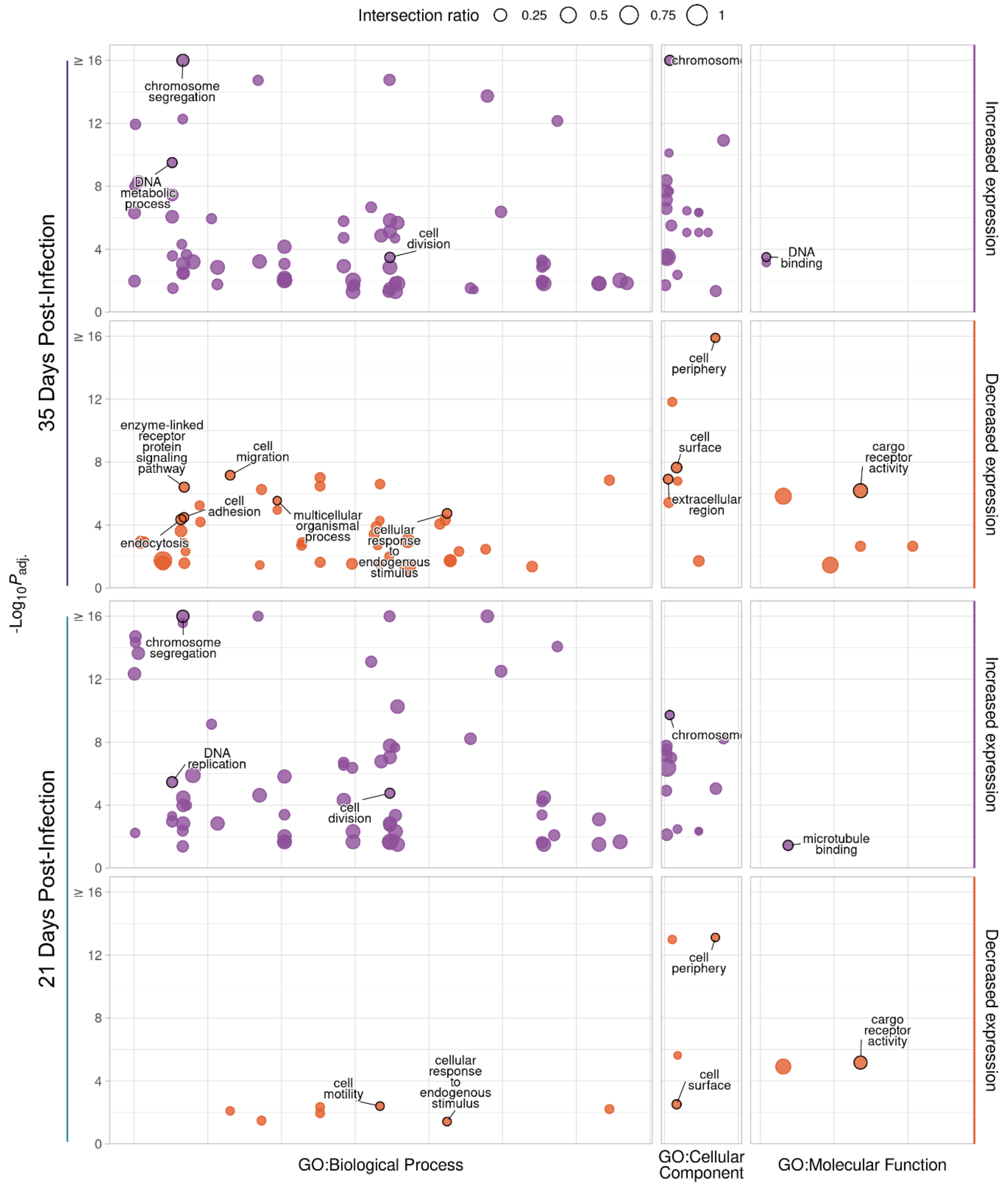


**Fig S30.** g:Profiler functional enrichment of significantly differentially expressed genes (DEGs) in the lymph node sample RESP contrasts. Each circle represents a significantly enriched GO term with the size indicating the ratio of the intersection between the term and the DEGs. The vertical axis shows the $-\log_{10}P_{adj.}$ value and the vertical panels and colours indicate the direction of change in expression. The horizontal panels indicate the source of the term and the position within the panels groups terms from the same GO subtree. The top driver GO terms (up to a maximum of 10) are indicated with a black outline and label.

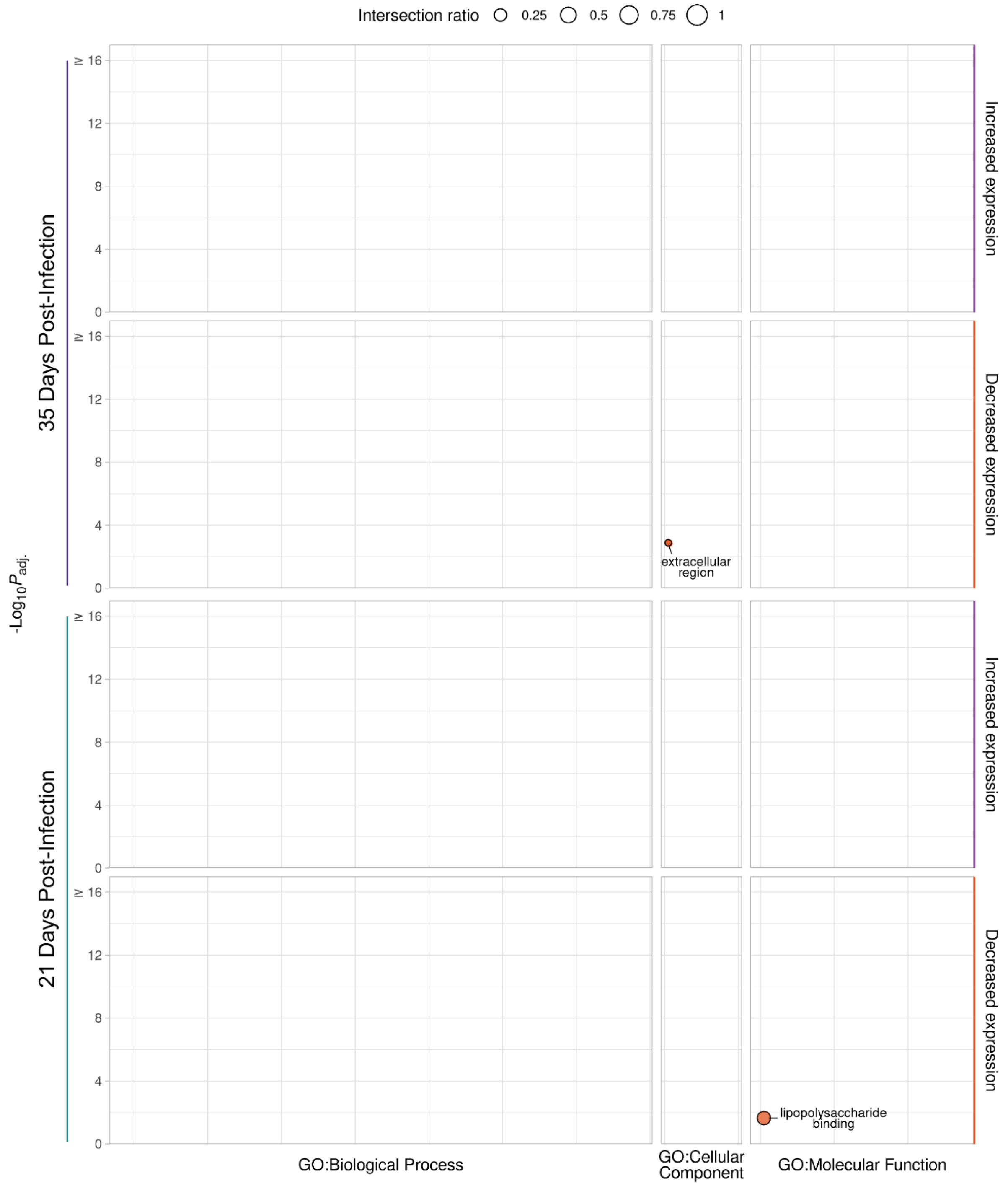


**Fig S31.** g:Profiler functional enrichment of significantly differentially expressed genes (DEGs) in the spleen sample RESP contrasts. Each circle represents a significantly enriched GO term with the size indicating the ratio of the intersection between the term and the DEGs. The vertical axis shows the -$\log_{10}P_{adj.}$ value and the vertical panels and colours indicate the direction of change in expression. The horizontal panels indicate the source of the term and the position within the panels groups terms from the same GO subtree. The top driver GO terms (up to a maximum of 10) are indicated with a black outline and label.

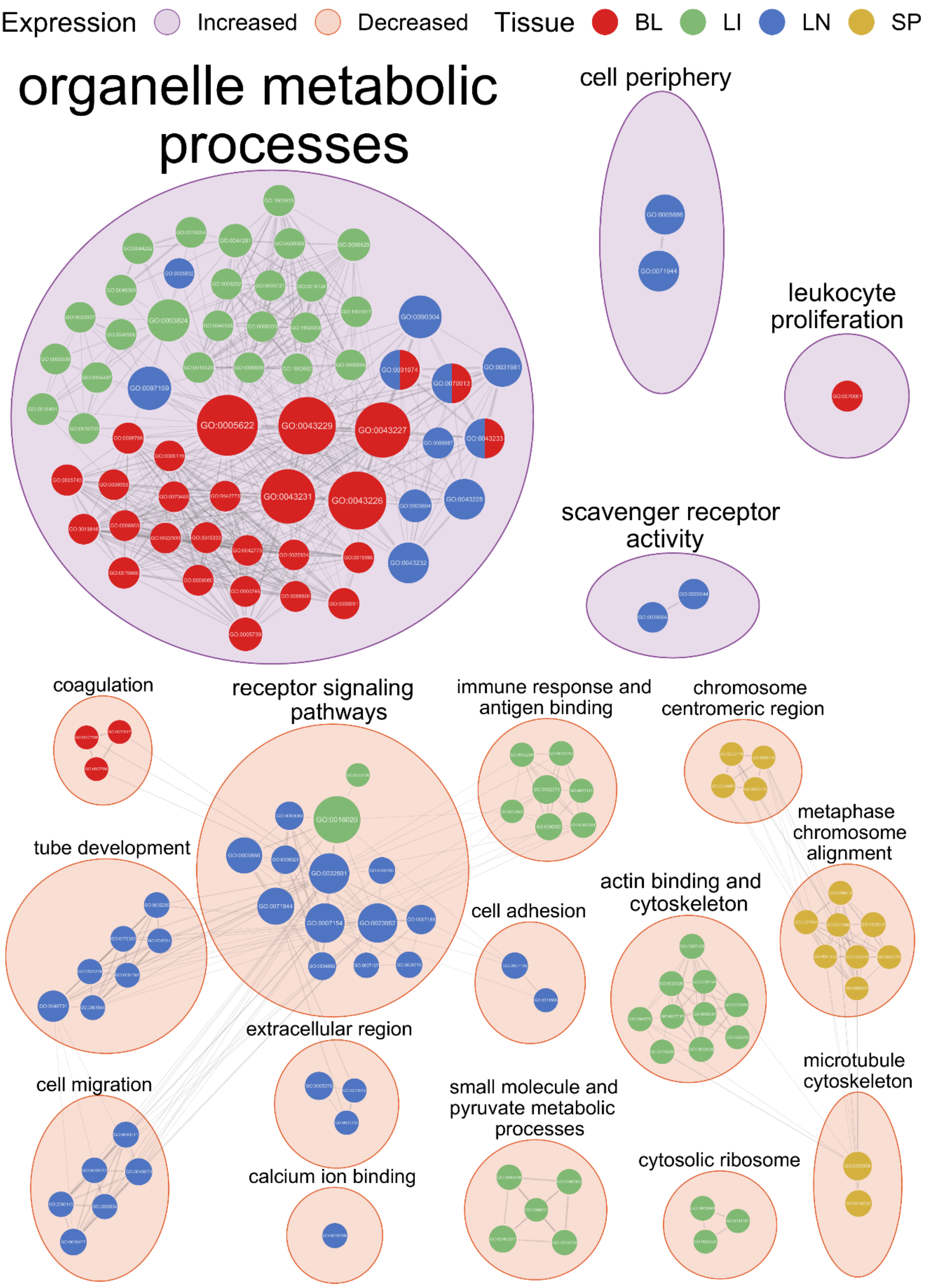


**Fig S32.** EnrichmentMap network of significantly enriched GO terms identified from g:Profiler functional enrichment of significant differentially expressed genes (DEGs) for the DIRE contrasts.Each node represents a GO term with the colour of the node representing the tissue and the size representing the number of genes in the GO term. The edges indicate overlap between the GO terms with the width of the edges representing the similarity coefficient for the connected GO terms. The GO terms are clustered by AutoAnotate with the background colour of the clusters representing the direction of expression. The clusters are labelled with the size of the label scaling with the number of GO terms in the cluster.

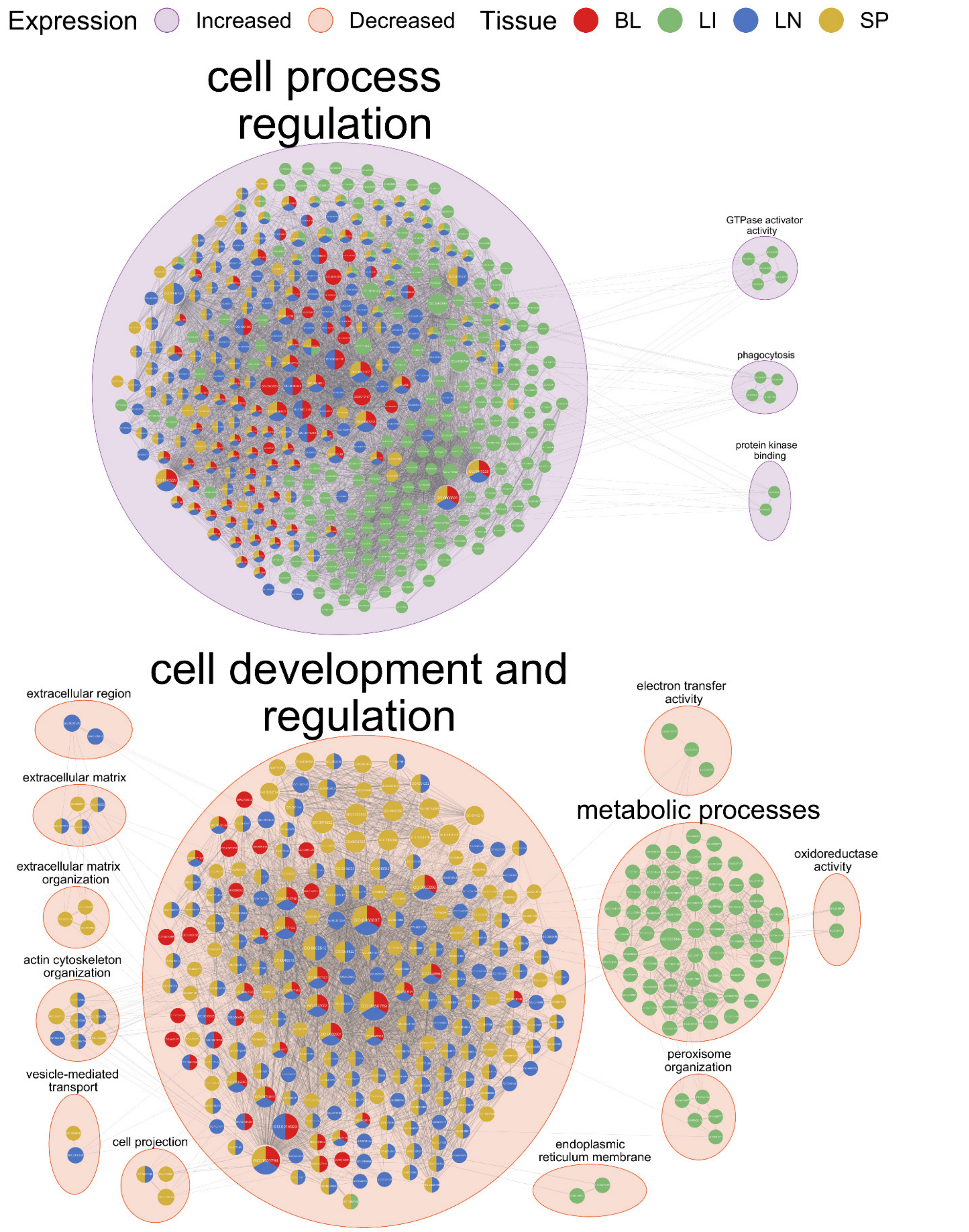


**Fig S33.** EnrichmentMap network of significantly enriched GO terms identified from g:Profiler functional enrichment of significant differentially expressed genes (DEGs) for the NDAM contrasts. Each node represents a GO term with the colour of the node representing the tissue and the size representing the number of genes in the GO term. The edges indicate overlap between the GO terms with the width of the edges representing the similarity coefficient for the connected GO terms. The GO terms are clustered by AutoAnotate with the background colour of the clusters representing the direction of expression. The clusters are labelled with the size of the label scaling with the number of GO terms in the cluster.

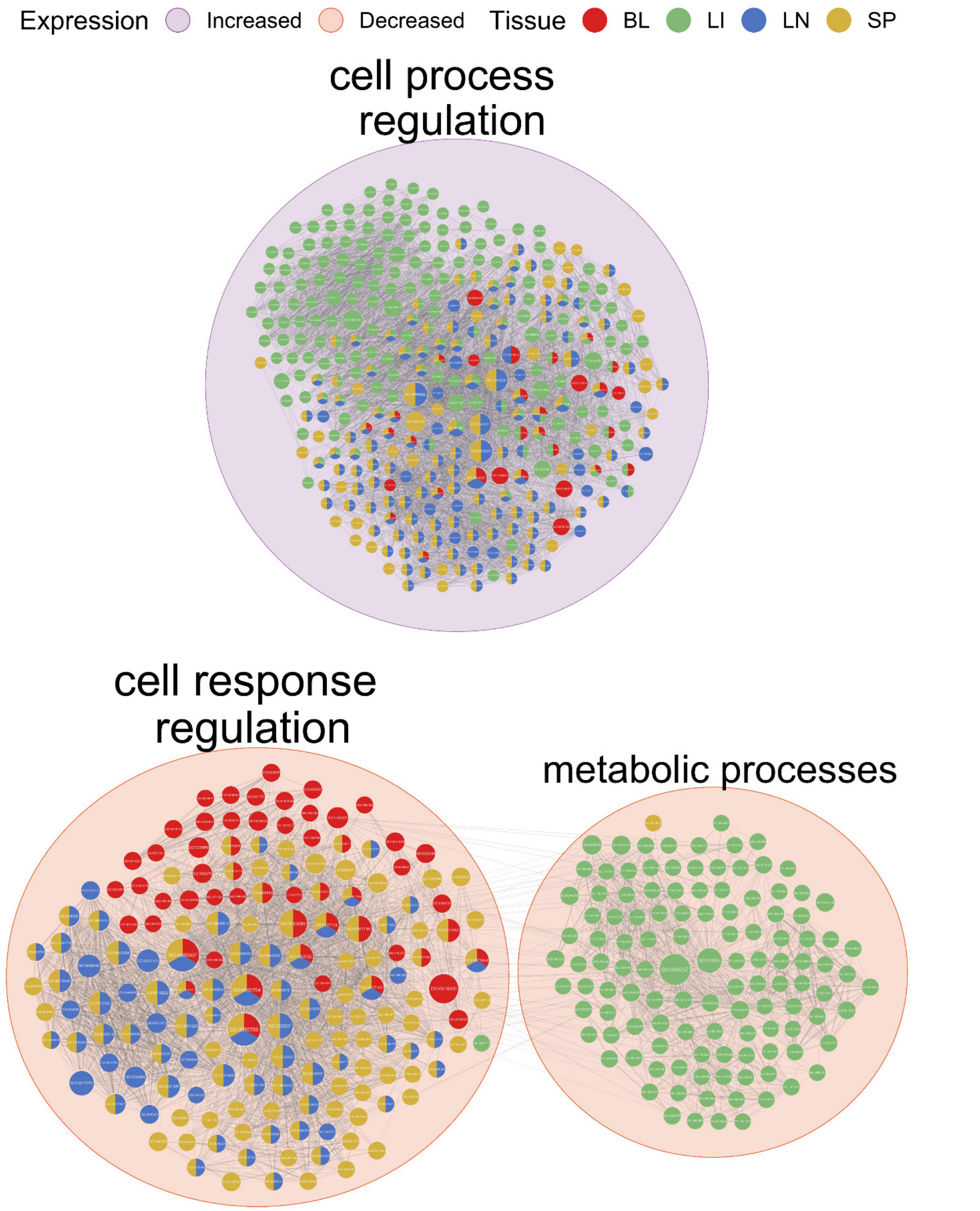


**Fig S34.** EnrichmentMap network of significantly enriched GO terms identified from g:Profiler functional enrichment of significant differentially expressed genes (DEGs) for the BORA contrasts. Each node represents a GO term with the colour of the node representing the tissue and the size representing the number of genes in the GO term. The edges indicate overlap between the GO terms with the width of the edges representing the similarity coefficient for the connected GO terms. The GO terms are clustered by AutoAnotate with the background colour of the clusters representing the direction of expression. The clusters are labelled with the size of the label scaling with the number of GO terms in the cluster.
